# Scalable nonparametric clustering with unified marker gene selection for single-cell RNA-seq data

**DOI:** 10.1101/2024.02.11.579839

**Authors:** Chibuikem Nwizu, Madeline Hughes, Michelle L. Ramseier, Andrew W. Navia, Alex K. Shalek, Nicolo Fusi, Srivatsan Raghavan, Peter S. Winter, Ava P. Amini, Lorin Crawford

**Affiliations:** Center for Computational Molecular Biology, Brown University, Providence, RI, USA; Warren Alpert Medical School of Brown University, Providence, RI, USA; Microsoft Research, Cambridge, MA, USA; Broad Institute of MIT and Harvard, Cambridge, MA, USA; Institute for Medical Engineering and Science, Massachusetts Institute of Technology, Cambridge, MA, USA; Koch Institute for Integrative Cancer Research, Massachusetts Institute of Technology, Cambridge, MA, USA; Department of Chemistry, Massachusetts Institute of Technology, Cambridge, MA, USA; Ragon Institute of MGH, MIT, and Harvard, Cambridge, MA, USA; Department of Medical Oncology, Dana-Farber Cancer Institute, Boston, MA, USA; Harvard Medical School, Boston, MA, USA; Department of Medicine, Brigham and Women’s Hospital, Boston, MA, USA

## Abstract

Clustering is commonly used in single-cell RNA-sequencing (scRNA-seq) pipelines to characterize cellular heterogeneity. However, current methods face two main limitations. First, they require user-specified heuristics which add time and complexity to bioinformatic workflows; second, they rely on post-selective differential expression analyses to identify marker genes driving cluster differences, which has been shown to be subject to inflated false discovery rates. We address these challenges by introducing nonparametric clustering of single-cell populations (NCLUSION): an infinite mixture model that leverages Bayesian sparse priors to identify marker genes while simultaneously performing clustering on single-cell expression data. NCLUSION uses a scalable variational inference algorithm to perform these analyses on datasets with up to millions of cells. Through simulations and analyses of publicly available scRNA-seq studies, we demonstrate that NCLUSION (i) matches the performance of other state-of-the-art clustering techniques with significantly reduced runtime and (ii) provides statistically robust and biologically relevant transcriptomic signatures for each of the clusters it identifies. Overall, NCLUSION represents a reliable hypothesis-generating tool for understanding patterns of expression variation present in single-cell populations.

## Introduction

Recent advances in sequencing technologies have increased the throughput of genomic studies to millions of single cells, necessitating computational workflows to explore and analyze these data ^1^. In single-cell RNA sequencing (scRNA-seq), unsupervised clustering and marker gene selection are integral steps in the exploratory phase of analyses ^2–5^. Clustering facilitates the identification of cell types, while marker gene selection enables the annotation of gene modules and cluster-specific biological programs. However, there has yet to be a consensus on the best approach to clustering cells and identifying the transcriptomic signatures that characterize them ^6,7^. This has resulted in a multitude of proposed clustering methods for single-cell data, many of which are reliant on various user-defined heuristics that prevent practitioners from performing an unbiased survey of data and limit each method’s “out-of-the-box” applicability when analyzing multiple studies.

Many current clustering approaches take a subset of highly variable genes as input, use dimensionality reduction techniques to simplify the representation of single-cell expression for these genes, and then perform clustering on top of this reduced representation. K-nearest neighbor (KNN) algorithms ^8^, for example, generate nearest-neighbor (NN) graphs using transcriptomic similarity scores between cells and then perform Louvain clustering on the estimated graphs. Popular methods such as Seurat ^9^ and scLCA ^10^ use principal component analysis (PCA) and singular value decomposition to learn a lower-dimensional representation of cells, respectively. Ensemble approaches such as scCCESS-SIMLR ^11^ learn over a mixture of kernels to generate a final cell-cell similarity matrix which is then used in a spectral clustering algorithm.

Notably, selecting an appropriate embedding for single-cell data is not always a straightforward task. Previous studies have shown that if the generated lower-dimensional representation does not accurately capture underlying biological relationships between cells, then both the quality of clustering and the generalizability of findings in downstream analyses can be compromised ^12,13^. Factors such as the number of highly variable genes retained during data preprocessing and the number of components used to define the lower-dimensional embedding can affect the ability of a clustering algorithm to identify fine-grained differences between cell types ^7,14^. Furthermore, nearly all state-of-the-art clustering methods require users to specify the number of clusters *K* to be used in the algorithm. Strategies such as consensus-finding ^11,15^, outlier detection ^16^, and iterative cluster merging and splitting ^17–19^ rely on human-in-the-loop interactive steps within their algorithms to determine an “optimal” choice for *K*. Generally, requiring users to make these additional decisions can add significant time and complexity when using clustering as a preliminary analysis in bioinformatic workflows.

Perhaps the biggest limitation of current single-cell clustering algorithms is that most do not directly identify top marker genes that are driving the inference of different biologically significant clusters; instead, they use post-selective inference to find genes that are differentially expressed between the inferred cell groups ^7,20^. Since the point of clustering algorithms is to separate dissimilar data into different groups, it is expected *a priori* that there are differences between the groups and any test statistics computed by comparing the groups are likely to be inflated due to “data double dipping”. Many studies have shown that performing this post-selective inference uncorrected can lead to inflated type I error rates ^20,21^. Though there has been work developed to correct for post-selection, these are still in nascency, and most do not yet scale to high-dimensional settings ^22–27^. Recently, others have proposed unified frameworks for simultaneous clustering and marker gene selection using hierarchical tree-based algorithms ^28^, regularized copula models ^29^, and *post hoc* sensitivity measures ^30^. However, these approaches rely on arbitrary thresholding to find “significant” marker genes and fail to theoretically test a well-defined null hypothesis, making them difficult to biologically interpret.

We present “Nonparametric CLustering of SIngle-cell populatiONs” (NCLUSION): a unified Bayesian nonparametric framework that simultaneously performs clustering and marker gene selection. NCLUSION works directly on normalized single-cell count data, bypassing the need to perform dimensionality reduction. By modeling the expression of each gene as a sparse hierarchical Dirichlet process normal mixture model ^31–37^, NCLUSION both learns the optimal number of clusters based on the variation observed between cellular expression profiles and uses sparse prior distributions to identify genes that significantly influence cluster definitions. The key to our proposed integrative framework is that clustering and extracting marker genes concurrently is a more efficient approach to the exploratory analysis of scRNA-seq data, as it effectively allows each process to inform the other. Most importantly, our approach eliminates the need for human-in-the-loop decisions, significantly reducing the runtime and complexity of these analyses. Altogether, NCLUSION mitigates the need for heuristic choices (e.g., choosing specific lower-dimensional embeddings), avoids iterative hyper-parameter optimization, bridges the interpretability gap suffered by many unsupervised learning algorithms in single-cell applications, and scales to accommodate the growing sizes of emerging scRNA-seq datasets.

## Results

### NCLUSION simplifies traditional clustering workflows

Conventional scRNA-seq clustering approaches include numerous steps that require user heuristics or human-in-the-loop decisions which increase runtime and complexity (Fig 1A). These can range from deciding how to optimally embed high-dimensional expression data into a lower-dimensional space to selecting the number of clusters, *K*, to identify in the data. Furthermore, current methods require that marker gene selection is performed post-clustering, which can lead to inflated rates of false discovery ^20,21^. In this work, we aim to address these challenges using a new approach: NCLUSION.

**Fig 1.**
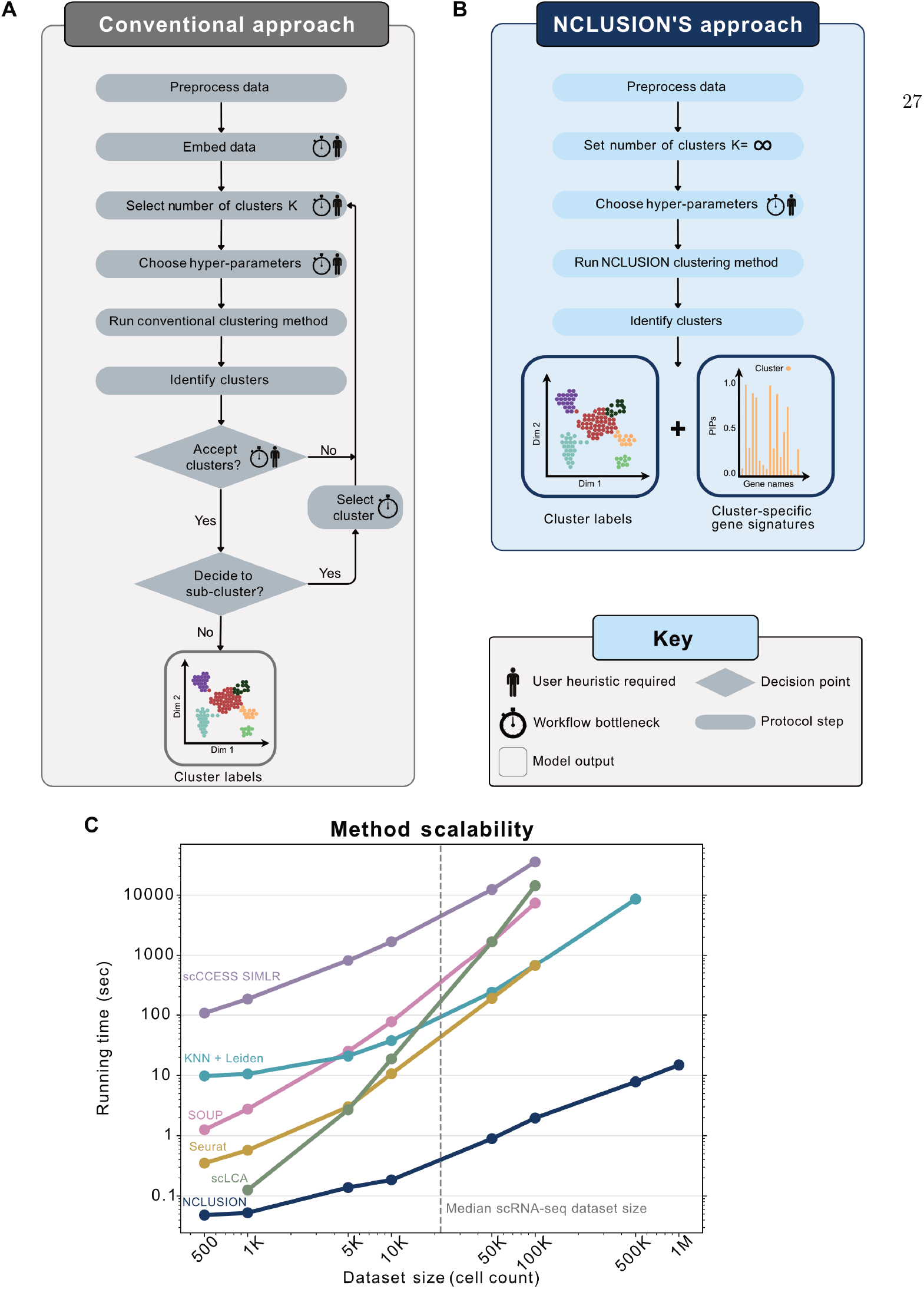
NCLUSION provides a scalable, unified workflow for both clustering and marker gene selection in single-cell analysis. **(A)** Conventional clustering algorithms require user heuristics and decision making steps that increase wall clock runtime (e.g., selection and human-in-the-loop refinement of the number of clusters *K*). **(B)** The nonparameteric workflow of NCLUSION reduces the number of choices and heuristics that users have to make while also performing cluster-specific variable selection to identify top marker genes for downstream investigation. **(C)** Runtimes of NCLUSION and other baselines on the BRAIN-LARGE dataset with a fixed set of 720 genes and an increasing sample size ranging from *N* = 500 to 1 million cells.

NCLUSION leverages a Bayesian nonparametric mixture modeling framework to reduce the number of choices that users need to make while simultaneously performing variable selection to identify top cluster-specific marker genes for downstream analyses (Fig 1B; see Methods for details). There are three key components of our model formulation that distinguish it from traditional bioinformatic workflows. First, NCLUSION is fit directly on single-cell expression matrices and does not require the data to be embedded within a lower-dimensional space. Second, we implicitly assume *a priori* that cells can belong to one of infinitely many different clusters. This is captured by placing a Dirichlet process prior over the cluster assignment probabilities for each cell. By allowing the number of possible clusters *K* = *∞*, we remove the need for users to iterate through different values until they find the optimal choice. Third, NCLUSION assumes that not all genes are important when assigning cells to a given cluster. To model this, we place a spike and slab prior on the mean expression of each gene within each cluster. This prior shrinks the mean of genes that play an insignificant role in cluster formation towards zero.

To identify cluster-specific marker genes, we start by estimating posterior inclusion probabilities (PIPs), which represent our confidence that a gene’s mean expression within a cluster is nonzero. These PIPs act as a signature that can be used to distinguish clusters. Since NCLUSION fits each gene and cell independently, signatures learned between clusters can share the same subsets of genes. To select for unique cell type markers, we multiply each PIP with a usage weight, which is calculated by performing min-max normalization over the proportion of clusters in which a gene is determined to be statistically significant (i.e., using the median probability model threshold ^38^ PIP ≥ 0.5). We use these adjusted inclusion probabilities with the effect size sign (ESS) and strictly standardized mean difference (SSMD) of each gene to filter for significant genes that are substantially up-regulated (indicated by positive ESS and large SSMD values). The genes remaining after filtering make up cluster-specific marker gene modules that provide insight into the biological features underlying cluster assignments.

To enable efficient posterior inference that can scale as the size of scRNA-seq datasets continues to grow, we train NCLUSION using variational expectation-maximization (EM). This algorithm leverages a “mean-field” assumption to approximate the true posterior distribution over model parameter estimates with a product of simpler distributions ^39–41^. In training, our objective is to minimize the Kullback–Leibler (KL) divergence between the variational posterior and the true posterior ^42^. Optimization during model fitting occurs using a coordinate ascent procedure where parameters are sequentially updated based on their gradients (Methods and Supplementary Material). With this variational approach, NCLUSION is capable of scaling well up to 1 million cells without applying any dimensionality reduction to the input data.

We evaluated the runtime of NCLUSION against a set of state-of-the-art single-cell clustering methods using publicly available datasets. The methods we used for comparison include: Seurat ^9^, scLCA ^10^, K-nearest neighbors followed by the Leiden clustering algorithm (KNN+Leiden) ^43^, SOUP ^44^, and scCCESS-SIMLR ^11^. Each of these methods operates by first reducing the dimensionality of the input data and then performing clustering on the reduced representation. To the best of our knowledge, NCLUSION is the only method to date that clusters on the expression matrix directly while jointly identifying cluster-specific salient genes. NCLUSION does not incur additional runtime cost due to this model design choice. In fact, NCLUSION boasts a faster runtime compared to other methods, particularly as the number of cells in a dataset grows. We showcase this scalability on the BRAIN-LARGE ^45^ dataset (Fig 1C). To facilitate comparison across all baseline methods, we follow Lopez et al. ^14^ and limit the analysis to 720 genes, while subsampling the number of cells from 500 to 1 million cells. As a practical reference, we use a grey dotted line to highlight the runtime for each method at the median scRNA-seq dataset size as determined in 2020: 31,000 cells ^46^. We observed that only NCLUSION and KNN+Leiden were able to scale past 100,000 cells, while NCLUSION was the only method able to run on 1 million cells. Additionally, NCLUSION records competitive runtimes across varying numbers of genes (Fig S1). These scalability results show the potential of NCLUSION’s utility for emerging large-scale single-cell studies.

### NCLUSION accurately identifies clusters and marker genes in simulations

We used simulations to evaluate the performance of NCLUSION in controlled settings. Here, we used scDesign3^47^ to generate synthetic datasets consisting of 10,000 cells and 1,000 genes (due to computational constraints of the software) distributed over five clusters. We considered four different scenarios (with 20 replicates per scenario), where we varied cluster size and marker gene composition (Methods). Scenario I was the simplest, where we evenly distributed all cells across the five clusters, and each cluster had 50 marker genes. In Scenarios II and III, all five clusters had 50 marker genes, but one cluster had significantly fewer cells than the other four clusters. Lastly, in Scenario IV, each of the five clusters had the same number of cells, but one cluster had a signature of only 20 marker genes, while the other four clusters had 50 marker genes each.

We first compared the cluster identification accuracy of NCLUSION with the previously described algorithms: Seurat ^9^, scLCA^10^, K-nearest neighbors followed by the Leiden clustering algorithm (KNN+Leiden) ^43^, SOUP ^44^, and scCCESS-SIMLR ^11^. The relatively smaller size of these simulated datasets also allowed us to include three additional widely used methods: CIDR ^48^, SC3^49^, and scDeepCluster ^50^. We should highlight that all competing methods, except for NCLUSION, first perform dimensionality reduction prior to clustering, allowing us to evaluate the impact of dimensionality reduction on cluster recovery.

The normalized mutual information (NMI) and adjusted Rand index (ARI) were calculated to quantitatively evaluate the clustering results given by each algorithm. In all scenarios, scDeepCluster had the best performance, with NCLUSION, scLCA, and scCCESS-SIMLR rounding out the top four. The reason for scDeepCluster’s top performance is that it first performs Leiden clustering on principal components from the expression data to obtain the initial cluster assignments. Then, it uses a deep neural network to refine the cluster assignments; therefore, performance is highly dependent on the initialization of the cluster assignments and on the success of the initial clustering. NCLUSION, on the other hand, only performs clustering once to achieve comparable results. Notably, all methods performed relatively worse in Scenarios II and III than their performance in Scenarios I and IV (Fig S2; Table S1) due to the class imbalance in how the cells were distributed across clusters.

We also evaluated the performance of NCLUSION on marker gene detection using the same simulated datasets and compared it with three popular differential expression algorithms: DUBStepR ^51^, singleCellHaystack ^25^, and FESETM ^52^. Notably, of these methods, only NCLUSION is able to find cluster-specific marker genes. FESTEM uses an alternative algorithm (via the Scott-Knott test) to assign marker genes to clusters found by an independent clustering method, while both DUBStepR and singleCellHaystack aim to identify differentially expressed genes. To that end, we evaluated the global marker gene detection of each model using true positive rate (TPR), false discovery rate (FDR), and false positive rate (FPR; computed as 1-Specificity for each method). We found no statistically significant differences when comparing the median power of NCLUSION to the other approaches (Kruskal–Wallis *H* -test *P >* 0.99; Fig S3 and Table S2). However, importantly, NCLUSION and FESTEM were the only two methods to have high power while maintaining a low FDR and FPR. On the other hand, singleCellHaystack and DUBStepR achieved similar power but only because they incorrectly labeled many genes as being marker genes (resulting in markedly higher FDR and FPR).

### NCLUSION achieves competitive clustering performance on PBMC data with less runtime

Next, we assessed the quality of clustering done by NCLUSION as compared to other baseline approaches on real data. Here, we analyzed scRNA-seq from FACS-purified peripheral blood mononuclear cells (PBMCs) ^53^. This dataset captures 10 cellular populations, including CD14+ monocytes, CD34+ cells, major lymphoid lineages (B and T cells), as well as other phenotypic lineages within the T cell population, including CD4+ helper T cells, CD8+ cytotoxic T cells, CD4+ regulatory T cells, and CD4+ memory T cells. After quality control (Methods), the final dataset contained 94,615 cells and 5,000 genes with the highest standard deviation (post log-normalization) ^20,54^. We evaluated the performance of NCLUSION, Seurat, scLCA, the KNN+Leiden algorithm, SOUP, and scCCESS-SIMLR by comparing inferred cluster assignments to the cell type annotations from the original study, which were obtained via a combination of FACS analysis and clustering with Seurat ^53^ (Fig 3A-B).

**Fig 2.**
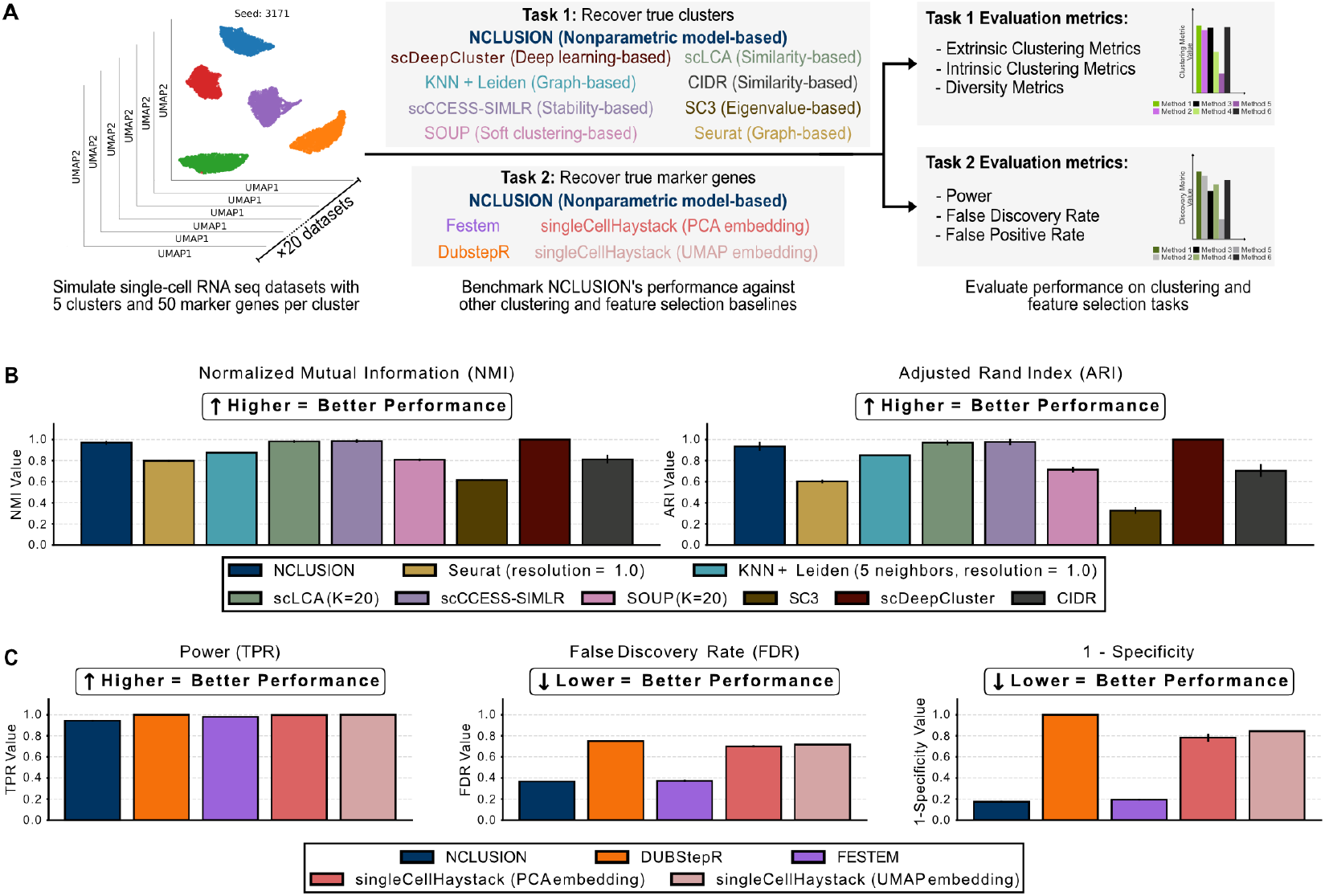
Comparing NCLUSION and competing algorithms on performing clustering and marker gene selection in a simulation study. Depicted are results for Scenario I where we evenly distributed all synthetically generated cells across five clusters and each cluster had a unique set of 50 marker genes. **(A)** Overview of the simulation framework used for evaluating the quality of clustering and marker gene selection for NCLUSION and each competing method. **(B)** Inferred cluster labels were compared to “true” annotations created during the simulation, where performance was measured according to (left) normalized mutual information (NMI) and (right) adjusted Rand index (ARI). **(C)** Assessment of marker gene selection was done on the global scale, where methods were evaluated on how well they could detect a “true” causal gene without taking cluster assignment into account. This was due to the limitation of competing methods not being able to identify cluster-specific genes. Evaluations were done by measuring the true positive rate (TPR; or power), false discovery rate (FDR), and false positive rate (FPR; computed as 1-Specificity) for each approach. Results for (B) and (C) are based on 20 simulations, with each bar plot representing the mean and the error bars covering a *±* 95% confidence interval.

**Fig 3.**
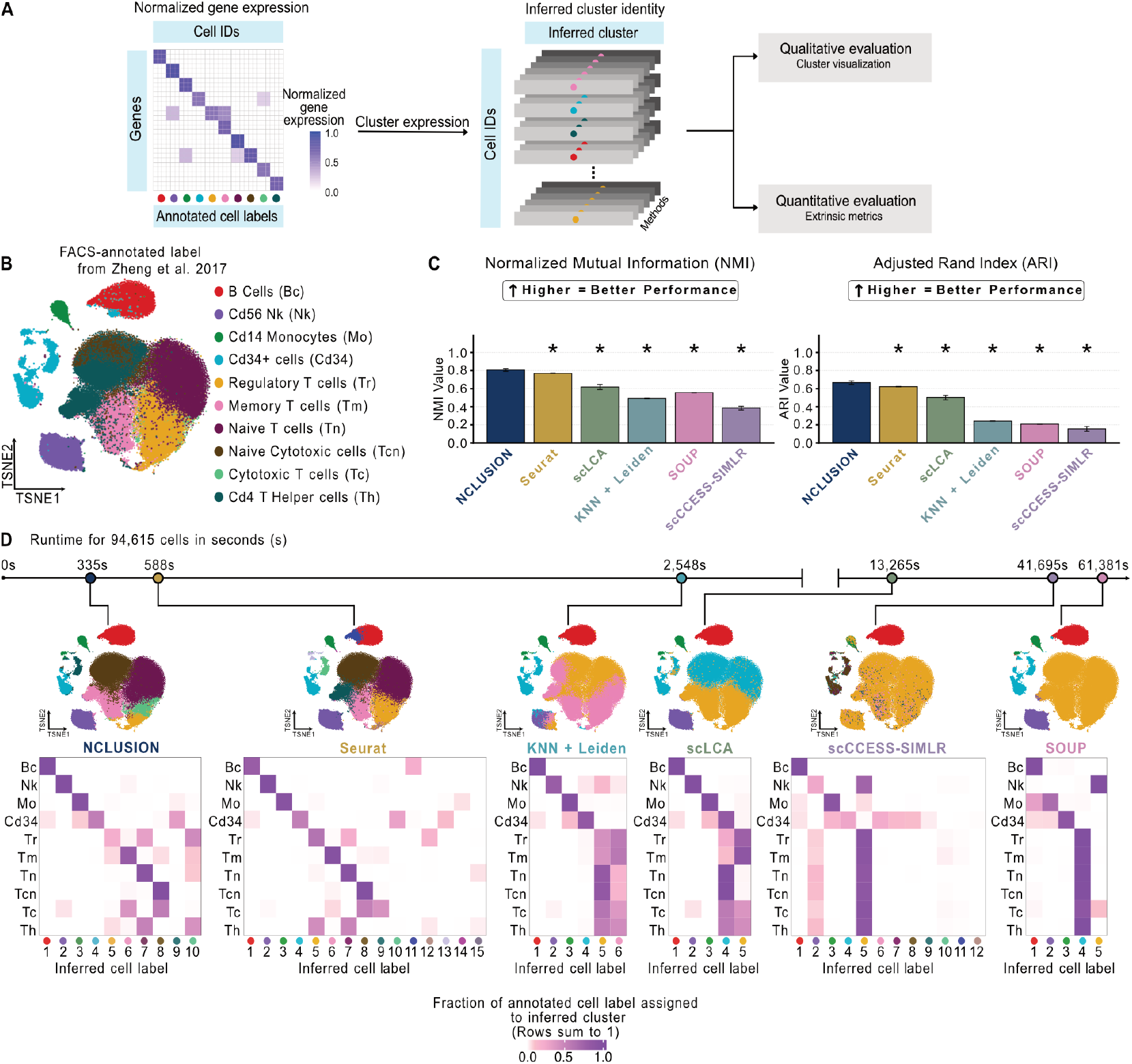
Clustering performance for NCLUSION and other baseline methods on the PBMC scRNA-seq dataset (*N* = 94,615 cells). **(A)** The framework used for evaluating the quality of clustering in each method. **(B)** Overview of FACS-based cell type annotations, visualized via t-distributed stochastic neighbor embedding (t-SNE), for the PBMC scRNA-seq dataset. These annotations serve as labels during the evaluation. **(C)** Assessment of the inferred cluster labels versus the experimental annotations, as quantified by two metrics: normalized mutual information (NMI) and adjusted Rand index (ARI) (for each method, we take five random 80% splits of the PBMC dataset; depicted in each bar plot is the mean *±*95% confidence interval). Asterisks indicate that there is a statistically significant difference in performance between NCLUSION and a corresponding method (two-sided t-test *P <* 0.05). **(D)** Visualizing the structure of the inferred clusters across all baselines using t-SNEs and a contingency heat map showing the prevalence of each cell type within each cluster. Methods are ordered from fastest (left) to slowest (right) in terms of runtime. The same lower dimensional representation of the data is reused with relabeling of the plots according to the results from each clustering algorithm.

To qualitatively assess clustering performance, we used contingency heat maps to evaluate how well each method captured the unique cell types across clusters (Fig 3C). For a given method, each *n*-th row of the heat map represents an annotation from the original study and each *k*-th column represents an inferred cluster identified by the method. The color saturation of each (*n, k*)-th element in the heat map indicates the fraction of a *n*-th cell annotation that a given method assigned to the *k*-th inferred cluster. Overall, all baselines were able to distinguish the B cell population from other cells, and each approach uniquely clustered a majority of the CD14+ monocytes together (Fig 3C). Furthermore, all methods except scCCESS-SIMLR and KNN+Leiden were able to separate the natural killer (NK) cell population with an inferred cluster occupancy rate of greater than 90% (Table S3). NCLUSION’s performance was most similar to that of Seurat (*χ*^2^-test *P* = 0.99 when assessing independence between their contingency tables). Both methods were able to identify major PBMC cell lineages and were the only approaches to divide the B cell, cytotoxic CD8+ T cells, and regulatory T cells into subpopulations ^55^ (Fig 3C).

As a quantitative assessment of each method’s clustering performance, we used the FACS-derived experimental annotations as reference labels and computed NMI and ARI, each measuring how well the clustering algorithm’s labels matched the reference labels (Methods). For both metrics, values closer to 1 indicate better clustering performance. To assess the robustness and consistency of each method, we ran them on five different randomly subsampled partitions containing 80% of the cells in the dataset. We report the mean metric score for each clustering algorithm across these partitions, along with corresponding 95% confidence intervals (Fig 3D). NCLUSION outperformed all competing approaches across both metrics. It obtained the highest mean NMI coefficient of 0.80 (*±*1.62 *×* 10^*−*2^ standard deviation), with Seurat and scLCA each scoring lower values of 0.77 (*±*2.78 *×* 10^*−*3^) and 0.62 (*±*3.04 *×* 10^*−*2^), respectively. These differences were statistically significant, as determined by two-sided t-tests (*P* = 1.08 *×* 10^*−*3^ and *P* = 1.90 *×* 10^*−*6^, respectively). When comparing performance using the ARI, NCLUSION remained competitive and significantly outperformed other methods (Table S4). Specifically, NCLUSION achieved a higher mean ARI of 0.67 (*±*2.02 *×* 10^*−*2^) when compared to 0.62 (*±*2.48 *×* 10^*−*3^; two-sided t-tests *P* = 1.17 *×* 10^*−*3^) for Seurat and 0.50 (*±*2.52 *×* 10^*−*2^; two-sided t-tests *P* = 3.10 *×* 10^*−*6^) for scLCA, respectively.

Lastly, NCLUSION recorded the shortest runtime for this analysis without any iterative processes or optimization of hyperparameters. It finished approximately 254 seconds (4.23 minutes) faster than Seurat, more than 12,900 seconds (215.00 mins or 3.58 hrs) faster than scLCA, and more than 41,331 seconds (688.85 mins or 11.48 hrs) faster than scCCESS-SIMLR.

### NCLUSION is well-powered to identify PBMC-specific marker genes

The key distinguishing property of NCLUSION is its inherent ability to perform variable selection. NCLUSION thus provides users with cluster labels for each cell as well as unique gene signatures that define each cluster. The statistical model underlying NCLUSION selects cluster-specific marker genes based on two criteria: (*i*) an adjusted posterior inclusion probability, PIP(*j*; *k*), which provides evidence that the *j*-th gene’s mean expression is uniquely nonzero within the *k*-th cluster, (*ii*) the sign of the *j*-th gene’s effect, ESS(*j*; *k*), which is used to determine whether it is uniquely up-regulated or down-regulated within the *k*-th cluster, and (*iii*) the magnitude of the *j*-th gene’s effect, SSMD(*j*; *k*), on the definition of the *k*-th cluster (Methods). We assessed the marker genes identified by NCLUSION for each of the inferred clusters in the PBMC dataset to determine whether they provide insights into the biology of different cell types (Fig 4A-B).

**Fig 4.**
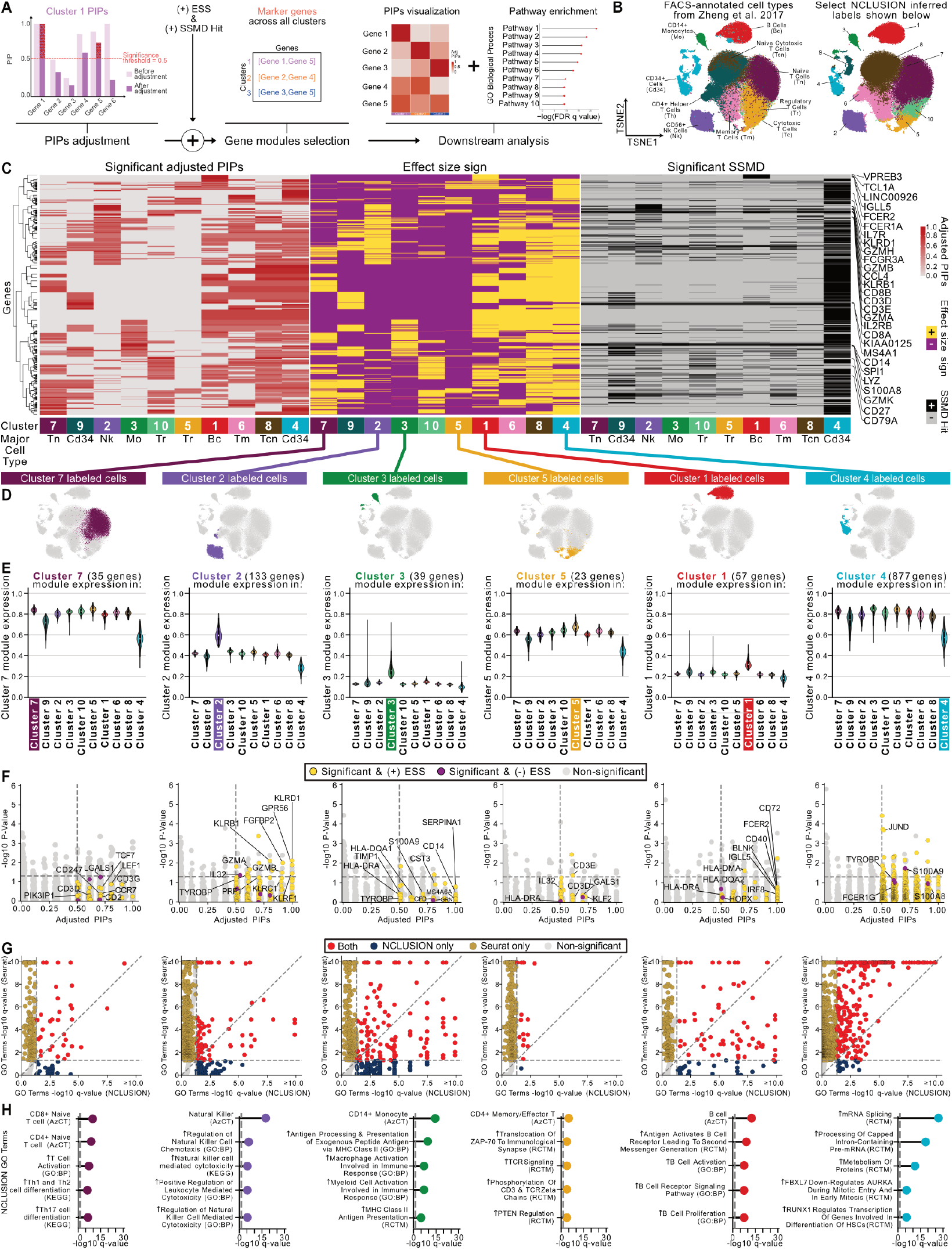
Evaluation of cluster-specific marker genes identified by NCLUSION on the PBMC dataset (*N* = 94,615 cells). **(A)** The framework used for assessing cluster-specific marker genes. (**B**) Embeddings of the experimental annotations for major cell types from the PBMC dataset compared to the clusters inferred by NCLUSION. **(C)** Heat maps of the adjusted posterior inclusion probabilities (PIPs) (left), effect size sign (ESS) (center), and strictly standardized mean difference (SSMD) (right) of significant genes in each cluster. Cluster-specific marker genes are selected as those that have a significant inclusion probability, are up-regulated in a given cluster, and have a large effect size magnitude such that PIP ≥ 0.5, ESS = +, and |SSMD(*j*; *k*) | ≥ *S*^*∗*^(*j*; *k*), respectively. Here *S*^*∗*^(*j*; *k*) is a threshold set to preserve a false positive rate of 0.05. **(D)** Highlighted location on t-SNEs of NCLUSION-inferred clusters that contain predominantly one cell type. **(E)** Violin plots comparing the normalized expression of cluster-specific marker genes in each of the inferred clusters. **(F)** Scatter plot comparing the marker genes identified using *post hoc* differential expression analysis with Seurat (yellow) versus the variable selection approach with NCLUSION (blue). Yellow points have PIP ≥ 0.5 and ESS = +, while purple points have PIP ≥ 0.5 and ESS = −, respectively. The vertical dashed line marks the median probability criterion ^38^, and the horizontal dashed line marks the Bonferroni-corrected threshold for significant *q*-values (i.e., an adjusted *P*). Genes in the top right quadrant are identified by both methods. **(G)** Scatter plot comparing gene ontology (GO) pathway enrichment analyses using cluster-specific marker genes from Seurat versus NCLUSION. The horizontal and vertical lines correspond to significant *q*-values being below 0.05. Pathways in the top right quadrant are selected by both approaches (red), while elements in the bottom right and top left quadrants are uniquely identified by NCLUSION (blue) and Seurat (orange), respectively. **(H)** Highlight of select top GO pathway enrichment analysis for the marker genes identified by NCLUSION. Plotted on the x-axis are the negative log-transformed *q*-values for each GO gene set. Gene sets with a *q*-value below 0.05 are deemed to be significant.

Overall, NCLUSION successfully identified cluster-specific marker genes that are known to be associated with examined cell types (Fig 4C and Table S5). For example, in cluster 8 which has cells mapping back to the cytotoxic and naive cytotoxic T cell population, we observed that NCLUSION correctly identified marker genes known to play an important role in cytotoxic T cell biology, such as *CD8A* (adjusted PIP = 0.90), *CD8B* (adjusted PIP = 0.70) ^56^, and *CD27* (adjusted PIP = 0.62) ^57^. Furthermore, genes associated with cytotoxicity, such as *GZMM*, tended to be selected in clusters largely containing CD8+ T cells (Cluster 8; adjusted PIP = 0.60) and NK cells (Cluster 2; adjusted PIP = 0.61)—two cell types that have been shown to have functionally similar cytotoxic activity ^58^. In other clusters, where we observed genes associated with both B cells (e.g., in Cluster 1, *CD19*, adjusted PIP = 1.00; *LINC00926*, adjusted PIP = 1.00; *MS4A1*, PIP = 0.90) ^59–61^ and myeloid lineages (e.g., in Cluster 3, *MS4A6A*, PIP = 1.00; *S100A8*, adjusted PIP = 0.93; *LYZ*, PIP = 0.90) ^62^, NCLUSION distinguished genes known to play an important role in T cell biology as statistically significant. This observation suggests that NCLUSION accounted for the variance among T cell and T cell-like expression patterns when distinguishing cell types in the PBMC dataset. We observed that our criteria for cluster-specific marker genes, based on high PIPs and positive ESS scores, strongly agreed with the relative over-expression of each gene in its respective cluster (Fig 4C). Imposing a threshold on SSMD allowed NCLUSION to filter out less relevant genes, narrowing the list of cluster-specific over-expressed genes to those most salient.

To further evaluate marker gene quality, we computed gene module scores in order to compare the normalized expression for signature genes across clusters (Fig 4D and Table S6). Here, we find that each module exhibits the highest expression within its respective cluster, with the most definitive signatures occurring within the B (inferred cluster 1), NK (inferred cluster 2), monocytic (inferred cluster 3), and CD34+ cells (inferred cluster 4) (Fig 4E and Fig S4). In clusters that contained heterogeneous combinations of T cell subpopulations, we still see an increased relative expression among cluster-specific marker genes, although not as distinct as in the other cell types.

As an additional analysis, we compared the similarity between the marker genes identified by NCLUSION with the list of marker genes that are identified by using a *post hoc* differential expression analysis with Seurat. Here, we took the FACS-derived experimental annotations from Zheng et al. ^53^ and found differentially expressed genes by doing a one-versus-all Wilcoxon rank sum test for each cluster (mirroring the typical procedure in a conventional bioinformatic workflow). As expected, this post-selective inference procedure resulted in Seurat identifying a multitude of candidate marker genes for each cell type, even after Bonferroni correction. A direct comparison between NCLUSION and Seurat showed that the proposed Bayesian variable selection approach in the NCLUSION framework results in smaller and more refined transcriptomic signatures for downstream investigation (Fig 4F and Figs S5-S6). For example, NCLUSION identified 134 cluster-specific marker genes for NK cells, 97% of which were also included in the 1,780 marker genes selected *post hoc* by Seurat. In total, an average of 96% of the marker genes identified by NCLUSION were included in the much larger sets of differentially expressed genes selected by Seurat across each of the FACS-annotated cell types.

Over-representation analysis using gene product annotations in Gene Ontology (GO) further confirmed that the selective set of gene modules inferred by NCLUSION reflect known immune cell biology ^63–65^ (Fig 4G-H and Table S7). For example, in the cluster containing predominantly B cells (inferred cluster 1), we observed an up-regulation of B cell receptor signaling (adjusted *P* = 6.68*×*10^*−*7^), B cell activation (adjusted *P* = 2.3*×*10^*−*7^), and B cell proliferation (adjusted *P* = 2.51*×*10^*−*7^) ^66,67^. Notably, some of the biologically relevant GO terms found when using NCLUSION were not statistically significant when applying the larger sets of marker genes provided *post hoc* by Seurat. For instance, in the cluster with NK cells (inferred cluster 2), significant gene sets from NCLUSION included the positive regulation of leukocyte chemotaxis (adjusted *P* = 1.53*×*10^*−*4^) and positive regulation of natural killer cell chemotaxis (adjusted *P* = 4.88*×*10^*−*6^) ^68–70^. We also observed enrichment of the up-regulation of natural killer cell mediated cytotoxicity (adjusted *P* =3.81*×*10^*−*5^), consistent with the known highly cytotoxic behavior of NK cells ^68,71^. Each of these gene sets was insignificant when using differentially expressed genes from Seurat (Fig 4G). Lastly, in the monocyte-dominated cluster (inferred cluster 3), the NCLUSION-generated module was enriched for macrophage activation involved in immune response (adjusted *P* = 2.41*×*10^*−*5^) and antigen-presenting activity (e.g., MHC Class II antigen presentation, adjusted *P* = 5.39*×*10^*−*4^). Complete marker gene and GO analyses for all clusters inferred by NCLUSION and Seurat as a baseline in the PBMC dataset can be found in Table S7. Together, these results demonstrate that NCLUSION can identify cluster-specific gene signatures that reflect underlying cellular phenotypes.

### NCLUSION’s performance generalizes to the other single-cell datasets

Finally, we assessed the generalizability of NCLUSION by testing it on three additional large scRNA-seq datasets of various sample sizes: a pancreatic ductal adenocarcinoma (PDAC) dataset from Raghavan et al. ^72^ with *N* = 23,042 cells; an acute myeloid leukemia (AML) dataset from van Galen et al. ^73^ with *N* = 43,690 cells; and a tissue immune (IMMUNE) atlas dataset from Domínguez Conde et al. ^74^ with *N* = 88,057 cells. These datasets represent a range of tissue and disease states to assess our method’s performance in different use cases. Both the PDAC and AML datasets contain a mixture of malignant and non-malignant cells from different patient biopsies, while the IMMUNE dataset contains healthy white blood cells from different anatomical locations. After performing quality control (Methods), we had a total of 5,000 genes with the highest standard deviation (after log-normalization) for the analysis. We observed similar scalability in the runtime of NCLUSION and competing baselines on all three datasets (Fig 5A). NCLUSION maintains its computational efficiency, now only being slightly outperformed by Seurat on the PDAC and AML datasets due to longer convergence time in its variational EM algorithm. We also found that NCLUSION continued to remain competitive in terms of clustering performance. When quantitatively evaluating the clustering ability of NCLUSION versus the competing baselines using the annotations provided by the original studies, NCLUSION was often statistically significantly better (as determined via a two-sided t-test, *P <* 0.05) according to ARI and NMI across all datasets (Fig 5B-C, Figs S7-S19, and Table S4).

**Fig 5.**
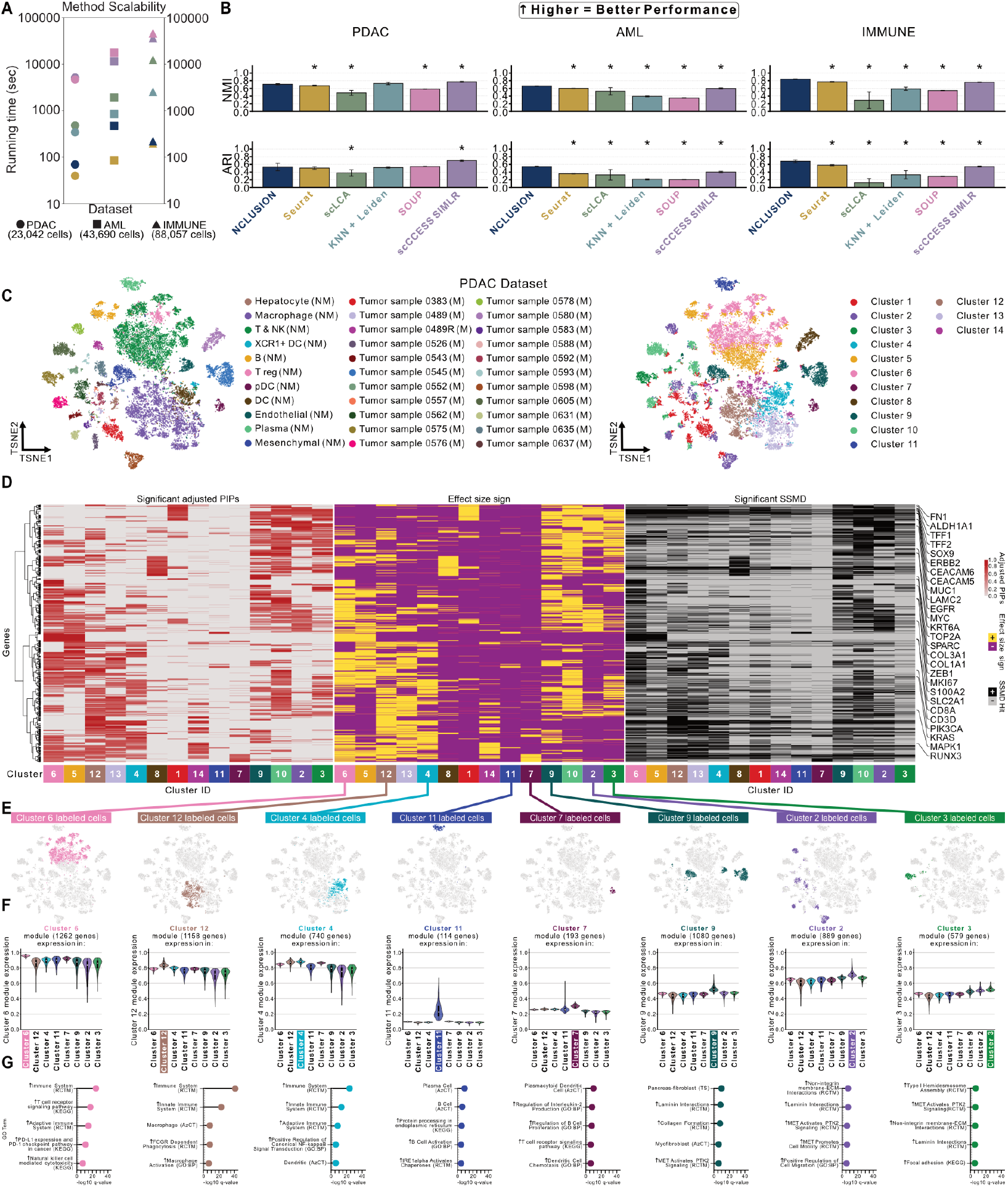
Scalability and generalizability of NCLUSION across diverse datasets. NCLUSION and baselines were applied to the following scRNA-seq datasets: PDAC (*N* = 23,042 cells) ^72^, AML (*N* = 43,690 cells) ^73^, and IMMUNE (*N* = 88,057 cells) ^74^. **(A)** Runtimes for all methods when applied to each dataset. **(B)** Assessment of the inferred cluster labels from each method versus cell type annotations from the original studies. Evaluation is quantified by normalized mutual information (NMI) and adjusted Rand index (ARI). Asterisks indicate that there is a statistically significant difference in performance between NCLUSION and a corresponding method (two-sided t-test *P <* 0.05). Panels **(C)**-**(F)** depict results from running NCLUSION on the PDAC dataset. **(C)** Shown is a t-SNE visualization of the PDAC scRNA-seq dataset, annotated by the cell type labels from the PDAC study (top) compared to the clusters inferred by NCLUSION (bottom), where the “NM” labels indicate non-malignant cells and the “M” labels indicate malignant cells. **(D)** Heat maps of the adjusted posterior inclusion probabilities (PIPs) (left), effect size sign (ESS) (center), and strictly standardized mean difference (SSMD) (right) of the significant genes in each cluster. **(E)** Highlighted location on t-SNEs of NCLUSION-inferred clusters that contain predominantly one cell type. **(F)** Violin plots comparing the normalized expression of cluster-specific marker genes across clusters. **(G)** Gene ontology (GO) pathway enrichment analysis for the marker genes identified for each cluster. Gene sets with a *q*-value below 0.05 are deemed to be significant.

We then analyzed the interpretability of the cluster-specific marker genes inferred by NCLUSION (Fig 5D-G, Figs S8-S24, and Tables S8-S19). For brevity, we highlight just notable results from the PDAC and IMMUNE datasets in the main text. Additional analyses for the AML dataset can be found in the Supplementary Material (see Figs S12-S17 and Tables S16-S19).

To begin, we focused on evaluating the gene modules generated from the NCLUSION inferred clusters in the PDAC dataset (Fig 5D-G). As with the PBMC data, we observed higher module expression within the respective clusters (Fig 5F). NCLUSION appeared to use immune cell signatures as the primary axis for distinguishing malignant and non-malignant populations (Fig 5E and Table S10). For example, the inferred cluster 6 predominantly contained cells that were originally annotated as NK and T cells by Raghavan et al. ^72^. This inferred cluster had immune cell type specific marker genes such as *CD2* (PIP = 0.92), *GZMB* (PIP = 0.85), *IL7R* (PIP = 0.85), and *NCAM1* (PIP = 1.00) ^75,76^ (Fig 5D). An additional GO analysis of this cluster showed an enrichment of natural killer cell mediated cytotoxicity (adjusted *P* = 6.60*×*10^*−*9^) and T cell receptor signaling (adjusted *P* = 1.17*×*10^*−*21^) (Fig 5G).

Notably, the other clusters that primarily contained non-malignant cells (inferred clusters 4, 5, 11, 12, 13, and 14) also directly aligned with cell type labels originally annotated by Raghavan et al. ^72^. For the clusters that primarily contained malignant and metastatic cells (i.e., inferred clusters 1, 2, 3, 9, and 10), a GO analysis revealed an enrichment of extracellular matrix (ECM) organization and cell migration processes (Fig 5G). Importantly, however, NCLUSION also had the power to divide these cells into more granular subpopulations based on their level of differentiation. For example, the inferred cluster 9 was enriched for both cell migration processes (e.g., MET-activated PTK2 signaling, adjusted *P* = 2.97*×*10^*−*5^; MET-promoted cell motility, adjusted *P* = 4.02*×*10^*−*5^) and fibroblast cell activity (pancreatic fibroblasts, adjusted *P* = 2.1*×*10^*−*9^; collagen formation, adjusted *P* = 1.13*×*10^*−*7^) ^77,78^.

Finally, we evaluated the granularity of NCLUSION’s clustering on the IMMUNE dataset, which contained 33 manually annotated labels from experts ^74^. When analyzing this study, NCLUSION was able to delineate between multiple T cell sub-lineages, whereas methods like Seurat, KNN+Leiden, and scCCESS SIMLR merged these subpopulations into 1 or 2 clusters. For example, the inferred cluster 12 by NCLLUSION was enriched (*∼*95% occupancy rate) for CD8+ effector memory (T_EM_) and effector memory cells re-expressing CD45RA (T_EMRA_), while NCLUSION’s inferred cluster 8 was enriched (*∼*90% occupancy rate) for CD8+ tissue-resident memory (T_RM_). These two populations have been shown to be functionally distinct subpopulations in the CD8+ T cell lineage ^79,80^. Likewise NCLUSION’s inferred clusters 6 and 10 were enriched (*∼*77% and *∼*72% occupancy rates, respectively) for functionally distinct subpopulations of CD4+ T cells, namely T follicular helper cells (Tfh) and CD4+ effector/effector memory T cells, respectively ^79,81,82^ (Table S12).

The GO analysis of NCLUSION-generated gene modules also showed an enrichment of T cell phenotypes. The inferred cluster 12’s top ontology term was indeed “CD8+ Effector Memory T4” (adjusted *P* = 4.18*×*10^*−*9^), while the inferred cluster 10 had “CD4+ Central Memory T1” as a top enriched term (adjusted *P* = 3.53*×*10^*−*3^). Similar results showing how NCLUSION-generated gene modules granularly distinguish these cellular populations can be found in the Supplementary Material (Table S15).

## Discussion

We present NCLUSION: a scalable Bayesian nonparametric framework designed to serve as an unbiased method for inferring phenotypic clusters and identifying cluster-specific marker genes in scRNA-seq experiments. We show how our approach simplifies traditional single-cell transcriptomic workflows, which often rely on the transformation of the data to a lower-dimensional representation to facilitate clustering and iteratively tune the number of clusters used to obtain optimal results. In contrast, NCLUSION operates on the full normalized gene expression matrix, eliminating the need for transformation to a lower-dimensional space; infers the optimal number of clusters without iterative user refinement; and simultaneously identifies the cluster-specific marker genes that significantly drive the clustering. By leveraging a variational inference algorithm, NCLUSION can scale to scRNA-seq studies with a million cells. Through the analysis of a collection of large-scale publicly available datasets, we show that NCLUSION not only achieves clustering performance comparable to state-of-the-art methods but also provides refined sets of gene candidates for downstream analyses. By unifying clustering and marker gene selection, NCLUSION provides a flexible and unified statistical framework for inferring complex differential gene expression patterns observed in heterogeneous tissue populations ^21,83,84^.

The current implementation of the NCLUSION framework offers many directions for future development and applications. First, NCLUSION assumes a normal mixture model for log-normalized gene expression data. We use this assumption both because log-based transformations have been shown to reduce the effects of sparsity in single-cell analyses ^20,85^ and because the Gaussian-based specification offers computational advantages for scalable posterior inference. Still, future extensions of the NCLUSION framework should explore the utility of Poisson-and negative binomial-based likelihoods to deal with the zero-inflated nature of scRNA-seq studies in their raw form.

Second, the current formulation of NCLUSION models the gene expression of each cell independently and does not consider, for example, the correlation between genes with similar functionality or co-expression patterns between genes within the same signaling pathway. One possible extension of NCLUSION would be to incorporate additional genomic information into the sparse prior distributions used for Bayesian variable selection. For example, previous studies have proposed an integrative approach where the importance of a variable also depends on an additional set of covariates ^40,86,87^. In the case of single-cell applications, we could assume that the prior probability of the *j*-th gene being a marker of the *k*-th cluster is also dependent upon its cellular pathway membership. Unlike the current spike-and-slab prior NCLUSION implements, this new prior would assume that biologically related pathways contain shared marker genes, essentially integrating the concept of gene set enrichment analysis into clustering. An alternative approach would be to extend NCLUSION to incorporate non-diagonal correlation structures by exploring sparse covariance models, which could provide a balance between the need for maintaining computational efficiency while representing a richer set of gene dependencies ^88,89^.

Third, although it helps NCLUSION scale to large datasets, variational expectation-maximization (EM) algorithms are known to both produce slightly miscalibrated parameter estimates and underestimate the total variation present within a dataset ^41,90,91^. While this does not greatly affect the performance of NCLUSION in the evaluations presented in this paper, this can be seen as a limitation depending on the application of interest. For example, in the PBMC dataset, NCLUSION is unable to resolve all the different T cell subtypes that were annotated by Zheng et al. ^53^ (Fig 3). This is most likely due to variational approximations being well-suited to describe the global variation across cells but at the cost of smoothing over local variation between smaller subpopulations. Considering other (equally scalable) ways to carry out approximate Bayesian inference may be relevant for future work ^92^.

Lastly, a thrust of recent work in genomics has been to develop methods that identify spatially variable marker genes as a key step during analyses of spatially-resolved transcriptomics data ^93^. Future efforts could extend NCLUSION to this emerging modality by, for example, reformulating the method as a spatial Dirichlet process mixture model ^94^.

In sum, NCLUSION provides a unified framework for simultaneous clustering and marker gene selection in single-cell transcriptomic data, yielding improvements in computational efficiency, interpretability, and scalability. We envision that NCLUSION will accelerate key analytic steps universal to single-cell analysis across diverse applications.

## Materials and methods

### Overview of NCLUSION

We provide a brief overview of the probabilistic framework underlying the “Nonparametric CLUstering of SIngle-cell populatiONs” (NCLUSION) model. Detailed derivations of the algorithm are provided in the Supplementary Material. Consider a study with single-cell RNA sequencing (scRNA-seq) expression data for *n* = 1, …, *N* cells that each have measurements for *j* = 1, …, *J* genes. Let this dataset be represented by the *N × J* matrix **X** where the row-vector **x**_*n*_ = (*x*_*n*1_, …, *x*_*nJ*_) denotes the expression profile for the *n*-th cell. We assume that the log-normalized gene expression for each cell follows a sparse hierarchical Dirichlet process normal mixture model ^31–33^ of the form

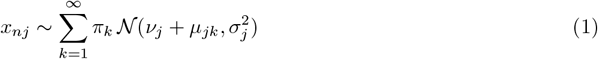

where *π*_*k*_ represents the marginal (unconditional) probability that a cell belongs to the *k*-th cluster, *ν*_*j*_ and 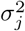 are the global means and variances for the *j*-th gene across all cells (i.e., not conditioned on cluster identity), and *µ*_*jk*_ is the mean shift of expression for the *j*-th gene within the *k*-th cluster. There are two key features in the model formulation of NCLUSION specified above. First, we assume that the formation of clusters is driven by a few important genes that have mean expression shifted away from a baseline gene-specific expression level, *ν*_*j*_. To that end, we place a sparsity-inducing spike and slab prior distribution on the mean effect of each gene

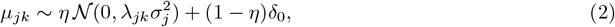

where *δ*_0_ is a point mass at zero, *λ*_*jk*_ scales the global variance to form a cluster-specific “slab” distribution for each gene, and *η* is the prior probability that any given gene has a nonzero effect when assigning a cell to any cluster. In practice, there are many different ways to estimate *η*. Following previous work ^40,41,95–97^, one choice would be to assume a uniform prior over log *η* to reflect our lack of knowledge about the correct number of “marker” genes for each cell type that is present in the data. Instead, in this work, we assume *η ∼* Beta(1, 1) to represent this uncertainty and learn its value during model inference. To facilitate posterior computation and interpretable inference, we introduce a binary indicator variable *ρ*_*jk*_ *∈ {*0, 1*}*

where we implicitly assume *a priori* that Pr[*ρ*_*jk*_ = 1] = *η*. Alternatively, we say that *ρ*_*jk*_ takes on a value of 1 when the effect of a gene *µ*_*jk*_ on cluster assignment is nonzero and deviates from the baseline gene expression level *ν*_*j*_. As NCLUSION is trained, the posterior mean for unimportant genes will trend towards the global mean (i.e., *µ*_*jk*_ *→* 0) as the model attempts to identify subsets of marker genes that are relevant for each cluster. We then use posterior inclusion probabilities (PIPs) as general summaries of evidence that the *j*-th gene is statistically important in determining when a cell is assigned to the *k*-th cluster where

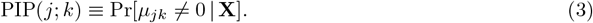

The second key feature in Eq. (1) is that we do not assume to know the true number of clusters *K*. Instead, we take a nonparametric approach and attempt to learn *K* directly from the data. Once again, to facilitate posterior computation, we introduce a categorical latent variable *ψ*_*n*_ which indicates that the *n*-th cell is in the *k*-th cluster with prior probability *π*_*k*_. Explicitly, we write this as Pr[*ψ*_*n*_ = *k*] = *π*_*k*_. Here, we implement the stick-breaking construction of the Dirichlet process ^31^ where we say

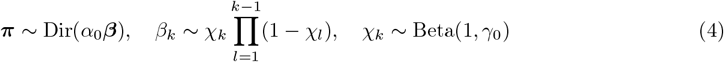

with ***π*** = (*π*_1_, …, *π*_*>K*_) having mean ***β*** and variance determined by the concentration hyper-parameters *α*_0_ and *γ*_0_ ^98^. The concentration hyper-parameters *α*_0_ and *γ*_0_ are both non-negative scalars that effectively help to determine the number of clusters used in the model ^31,39^. Larger values for these parameters increase the model’s sensitivity to variation in the data and encourage the creation of a greater number of smaller clusters. Smaller values for these parameters, on the other hand, decrease the model’s sensitivity to variation in the data and encourage the creation of fewer larger clusters. In this work, we encourage the creation of fewer clusters and fix *α*_0_ and *γ*_0_ to be less than or equal to 1 (Supplementary Material). After model training, we use the posterior distribution over the latent categorical indicators Pr[*ψ*_*n*_ = *k* | **X**] to determine the cluster assignment for each cell. It is worth noting that, although the prior number of normal components is infinite in Eq. (1), the posterior number of components after model fitting will be finite. This truncation reflects the fact that not all infinite states are used when conditioning on finite data ^98^. Additionally, the algorithm used to estimate the parameters in the NCLUSION software penalizes empty clusters (see Supplementary Material), and, as a result, the model has the flexibility to automatically adjust its complexity based on the inferred complexity of the data being analyzed. This helps to increase the utility and adaptability of NCLUSION across a wide range of single-cell applications.

### Selection of cluster-specific marker genes

NCLUSION jointly performs clustering on single-cell populations while also learning cluster-specific gene signatures. To achieve this, we use the spike and slab prior distribution specified in Eq. (2) and the resulting PIPs defined in Eq. (3) to find the most salient genes per cluster. Since the model fits to each *j*-th gene expression for the *n*-th cell independently, signatures learned between clusters can share subsets of the same genes. Genes that are identified as “important” across many different clusters can effectively be seen as ubiquitous housekeeping variables rather than significant marker genes of unique cell types. Therefore, we down-weight the inclusion probabilities to proportionally penalize genes based on the number of clusters in which they appear

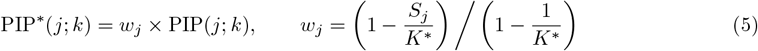

where *K*^*∗*^ *≤ K* is the finite number of occupied clusters learned by the model, and *S*_*j*_ is the number of clusters that the *j*-th gene is significant in according to a given selection threshold. We set this threshold to be 0.5 which corresponds to the median probability criterion in Bayesian statistics ^38^.

While Eqs. (3) and (5) can be used to identify the genes that are differentially expressed in a given cluster, they do not indicate the direction or magnitude of this shift. Therefore, for each gene, we combine the adjusted posterior inclusion probabilities with effect size sign (ESS) and strictly standardized mean difference (SSMD) measures to find the most salient markers per cluster. Here, we obtain the effect size sign by taking the sign of Cohen’s *d* ^99^ between the expression of the *j*-th gene for cells in the *k*-th cluster and cells not in the *k*-th cluster (denoted by *k*^*′*^)

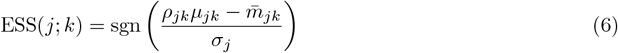

where, in addition to previous notation, 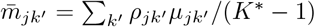 is the average mean shift for the *j*-th gene in all clusters outside of the *k*-th. Here, sgn(*·*) is the piecewise sign function where sgn(*u*) = + (i.e., positive) when *u >* 0, sgn(*u*) = *−* (i.e., negative) when *u <* 0, and sgn(*u*) = 0 when *u* = 0.

The strictly standardized mean difference (SSMD) is a metric often used in high-throughput screenings to test for the significance of an effect size magnitude ^100–103^. It is computed as the following

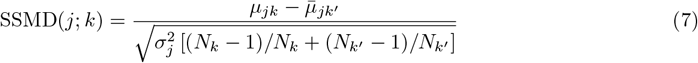

where 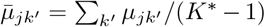 is the average global mean for the *j*-th gene in all clusters outside of the *k*-th. Asymptotically, the SSMD follows a normal distribution ^100,101,103^. To determine a significant value, we follow a previous procedure ^103^ by calculating a threshold |SSMD(*j*; *k*)| *≥ S*^*∗*^(*j*; *k*) which controls for a predetermined false positive rate (FPR). Here, this threshold is given by

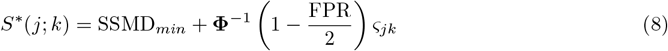

where FPR is set to 0.05, **Φ**^*−*1^(*·*) is the inverse cumulative distribution function of a standard normal, and SSMD_*min*_ is the minimum SSMD magnitude that one considers to be significant. In practice, this minimum value is often set between 0 and 0.25 in order to identify weak effect sizes. In the main text, we follow previous work ^103–105^ and let SSMD_*min*_ = 0.15. The parameter *ς*_*jk*_ is used to denote the asymptotic variance which is given by

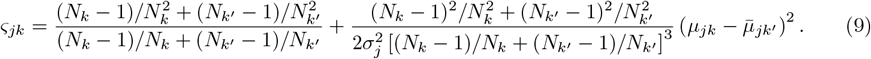

In the main text, cluster-specific marker genes are selected as those that have a significant adjusted inclusion probability and are notably up-regulated in a given cluster meaning that they satisfy the following criteria: (1) PIP^*∗*^(*j*; *k*) *≥* 0.5, (2) ESS(*j*; *k*) = +, and (3) SSMD(*j*; *k*) *≥ S*^*∗*^(*j*; *k*), respectively.

### Posterior inference via variational EM algorithm

We combine the likelihood in Eq. (1) and the prior distributions in Eqs. (2) and (4) to perform Bayesian inference. In current scRNA-seq datasets, it is less feasible to implement traditional Markov Chain Monte Carlo (MCMC) algorithms due to the large number of cells being studied. For model fitting, we instead use a variational expectation-maximization (EM) algorithm ^31,32,98^, which allows us to estimate parameters within an optimization framework. The overall goal of variational inference is to approximate the true posterior distribution for model parameters using a set of approximating distributions. The EM algorithm optimizes parameters such that it minimizes the Kullback-Leibler divergence between the exact and approximate posterior distributions. To compute the variational approximations, we make the mean-field assumption that the true posterior can be “fully-factorized” ^106^. The algorithm then follows two general steps. In the first step, we iterate through a combination of hyper-parameter values and compute variational updates for the other parameters using coordinate ascent. In the second step, we empirically compute (approximate) posterior values for the main model parameters *{****µ, ρ, ψ****}*. Detailed steps in the variational EM algorithm, explicit coordinate ascent updates for the model parameters, pseudocode, and other derivations are given in the Supplementary Material. Parameters in the variational EM algorithm are initialized by taking random draws from their assumed prior distributions. Iterations in the algorithm are terminated when at least one of two stopping criteria are met: (i) the difference between the lower bound of two consecutive updates is within some small range (specified by argument *ϵ*), or (ii) a maximum number of iterations is reached. For the analyses run in this paper, we set *ϵ* = 1 for the first criterion and used a maximum of 1 *×* 10^4^ iterations for the second.

### Simulation study design

#### Generating simulated datasets

To evaluate the robustness and sensitivity of NCLUSION under controlled conditions, we generated synthetic single-cell RNA-seq datasets using `scDesign3`^47^ (v.1.4.0). The reference dataset we used was derived from the FACS-sorted peripheral blood mononuclear cell (PBMC) dataset produced by Zheng et al. ^53^. Initial preprocessing for this reference dataset included mitochondrial gene content assessment, ribosomal and hemoglobin gene filtering, and quality control to remove both low-quality cells and lowly expressed genes. Highly variable genes (HVGs) were identified using the `modelGeneVar` function in the `scran R` package, and the top 1000 HVGs were retained for downstream simulation. We used five immune cell types (B cells, CD14+ monocytes, CD56+ natural killer (NK) cells, cytotoxic T cells, and regulatory T cells) for these analyses. To ensure balanced representation across cell types, we implemented a stratified subsampling scheme which selected an equal number of cells per type while enforcing non-zero gene expression across all selected cells and genes.

Each simulated dataset comprised of *N* = 10,000 cells across five clusters and 1,000 genes where we preserved realistic transcriptomic correlation structures through Gaussian copula modeling. Simulations

were conducted across four different scenarios (with 20 replicates per scenario), each varying in cluster size imbalance and marker gene composition.

- **Scenario I:** Balanced clusters of 2000 cells per cell type, each with 50 marker genes.
- **Scenario II:** Imbalanced cluster design where one small cluster had 200 cells and the other four larger clusters each had 2450 cells. All clusters contained 50 marker genes.
- **Scenario III:** Imbalanced cluster design where one cluster had 20 (rare) cells and the other four larger clusters each had 2495 cells each. All clusters contained 50 marker genes.
- **Scenario IV:** Balanced clusters of 2000 cells per cell type, but one cluster had only 20 marker genes while the other four clusters had 50 marker genes.

More specifically, synthetic datasets were generated using the `construct_data, fit_marginal, fit _copula, extract_para`, and `simu_new` functions within `scDesign3` to create gene expression vectors using a negative binomial distribution that is conditioned on cell type from the reference data. To introduce differentially expressed genes (DEGs), we first ranked genes by their cell type specific mean expression in the reference data and sampled a number of top-ranked genes to be markers. Then in the synthetic data, these DEGs were then artificially upregulated in one cluster while maintaining the baseline expression in others. This was done by apply a log-fold change factor sampled uniformly over the interval [1.5, 2.5]. This ensured that we maintained realistic variance but still had distinct signal between cell types.

### Real datasets and preprocessing

Below we briefly describe all of the datasets and the preprocessing steps used in this work. Each of these datasets is relatively large (containing at least 20,000 cells) with unique molecular identifiers (UMI). The latter is important because prior research suggests that UMIs provide enough information to avoid overcounting issues due to amplification and zero-inflation ^14,107,108^. We use an asterisk by the BRAIN-LARGE dataset to indicate that it was exclusively to test the scalability of NCLUSION and competing methods; therefore, clustering performance was not recorded. For the other datasets, we use cell type annotations provided by the original study as “true” reference labels during our analyses. Cells were filtered for quality using a custom `scanpy`^109^ (v.1.9.1) pipeline script (see Software availability). Unless otherwise stated, all data was preprocessed by taking the logarithm (to the base 2) of the counts, dividing by a scaling factor of 10000, and then adding a pseudo-count of 1.0 for stability. Additionally, unless otherwise stated, all results were produced using the top 5000 highly variable genes (HVG), which were determined by sorting the standard deviation of the transformed counts ^33^.

#### BRAIN-LARGE*

This dataset originally contains 1.3 million mouse brain cells from 10x Genomics ^45^. During preprocessing, we subset the data to a collection of 720 genes following a procedure outlined by Lopez et al. ^14^. Next, we further filtered by only keeping cells that had at least one of these genes expressed. This left a total of 64,071 cells. Since the original study did not provide cell labels, we exclusively used this dataset to compare runtime performance. To do so, we up-sampled by randomly selecting groups of 64,071 cells to create a synthetic dataset of 1 million cells. We report the runtime for each method on datasets with 500, 1K, 5K, 10K, 50K, 100K, 500K, and 1M cells.

#### PBMC

We took scRNA-seq data from fluorescence-activated cell sorted (FACS) populations of peripheral blood mononuclear cells (PBMCs) provided by Zheng et al. ^53^ and concatenated each population into one dataset. During preprocessing, we filtered out genes that were expressed in fewer than three cells. We also dropped cells with (i) fewer than 200 genes expressed, (ii) greater than 20% mitochondrial reads, and (iii) fewer than 5% ribosomal reads. This resulted in a final dataset with 94,615 high-quality cells representing 10 distinct cell types.

#### PDAC

We used scRNA-seq data from pancreatic ductal adenocarcinoma (PDAC) tissue obtained from 23 patients according to methods documented in Raghavan et al. ^72^. This dataset contains 23,042 total cells made up of 15,302 non-malignant cells of 11 distinct cell types and 7,740 malignant cells.

#### AML

The scRNA-seq data obtained from van Galen et al. ^73^ contains 43,690 acute myeloid leukemia (AML) and non-malignant donor cells taken from 16 AML patients, 5 healthy donors, and 2 cell lines. It is comprised of 13,489 patient-derived malignant cells, 23,005 non-malignant donor cells, 6,018 cells from the MUTZ-3 AML cell line, and 1,178 cells from the OCI-AML3 cell line. To account for the biological differences between cell lines and donor cells of the same cell type annotation, we appended the cell line name onto cell type labels where applicable, producing 33 distinct cell types overall. To process the data, we filtered out all cells with “unclear” cell state labels, retaining only “malignant” or “non-malignant” cells.

#### IMMUNE

We obtained filtered scRNA-seq data from approximately 330,000 immune cells from 12 organ donors in Domínguez Conde et al. ^74^. To mitigate batch effects, we isolated 88,057 cells that were taken from a single organ donor (donor D496) with uniform chemistry annotations, containing 44 distinct immune cell types.

### Other methods

We selected five additional methods to compare against the performance of NCLUSION in real data and in simulations: (1) a Louvain algorithm implemented using the FindClusters function in Seurat ^9^ (v.4.3.0.1); (2) a spectral clustering method called scLCA ^10^ (v.0.0.0.9000), which optimizes both intra- and inter-cluster similarity; (3) a combination of a K-Nearest Neighbor (KNN) classifier with the Louvain community detection algorithm to find clusters implemented via `scikit-learn`^43^ (v.1.2.2) and `scanpy`^109^ (v.1.9.1), respectively; (4) a semi-soft clustering algorithm called SOUP ^44^ (v.0.0.0.9000); and (5) an ensemble method called scCCESS-SIMLR (v.0.2.1), which leverages the spectral clustering approach SIMLR ^11,110^. In the simulation experiments, we also compared the clustering performance of NCLUSION against three additional methods: (6) a consensus clustering method, SC3^49^(v.1.34.0); (7) a deep-learning based method, scDeepCluster ^50^ (v.1.0.0); and (8) an imputation and dimensionality reduction method, CIDR ^48^ (v.0.1.5). Also in simulations, when assessing the ability of NCLUSION to perform robust marker gene selection, we compare it against: (9) a method that leverage differential correlation patterns in the local structure of a PCA-derived cell neighborhood graph, DUBStepR ^51^ (v.1.2.0); (10) a feature selection via an EM algorithm, FESTEM ^52^ (v.1.2.1); and (11) a divergence-based strategy with permutation tests, singleCellHaystack ^25^ (v.1.0.2). Additional details about each method are provided in the Supplementary Material.

### Evaluation metrics

Below we describe the metrics and approaches used to compare performance across all methods. Our clustering evaluation procedure used extrinsic metrics that require reference labels to serve as the ground truth in our calculations.

#### Normalized mutual information (NMI)

This metric is a normalized variant of mutual information (MI). It is an entropy-based metric that captures the amount of shared information between the inferred label distribution and the reference label distribution. NMI ranges between [0, 1] where 1 represents total information sharing between label sets and 0 represents no information sharing between label sets. NMI is calculated by

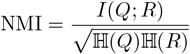

where *Q* and *R* are the empirical label distributions from the inferred and reference labels, respectively. The function *I*(*·*) is the mutual information between the inferred labels distribution and reference labels distributions; H(*·*) represents the Shannon entropy of a given label distribution ^110,111^.

#### Adjusted Rand index (ARI)

This metric captures the similarity between labels inferred by a method and the reference labels. It is based on the Rand index (RI) but corrects for the measurement’s sensitivity to chance. ARI ranges between [-1, 1] where 1 represents perfect agreement between label sets, 0 represents random agreement, and -1 represents perfect disagreement. ARI is calculated by

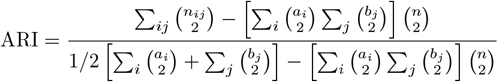

where *n*_*ij*_, *a*_*i*_, and *b*_*j*_ are values obtained from a contingency table, and *n* = ∑_*ij*_ *n*_*ij*_ ^110–112^.

#### Metrics to evaluate marker gene selection in simulations

In the simulation studies, we evaluated the accuracy of marker gene detection of NCLUSION and competing methods by treating the task as a classification problem. In order to do so, we defined the confusion matrix defined below.

**Table 1.** Confusion matrix showing true and inferred marker gene labels.

|  |  | (Inferred Label) |  | Total |
| --- | --- | --- | --- | --- |
|  |  | Marker gene | Non-marker gene |  |
| (True Label) | Marker gene | TP | FN | $b_1$ |
| | Non-marker gene | FP | TN | $b_2$ |
| Total | | $a_1$ | $a_2$ | $n$ |

Here, TP represents the number of correctly identified marker genes (true positives), FN represents the number of incorrectly identified marker genes (false negatives), TN represents the number of correctly identified non-marker genes (true negatives), and FP represents the number of incorrectly identified non-marker genes (false positives). In the table above, we let *a*_1_ and *a*_2_ be the total number of genes inferred as markers and the total number of genes not inferred as markers, respectively. Likewise, we let *b*_1_ and *b*_2_ be the total number of genes that are truly markers and non-markers, respectively. It follows that the total number of genes is defined as *n* = *a*_1_ + *b*_1_ + *a*_2_ + *b*_2_. From here, we can compute the following metrics.

- **True positive rate (TPR; also referred to as power)** captures the proportion of correctly identified marker genes using a given method. It is defined as TPR = TP/(TP+FP).
- **False discovery rate (FDR)** details the proportion of all identified marker genes that are actually non-marker genes It is defined as FDR = FP/(TP + FP).
- **False positive rate (FPR)** captures the proportion of non-marker genes that will be incorrectly labeled marker genes. It is defined as FP/(TN + FP).

Note that the false positive rate can also be computed as FPR = 1-Specificity.

#### Normalized module expression

Genes with significantly adjusted PIPs in Eq. (5), positive ESS in Eq. (6), and significant SSMD in Eq. (7) were used to generate modules (i.e., a collection of marker genes) for each cluster. We calculated a score for each cluster to asses the exclusivity of expression within each module. This was done using the `score_genes` function in `scanpy` (v.1.10.4). The violin plots were generated using the `violinplot` function in matplotlib (v.3.10.0).

#### Gene set over-enrichment analysis

We also performed gene set enrichment analysis on each of the learned gene modules across clusters. This was done via an over-enrichment analysis within the `GSEApy` package ^113^ (v.1.1.5) in Python (v.3.11.0). This method uses a hypergeometric test to calculate the enrichment of genes in a supplied module with respect to the gene sets within an ontology. In this work, we use the ontology labeled `GO_Biological_Process 2025`^63–65,114^, `Tabula_Sapiens`^115,116^, `Azimuth_Cell_Types_2021`^117^, `KEGG_2021_Human` ^118–120^, and `Reactome_2022`^121–128^. The gene sets in this particular ontology represent a combination of biological processes, pathways, and phenotypes. In this analysis, we use *q*-values to determine the enrichment of a given gene set with a significance threshold set to 0.05. The *q*-value is the analog of a *p*-value that has been corrected for testing multiple hypotheses (i.e., an adjusted *P*).

## Software availability

An open-source software implementation of NCLUSION is available on GitHub athttps://github.com/microsoft/Nclusion.jl. Guided tutorials and all code needed to reproduce the results and figures in this work can be found at https://microsoft.github.io/Nclusion.jl/.

## Data availability

All of the datasets analyzed in this paper are publicly available. The PDAC dataset from Raghavan et al. ^72^ can be accessed at https://singlecell.broadinstitute.org/single_cell/study/SCP1644/microenvironment-drives-cell-state-plasticity-and-drug-response-in-pancreatic-cancer#/. The AML data from van Galen et al. ^73^ can found at https://www.dropbox.com/s/399x045zc57fiut/Seurat_AML.rds?dl=0. The BRAIN-LARGE dataset can be accessed at https://www.10xgenomics.com/datasets/1-3-million-brain-cells-from-e-18-mice-2-standard-1-3-0. The individual PBMC data from Zheng et al. ^53^ can be downloaded directly from https://www.10xgenomics.com/resources/datasets. Lastly, the immune cell atlas dataset can be accessed at https://cellgeni.cog.sanger.ac.uk/pan-immune/CountAdded_PIP_global_object_for_cellxgene.h5ad.

## Acknowledgements

We thank members of the Crawford, Raghavan, and Shalek Labs for insightful comments on earlier versions of this manuscript. This research was conducted by using a combination of computational resources and services provided by Microsoft Research and the Center for Computation and Visualization at Brown University. This research was also supported in part by an Alfred P. Sloan Research Fellowship and a David & Lucile Packard Fellowship for Science and Engineering awarded to LC. CN was a trainee supported under the Brown University Predoctoral Training Program in Biological Data Science (NIH T32GM128596). SR is supported by NCI K08 award 1K08CA260442, the Claudia Adams Barr Program in Innovative Basic Cancer Research, and the Dana-Farber Cancer Institute Hale Family Center for Pancreatic Cancer Research. Any opinions, findings, and conclusions or recommendations expressed in this material are those of the author(s) and do not necessarily reflect the views of any of the funders.

## Author contributions

CN, APA, and LC conceived the study and developed the methods. CN and MH developed the algorithm and software. CN, MH, and MR led the analyses. SR, PSW, APA, and LC provided resources, supervised the project, and conducted secondary analyses. All authors interpreted the results and wrote and revised the manuscript.

