## Supplementary Text, Figures, and Tables for "Scalable nonparametric clustering with unified marker gene selection for single-cell RNA-seq data"

### 1 Overview of Hierarchical Bayesian Nonparametric Clustering

In this section, we detail the probabilistic model underlying the "Nonparametric CLustering Of SIngle cell populatiONs" (NCLUSION) framework. Assume that we have a study with single-cell RNA sequencing (scRNA-seq) expression data for $n = 1, \dots, N$ cells that each have measurements for $j = 1, \dots, J$ genes. We will denote this data as an $N \times J$ matrix $\mathbf{X}$ with the row-vector $\mathbf{x}_n = (x_{n1}, \dots, x_{nJ})$ representing the expression for the $n$-th cell. The NCLUSION methodology assumes that the log-normalized gene expression for each cell follows a sparse hierarchical Dirichlet process normal mixture model [1–3]

$$x_{nj} \sim \sum_{k=1}^{\infty} \pi_k \mathcal{N}(\nu_j + \mu_{jk}, \sigma_j^2), \tag{1}$$

where $\pi_k$ represents the marginal (unconditional) probability that a cell belongs to the $k$-th cluster, $\nu_j$ and $\sigma_j^2$ are the global means and variances for the $j$-th gene across all cells (i.e., not conditioned on cluster identity), and $\mu_{jk}$ is the mean shift of expression for the $j$-th gene within the $k$-th cluster. The framing of NCLUSION as a nonparametric mixture model facilitates our ability to simultaneously perform classic variable selection when determining cluster assignments. Here, we assume that only a few genes are "markers" in defining cell types or cellular states. To do this, we place a sparsity-inducing spike and slab prior distribution on the cluster means

$$\mu_{jk} \sim \eta \mathcal{N}(0, \lambda_{jk}\sigma_j^2) + (1 - \eta)\delta_0, \qquad \Pr[\rho_{jk} = 1] = \eta \tag{2}$$

where $\delta_0$ is a point mass at zero, $\lambda_{jk}$ is scaling component of the "slab" distribution, and $\rho_{jk} \in \{0, 1\}$ is an indicator variable signifying that the $j$-th gene has a nonzero effect when assigning a cell to the $k$-th cluster with prior probability $\eta$. Following previous work [4–8], we set $\eta \sim \text{Beta}(\varphi_1, \varphi_2)$ where $\varphi_1 = \varphi_2 = 1$ to reflect our lack of knowledge *a priori* about the number of driver genes per cluster and to represent our assumption that each cluster is formed with at least one marker gene with a nonzero mean. In the main text, we refer to the indicators $\rho_{jk}$ as inclusion probabilities [9] and we use the marginal posterior means of these quantities $\text{PIP}(j; k) \equiv \Pr[\rho_{jk} = 1 \,|\, \mathbf{X}] = \Pr[\mu_{jk} \neq 0 \,|\, \mathbf{X}]$ as general summaries of evidence that a gene is statistically important in determining when a cell is assigned to the $k$-th cluster. In our framework, we treat $\nu_j$, $\sigma_j^2$, and $\lambda_{jk}$ as hyper-parameters that we assume follow conjugate prior distributions

$$\nu_j \sim \mathcal{N}(\omega, \tau^2), \quad \sigma_j^2 \sim \text{Inv-Gamma}(\xi_1, \xi_2), \quad \lambda_{jk} \sim \text{Inv-Gamma}(\kappa_1, \kappa_2) \tag{3}$$

Following previous work from Zeng and Zhou[10], we place a limiting normal prior for each $\nu_j$ by setting $\omega = 0$ and $\tau^2 = 1 \times 10^{12}$). Similarly, following previous work from Hughes et al.[11], we assume relatively uninformative priors with large variances for $\sigma_j^2$ and $\lambda_{jk}$ by setting the shape and scale of the inverse-gamma distributions to be $\xi_1 = \xi_2 = 1$, $\kappa_1 = 0.1$, and $\kappa_2 = 1 \times 10^{-3}$, respectively.

In practice, we do not know the true number of unique cell clusters $K$ in any given study. As a result, we assume that there can effectively be infinitely many clusters *a priori* and attempt to learn $K$ directly from the data during model fitting. Specifically, using the stick-breaking constructive representation of the Dirichlet process[1], we introduce one final latent variable $\psi_n$ to denote a categorical indicator that the $n$-th cell is in the $k$-th cluster with probability $\pi_k$. Here,

$$\psi_n \sim \text{Cat}(\boldsymbol{\pi}), \quad \boldsymbol{\pi} \sim \text{Dir}(\alpha_0 \boldsymbol{\beta}), \qquad \beta_k \sim \chi_k \prod_{l=1}^{k-1}(1 - \chi_l), \qquad \chi_k \sim \text{Beta}(1, \gamma_0) \tag{4}$$

which implies that each element in $\boldsymbol{\pi}$ has mean $\boldsymbol{\beta}$ and a set of variances that are determined by the concentration parameters $\alpha_0$ and $\gamma_0$. During analyses, we use the posterior distribution over the latent categorical indicators $\Pr[\psi_n = k \mid \mathbf{X}]$ to determine cell cluster assignment. In the literature, the latter half of Eq. (4) represents the Griffiths/Engen/McCloskey distribution such that $\boldsymbol{\beta} \sim \text{GEM}(\gamma_0)$ where $\gamma_0$ is another concentration parameter that determines the number of clusters used in the model. As mentioned in the main text, although the prior number of normal components in our model is infinite, the posterior number of components for any given dataset will be finite. In practice, we adequately approximate an infinite Dirichlet process with $K = 25$ cluster components in the stick-breaking construction. The concentration hyper-parameters $\alpha_0$ and $\gamma_0$ are both non-negative scalars that effectively help to determine the final number of clusters $K^*$ that are used in the model[1,10,11]. Larger values for these hyper-parameters encourage the model to create a greater number of small clusters. In this work, we aim to decrease the model's sensitivity to variation in the data and encourage the creation of fewer large clusters. As a result, we fix $\alpha_0$ and $\gamma_0$ to be less than or equal to 1. Altogether, our model has the potential to automatically adjust its complexity based on the inferred complexity of the data being analyzed. This increases the utility of NCLUSION as it can adapt to a wide range of single-cell applications.

#### 1.1 Selection of Marker Genes

As mentioned in the main text, NCLUSION selects a final set of cluster-specific marker genes based on two sets of criteria: ($i$) an adjusted posterior inclusion probability (PIPs) which provides evidence that the $j$-th gene is statistically important in determining when a cell is assigned to the $k$-th cluster and ($ii$) the sign and magnitude of the $j$-th gene's effect which is used to determine whether it is significantly up-regulated or down-regulated within the $k$-th cluster. In the former, we consider genes that are "important" to many clusters to effectively be housekeeping variables rather than unique cell type markers. As a result, we adjust the inclusion probabilities to penalize genes according to the number of clusters in which they appear

$$\text{PIP}^*(j;k) = w_j \times \text{PIP}(j;k), \qquad w_j = \left(1 - \frac{S_j}{K^*}\right) \bigg/ \left(1 - \frac{1}{K^*}\right)$$

where, again, $K^* \leq K$ is the finite number of occupied clusters learned by the model, and $S_j$ is the number of clusters that the $j$-th gene is significant according to a given selection threshold. We set this threshold to be 0.5 which corresponds to the median probability criterion in Bayesian statistics[12]. We combine the adjusted posterior inclusion probabilities with the effect size sign and strictly standardized mean difference (SSMD). The former for each gene is computed by taking the sign of Cohen's $d$[13] between the expression of the $j$-th gene for cells in the $k$-th cluster and cells not in the $k$-th cluster (denoted by $k'$)

$$\text{ESS}(j;k) \simeq \text{sgn}\left(\mu_{jk} - \bar{\mu}_{jk'}\right)$$

where $\bar{\mu}_{jk'} = \sum_{k'} \mu_{jk'}/(K^* - 1)$ is the average mean for the $j$-th gene in all clusters outside of the $k$-th. Here, sgn$(\cdot)$ is the piecewise sign function where sgn$(u) = +$ (i.e., positive or over-expression) when $u > 0$, sgn$(u) = -$ (i.e., negative or under-expression) when $u < 0$, and sgn$(u) = 0$ when $u = 0$. The strictly standardized mean difference (SSMD) tests for the significance of an effect size magnitude[14–17]. It is computed as the following

$$\mathrm{SSMD}(j;k) = \frac{\mu_{jk} - \bar{\mu}_{jk'}}{\sqrt{\sigma_j^2\left[(N_k - 1)/N_k + (N_{k'} - 1)/N_{k'}\right]}} \tag{5}$$

where $\bar{\mu}_{jk'} = \sum_{k'} \mu_{jk'}/(K^* - 1)$ is the average global mean for the $j$-th gene in all clusters outside of the $k$-th. To determine a significant value SSMD value, we compute a threshold is given by

$$S^*(j;k) = \mathrm{SSMD}_{min} + \mathbf{\Phi}^{-1}\left(1 - \frac{\mathrm{FPR}}{2}\right)\varsigma_{jk} \tag{6}$$

which is meant to control for a predetermined false positive rate (FPR) which we set to be 0.05. In addition, $\mathbf{\Phi}^{-1}(\cdot)$ is the inverse cumulative distribution function of a standard normal, and $\mathrm{SSMD}_{min}$ is the minimum SSMD magnitude that one considers to be significant. In the main text, we follow previous work[17–19] and let $\mathrm{SSMD}_{min} = 0.15$. The parameter $\varsigma_{jk}$ is used to denote the asymptotic variance which is given by

$$\varsigma_{jk} = \frac{(N_k - 1)/N_k^2 + (N_{k'} - 1)/N_{k'}^2}{(N_k - 1)/N_k + (N_{k'} - 1)/N_{k'}} + \frac{(N_k - 1)^2/N_k^2 + (N_{k'} - 1)^2/N_{k'}^2}{2\sigma_j^2\left[(N_k - 1)/N_k + (N_{k'} - 1)/N_{k'}\right]^3}\left(\mu_{jk} - \bar{\mu}_{jk'}\right)^2. \tag{7}$$

In the main text, cluster-specific marker genes are selected as those that have a significant adjusted inclusion probability and are notably up-regulated in a given cluster meaning that they satisfy the following criteria: (1) $\mathrm{PIP}^*(j;k) \geq 0.5$, (2) $\mathrm{ESS}(j;k) = +$, and (3) $\mathrm{SSMD}(j;k) \geq S^*(j;k)$, respectively.

### 2 Variational Expectation-Maximization (EM) Algorithm

We use a variational expectation maximization (EM)-like algorithm known as coordinate ascent to estimate the posterior distribution of parameters in the NCLUSION framework. The derivations in this section largely follow those developed in previous work[1,2,11]. As mentioned in the main text, the overall goal of variational inference is to approximate the true posterior distribution for the model parameters $p(\mathbf{\Theta} \mid \mathbf{X})$ with a similar distribution from an approximating family $q(\mathbf{\Theta})$[20–24]. This is done by tuning a set of free parameters until the Kullback-Leibler (KL) divergence between the exact and approximate posterior distributions is minimized. We then run the EM algorithm while iterating through a series of expectation (E) and maximization (M) steps. In the E-step, we use coordinate ascent to update the free parameters of the approximate variational posterior [5]. In the M-step, we derive updates for the model hyper-parameters by solving for the roots of their gradients. A complete overview of the algorithm is given below. Iterations in the NCLUSION software are terminated when either one of two stopping criteria are met: $(i)$ the difference between the lower bound of two consecutive updates are within some small range (specified by tolerance argument $\epsilon$), or $(ii)$ a maximum number of iterations is reached. For the simulations and real data analyses ran in this paper, we set $\epsilon = 1 \times 10^{-4}$ for the first criterion and used a maximum of 10,000 iterations for the second.

To formally derive the variational EM algorithm, first let $\boldsymbol{\Theta} = \{\boldsymbol{\nu}, \boldsymbol{\mu}, \boldsymbol{\sigma}^2, \boldsymbol{\lambda}, \boldsymbol{\rho}, \eta, \boldsymbol{\pi}, \boldsymbol{\psi}, \boldsymbol{\chi}\}$. We begin with the KL divergence which is defined as the following

$$\mathrm{KL}(q(\boldsymbol{\Theta}) \,\|\, p(\boldsymbol{\Theta} \,|\, \mathbf{X})) = \int \ln \left[ \frac{q(\boldsymbol{\Theta})}{p(\boldsymbol{\Theta} \,|\, \mathbf{X})} \right] q(\boldsymbol{\Theta}) \, \mathrm{d}\boldsymbol{\Theta} = \mathbb{E}_{q(\boldsymbol{\Theta})} \left\{ \ln \left[ \frac{q(\boldsymbol{\Theta})}{p(\boldsymbol{\Theta} \,|\, \mathbf{X})} \right] \right\}. \tag{8}$$

In many cases, calculating the KL divergence directly is intractable. Instead, we optimize an alternative objective that is equivalent to minimizing the KL up to an added constant. We can rewrite Eq. (8) as

$$\begin{aligned} \mathrm{KL}(q(\boldsymbol{\Theta}) \,\|\, p(\boldsymbol{\Theta} \,|\, \mathbf{X})) &= \mathbb{E}_{q(\boldsymbol{\Theta})} \left\{ \ln \left[ \frac{q(\boldsymbol{\Theta})}{p(\boldsymbol{\Theta} \,|\, \mathbf{X})} \right] \right\} \\ &= \mathbb{E}_{q(\boldsymbol{\Theta})} \left[ \ln q(\boldsymbol{\Theta}) \right] - \mathbb{E}_{q(\boldsymbol{\Theta})} \left[ \ln p(\boldsymbol{\Theta} \,|\, \mathbf{X}) \right] \\ &= \mathbb{E}_{q(\boldsymbol{\Theta})} \left[ \ln q(\boldsymbol{\Theta}) \right] - \mathbb{E}_{q(\boldsymbol{\Theta})} \left\{ \ln \left[ \frac{p(\boldsymbol{\Theta}, \mathbf{X})}{p(\mathbf{X})} \right] \right\} \\ &= \mathbb{E}_{q(\boldsymbol{\Theta})} \left[ \ln q(\boldsymbol{\Theta}) \right] - \mathbb{E}_{q(\boldsymbol{\Theta})} \left[ \ln p(\boldsymbol{\Theta}, \mathbf{X}) \right] + \mathbb{E}_{q(\boldsymbol{\Theta})} \left[ \ln p(\mathbf{X}) \right]. \end{aligned} \tag{9}$$

Note that $\mathbb{E}_{q(\boldsymbol{\Theta})} \left[ \ln p(\mathbf{X}) \right] = \ln p(\mathbf{X})$ because the marginal log-likelihood does not depend on the variational parameters. Solving for this marginal log-likelihood in Eq. (9) yields the following

$$\begin{aligned} \ln p(\mathbf{X}) &= \mathrm{KL}(q(\boldsymbol{\Theta}) \,\|\, p(\boldsymbol{\Theta} \,|\, \mathbf{X})) + \mathbb{E}_{q(\boldsymbol{\Theta})} \left[ \ln p(\boldsymbol{\Theta}, \mathbf{X}) \right] - \mathbb{E}_{q(\boldsymbol{\Theta})} \left[ \ln q(\boldsymbol{\Theta}) \right] \\ &= \mathrm{KL}(q(\boldsymbol{\Theta}) \,\|\, p(\boldsymbol{\Theta} \,|\, \mathbf{X})) + \mathcal{L}(\boldsymbol{\Theta}) \end{aligned} \tag{10}$$

where $\mathcal{L}(\boldsymbol{\Theta}) = \mathbb{E}_{q(\boldsymbol{\Theta})} \left[ \ln p(\boldsymbol{\Theta}, \mathbf{X}) \right] - \mathbb{E}_{q(\boldsymbol{\Theta})} \left[ \ln q(\boldsymbol{\Theta}) \right]$ denotes the “evidence lower bound” (ELBO) and is the new objective that we seek to maximize. The first term in $\mathcal{L}(\boldsymbol{\Theta})$ is the expectation of the joint log likelihood for the generative model given by the distributions outlined in Eqs. (1)-(4). This term is

$$\begin{aligned}
\ln p(\boldsymbol{\Theta}, \mathbf{X}) = & \sum_{n=1}^{N}\sum_{j=1}^{J}\sum_{k=1}^{K}[\psi_n = k]\left[-\frac{1}{2}\ln 2\pi - \frac{1}{2}\ln\sigma_j^2 - \frac{1}{2\sigma_j^2}\left(x_{nj} - \nu_j - \mu_{jk}\right)^2\right] \\
& + \sum_{n=1}^{N}\sum_{k=1}^{K}[\psi_n = k]\ln\pi_k + \sum_{j=1}^{J}\sum_{k=1}^{K}\rho_{jk}\left(-\frac{1}{2}\ln 2\pi - \frac{1}{2}\ln\lambda_{jk} - \frac{1}{2}\ln\sigma_j^2 - \frac{1}{2\lambda_{jk}\sigma_j^2}\mu_{jk}^2\right) \\
& + \sum_{j=1}^{J}\sum_{k=1}^{K}[\rho_{jk}\ln\eta + (1-\rho_{jk})]\ln(1-\eta) + \sum_{j=1}^{J}\left[-\frac{1}{2}\ln 2\pi - \frac{1}{2}\ln\tau^2 - \frac{1}{2\tau^2}(v_j - \omega)^2\right] \\
& + \sum_{j=1}^{J}\left[\ln\frac{\xi_2^{\xi_1}}{\Gamma(\xi_1)} - (\xi_1+1)\ln\sigma_j^2 - \frac{\xi_2}{\sigma_j^2}\right] + \sum_{j=1}^{J}\sum_{k=1}^{K}\left[\ln\frac{\kappa_2^{\kappa_1}}{\Gamma(\kappa_1)} - (\kappa_1+1)\ln\lambda_{jk} - \frac{\kappa_2}{\lambda_{jk}}\right] \\
& - \ln\left(\frac{\prod_{k=1}^{K}\Gamma\left[\alpha_0\chi_k\prod_{l=1}^{k-1}(1-\chi_l)\right]}{\Gamma\left[\sum_{k=1}^{K}\alpha_0\chi_k\prod_{l=1}^{k-1}(1-\chi_l)\right]}\right) + \sum_{k=1}^{K}\left[\alpha_0\chi_k\prod_{l=1}^{k-1}(1-\chi_l)\right]\ln\pi_k \\
& + \sum_{k=1}^{K}(\gamma_0 - 1)\ln(1-\chi_k) - K\ln\left(\frac{\Gamma(1)\Gamma(\gamma_0)}{\Gamma(1+\gamma_0)}\right) - \ln\left(\frac{\Gamma(\varphi_1)\Gamma(\varphi_2)}{\Gamma(\varphi_1+\varphi_2)}\right)
\end{aligned} \tag{11}$$

where $\Gamma(\cdot)$ is the gamma function. The second term in $\mathcal{L}(\boldsymbol{\Theta})$ is the variational approximation of our generative model which must be selected. Our choices for the variational distribution will greatly impact our optimization procedure. To that end, we restrict $q(\boldsymbol{\Theta}; \mathbf{Z})$ to be in a form that is easily factorized where $\mathbf{Z} = \{\boldsymbol{r}, \boldsymbol{y}, \boldsymbol{f}, \boldsymbol{t}^2, \boldsymbol{m}, \boldsymbol{s}^2, \boldsymbol{u}, \boldsymbol{v}, \boldsymbol{a}, \boldsymbol{b}, h_1, h_2, \boldsymbol{d}, \boldsymbol{g}_1, \boldsymbol{g}_2\}$ is the collection of the free parameters we will update during model training. We specifically choose the following variational distributions for our approximating family to be in the same family as the prior they are approximating.

First, we choose a Gaussian distribution to approximate the global mean expression for each gene,

$$q(\nu_j; f_j, t_j^2) = \mathcal{N}(\nu_j \mid f_j, t_j^2). \tag{12}$$

Next, following previous work[5,25,26], we approximate the joint distribution of $\mu_{jk}$ and $\rho_{jk}$ with

$$q(\mu_{jk}, \rho_{jk}; y_{jk}, m_{jk}, s_{jk}^2) = \begin{cases} y_{jk}\,\mathcal{N}(\mu_{jk} \mid m_{jk}, s_{jk}^2) & \text{if } \rho_{jk} = 1; \\ 0 & \text{otherwise.} \end{cases} \tag{13}$$

To approximate the probability that the latent indicator variables $\rho_{jk} = 1$, we choose a Bernoulli where

$$q[\rho_{jk} = 1] = y_{jk} \tag{14}$$

with $y_{jk}$ being the probability that the expression of a gene deviates from the baseline across all cells. To approximate the slab scaling factor $\lambda_{jk}$ and the global gene-specific variance $\sigma_j^2$, we choose the following inverse-gamma distribution specifications

$$q(\lambda_{jk}; u_{jk}, v_{jk}) = \text{Inv-Gamma}(\lambda_{jk} \mid u_{jk}, v_{jk}), \tag{15}$$

$$q(\sigma_j^2; a_j, b_j) = \text{Inv-Gamma}(\sigma_j^2 \mid a_j, b_j). \tag{16}$$

To approximate $\eta$, we choose a Beta distribution which is specified as the following

$$q(\eta; h_1, h_2) = \text{Beta}(\eta \mid h_1, h_2). \tag{17}$$

For the second half of the model, we approximate the cluster indicator variables $\psi_n$ with a categorical distribution such that

$$q(\psi_n; \boldsymbol{r}_n) = \text{Cat}(\boldsymbol{r}_n), \quad \sum_k r_{nk} = 1 \tag{18}$$

where $\boldsymbol{r}_n = [r_{n1}, \ldots, r_{nK}]$ is a $K$-dimensional vector containing the probability that the $n$-th cell belongs to a given $k$-th cluster. To approximate $\boldsymbol{\pi}$, we choose a Dirichlet distribution

$$q(\boldsymbol{\pi}; \boldsymbol{d}) = \text{Dir}(\boldsymbol{d}). \tag{19}$$

Finally, previous work using variational inference with hierarchical Dirichlet proccess (HDP) priors [27,28] focused on using point estimate approximations to emulate the stick-breaking procedure of the top-level Dirichlet process. However, recent work [11] has shown that this approximation leads to the creation of many small and redundant clusters. Instead, it has been suggested to use a proper Beta distribution over each stick-breaking parameter. Following this finding, we approximate $\chi_k$ with the following

$$q(\chi_k; g_{1k}, g_{2k}) = \text{Beta}(g_{1k}g_{2k}, (1-g_{1k})g_{2k}) \tag{20}$$

where $0 < g_{1k} < 1$ defines the average length of the $k$-th stick-breaking length and $0 < g_{2k}$ controls the variance of the length of $k$-th stick-breaking parameter.

Note that, for simplicity, we use the Greek characters to denote the parameters from the true posterior distribution and the Latin alphabet to denote the free parameters of the variational posterior. Below, we will find closed-form updates for all the free parameters, except for $g_{1k}$ and $g_{2k}$ which we will solve using stochastic gradient descent (SGD)[29]. Given our fully specified model in Eqs. (1)-(4) and the variational approximating family of distributions in Eqs. (12)-(20), the joint ELBO using all the data is given as

$$\begin{aligned}
\mathcal{L}(\boldsymbol{\Theta}) &= \mathbb{E}_{q(\boldsymbol{\Theta})}\left[\ln p(\boldsymbol{\Theta}, \mathbf{X})\right] - \mathbb{E}_{q(\boldsymbol{\Theta})}\left[\ln q(\boldsymbol{\Theta})\right] \qquad (21)\\
&= \sum_{j=1}^{J}\sum_{k=1}^{K}\left\{-\frac{N_k}{2}\ln 2\pi - \frac{N_k}{2}\left[\ln b_j - \boldsymbol{\Psi}(a_j)\right] - \frac{a_j}{2b_j}\left[\widehat{x}_{jk}^2 - 2\widehat{x}_{jk}\mathbb{E}_{q(\boldsymbol{\Theta})}[\rho_{jk}\mu_{jk} + \nu_j] + N_k\mathbb{E}_{q(\boldsymbol{\Theta})}[(\rho_{jk}\mu_{jk} + \nu_j)^2]\right]\right\}\\
&+ \sum_{j=1}^{J}\sum_{k=1}^{K}\left\{-\left(\kappa_1 + \frac{y_{jk}}{2} - u_{jk}\right)\left[\ln v_{jk} - \boldsymbol{\Psi}(u_{jk})\right] - \kappa_2\left(\frac{u_{jk}}{v_{jk}}\right)\right\}\\
&+ \sum_{j=1}^{J}\sum_{k=1}^{K}\left\{-\frac{y_{jk}}{2}\left(\frac{u_{jk}}{v_{jk}}\right)\left(\frac{a_j}{b_j}\right)\mathbb{E}_{q(\boldsymbol{\Theta})}[\mu_{jk}^2] + \frac{y_{jk}}{2} + \frac{y_{jk}}{2}\ln s_{jk}^2\right\}\\
&+ \sum_{j=1}^{J}\left\{-\ln\left(\frac{b_j^{a_j}}{\Gamma(a_j)}\right) + a_j\right\} + \sum_{j=1}^{J}\sum_{k=1}^{K}\left\{-\ln\left(\frac{v_{jk}^{u_{jk}}}{\Gamma(u_{jk})}\right) + u_{jk}\right\}\\
&+ \sum_{j=1}^{J}\left\{-\left(\xi_1 + \frac{1}{2}\sum_{k=1}^{K}y_{jk} - a_j\right)\left[\ln b_j - \boldsymbol{\Psi}(a_j)\right] - \xi_2\left(\frac{a_j}{b_j}\right) - \frac{1}{2\tau^2}\left(\mathbb{E}_{q(\boldsymbol{\Theta})}[\nu_j^2] - 2\omega\mathbb{E}_{q(\boldsymbol{\Theta})}[\nu_j]\right) + \frac{1}{2}\ln t_j^2\right\}\\
&+ \left(\varphi_1 - h_1 + \sum_{k=1}^{K}\sum_{j=1}^{J}y_{jk}\right)\left[\boldsymbol{\Psi}\left(h_1\right) - \boldsymbol{\Psi}\left(h_1 + h_2\right)\right] + \left(\varphi_2 - h_2 + \sum_{k=1}^{K}\sum_{j=1}^{J}(1 - y_{jk})\right)\left[\boldsymbol{\Psi}\left(h_2\right) - \boldsymbol{\Psi}\left(h_1 + h_2\right)\right]\\
&- \ln\left(\frac{\Gamma(\varphi_1)\Gamma(\varphi_2)}{\Gamma(\varphi_1 + \varphi_2)}\right) + \ln\left(\frac{\Gamma(h_1)\Gamma(h_2)}{\Gamma(h_1 + h_2)}\right) + KJ\ln\left(\frac{\kappa_2^{\kappa_1}}{\Gamma(\kappa_1)}\right) + J\ln\left(\frac{\xi_2^{\xi_1}}{\Gamma(\xi_1)}\right) - \frac{J}{2}\ln\tau^2 - \frac{J\omega^2}{2\tau^2} + \frac{J}{2}\\
&+ \sum_{j=1}^{J}\sum_{k=1}^{K}\left\{-y_{jk}\ln y_{jk} + (1 - y_{jk})\ln(1 - y_{jk})\right\} + \sum_{n=1}^{N}\sum_{k=1}^{K+1}\left\{r_{nk}\left[\boldsymbol{\Psi}\left(d_k\right) - \boldsymbol{\Psi}\left(\sum_{k'=1}^{K}d_{k'}\right)\right] - \ln\left(\frac{\prod_{k=1}^{K}\Gamma\left(d_k\right)}{\Gamma\left(\sum_{k=1}^{K}d_k\right)}\right)\right\}\\
&- \sum_{n=1}^{N}\sum_{k=1}^{K}r_{nk}\ln r_{nk} + \sum_{k=1}^{K}\left[\boldsymbol{\Psi}(g_{1k}g_{2k}) - \boldsymbol{\Psi}(g_{2k})\right] + \sum_{k=1}^{K}(K + 1 - k)\left[\boldsymbol{\Psi}((1 - g_{1k})g_{2k}) - \boldsymbol{\Psi}(g_{2k})\right]
\end{aligned}$$

$$+ K \ln \alpha_0 + \sum_{k=1}^{K+1} \left( \alpha_0 g_{1k} \prod_{l=1}^{k-1} (1 - g_{1\ell}) - d_k \right) \left[ \mathbf{\Psi} \left( d_k \right) - \mathbf{\Psi} \left( \sum_{k'=1}^{K} d_{k'} \right) \right]$$

$$+ \sum_{k=1}^{K} (\gamma_0 - (1 - g_{1k}) g_{2k}) \left[ \mathbf{\Psi}((1 - g_{1k}) g_{2k}) - \mathbf{\Psi}(g_{2k}) \right] + (1 - g_{1k} g_{2k}) \left[ \mathbf{\Psi}(g_{1k} g_{2k}) - \mathbf{\Psi}(g_{2k}) \right]$$

$$- K \ln \left( \frac{\Gamma(1)\Gamma(\gamma_0)}{\Gamma(1 + \gamma_0)} \right) + \sum_{k=1}^{K} \ln \left( \frac{\Gamma(g_{1k} g_{2k}) \Gamma\left((1 - g_{1k}) g_{2k}\right)}{\Gamma(g_{2k})} \right)$$

where, according to the chosen variational approximating distributions, we have the following:

$$\mathbb{E}_{q(\mathbf{\Theta})}[\rho_{jk}\mu_{jk}] = y_{jk} m_{jk} \tag{22}$$

$$\mathbb{E}_{q(\mathbf{\Theta})}[\rho_{jk}\mu_{jk}^2] = y_{jk}(m_{jk}^2 + s_{jk}^2) \tag{23}$$

$$\mathbb{E}_{q(\mathbf{\Theta})}[\nu_j^2] = f_j^2 + t_j^2 \tag{24}$$

$$\mathbb{E}_{q(\mathbf{\Theta})}[(\mu_{jk} + \nu_j)^2] = 2 y_{jk} m_{jk} f_j + y_{jk}(m_{jk}^2 + s_{jk}^2) + (f_j^2 + t_j^2). \tag{25}$$

Furthermore, in the expression for the ELBO above, we have $N_k = \sum_{n=1}^{N} r_{nk}$ which is the variational estimate of the number of cells assigned to a given cluster, $\widehat{x}_{jk} = \sum_{i=1}^{N} r_{nk} x_{nj}$ is the weighted expression of the $j$-th gene in the $k$-th cluster after taking into account all cell assignments, and $\mathbf{\Psi}(\cdot)$ is the digamma function.

It is worth noting that we follow the truncated stick-breaking approximation approach proposed by Blei and Jordan[1] and use a finite mixture with a fixed number $K$ of normal components to estimate the number of clusters. Even though we use a finite mixture as an approximation to the posterior distribution, our likelihood still consists of a mixture of infinitely many normal distributions[1,10]. In practice, we generally set $K$ to be a very large number (i.e., $K = 25$ components for the results in the main text)—this is because placing a proper beta distribution as the variational approximating distribution for each $\chi_k$ encourages the penalization of empty clusters[11]. As a result, only some subset of clusters $K^* \leq K$ will be used in the final model.

We now describe the expectation and maximization steps of the variational EM algorithm, formally known as coordinate ascent variational inference. As stated above, the goal is to optimize the evidence lower bound (ELBO) by approximating the intractable posterior distribution with a more tractable family of distributions. One common approach to deriving variational updates is to take the partial derivative of the ELBO with respect to each parameter, set this derivative to zero, and solve for the optimal values.

We take an equivalent approach that leverages the fact that each variational distribution belongs to the exponential family and is, therefore, conditionally conjugate. This property simplifies the derivation of each update by allowing us to compute the expectation of the full log joint distribution given in Eq. (11) with respect to all latent parameters except the parameter of interest.

Recall that $\boldsymbol{\Theta} = \{\boldsymbol{\nu}, \boldsymbol{\mu}, \boldsymbol{\sigma}^2, \boldsymbol{\lambda}, \boldsymbol{\rho}, \eta, \boldsymbol{\pi}, \boldsymbol{\psi}, \boldsymbol{\chi}\}$ denotes the full set of parameters for which we make a variational approximation with each $\ell$-th element being represented by $\boldsymbol{\theta}_\ell \in \boldsymbol{\Theta}$.The optimal variational distribution for each parameter is then given by the following

$$q^*(\boldsymbol{\theta}_\ell) \propto \exp\left\{\mathbb{E}_{q(\boldsymbol{\Theta}\setminus\boldsymbol{\theta}_\ell)}\left[\ln p(\mathbf{X}, \boldsymbol{\Theta})\right]\right\}, \tag{26}$$

where the expectation is taken with respect to the variational distributions of all other parameters in $\boldsymbol{\Theta}$ except the $\ell$-th. This approach guarantees $(i)$ the ELBO in Eq. (21) is maximized after each step and $(ii)$ that each variational factor remains in the same distributional family as its prior, resulting in closed-form updates that resemble posterior distributions in Bayesian inference. Below, we outline the closed-form variational updates for each parameter in our model.

#### 2.1 Step #1: Variational E-Step

In the E-step of the algorithm, we use the current variational estimates of $\{\boldsymbol{\nu}, \boldsymbol{\mu}, \boldsymbol{\sigma}^2, \boldsymbol{\lambda}, \eta, \boldsymbol{\chi}\}$ to estimate the the approximate inclusion probability that the $j$-th gene in the $k$-th cluster $(y_{jk})$ and the approximate probability that the $n$-th cell is in $k$-th cluster $(r_{nk})$.

##### 2.1.1 Variational Update to $\boldsymbol{y}_{jk}$

The probability $y_{jk}$ represents the (approximate) probability that the $j$-th gene is a driver of the phenotypic profile for cells assigned in the $k$-th cluster. Using the joint log-likelihood in Eq. (11) and the update strategy detailed in Eq. (26), we get the following

$$\begin{aligned} q^*(\boldsymbol{\rho}) \propto \exp\Bigg\{\mathbb{E}_{q(\boldsymbol{\Theta}\setminus\boldsymbol{\rho})}\Bigg[\sum_{j=1}^{J}\sum_{k=1}^{K}\rho_{jk}\left(-\frac{1}{2}\ln 2\pi - \frac{1}{2}\ln\lambda_{jk} - \frac{1}{2}\ln\sigma_j^2 - \left(\frac{1}{2\lambda_{jk}\sigma_j^2}\right)\mu_{jk}^2 + \ln\frac{\eta}{1-\eta}\right) \\ + \sum_{n=1}^{N}[\psi_n = k]\left[-\frac{1}{2\sigma_j^2}\left(x_{nj} - \nu_j - \rho_{jk}\mu_{jk}\right)^2\right] + \text{Constant}\Bigg]\Bigg\} \end{aligned}$$

where the constant term contains all terms in Eq. (11) that do not involve $\rho_{jk}$ (from hereafter, we will omit these terms for simplicity). We can distribute the expectation operator throughout the sum and replace the terms involving the other parameters in the model with closed-form expressions. Namely, we let $\mathbb{E}_q[\ln \lambda_{jk}] = \ln v_{jk} - \boldsymbol{\Psi}(u_{jk})$; $\mathbb{E}_q\left[\ln \sigma_j^2\right] = \ln b_j - \boldsymbol{\Psi}(a_j)$; $\mathbb{E}_q[1/\lambda_{jk}] = u_{jk}/v_{jk}$; $\mathbb{E}_q\left[1/\sigma_j^2\right] = a_j/b_j$; $\mathbb{E}_q\left[\mu_{jk}^2\right] = m_{jk}^2 + s_{jk}^2$, $\mathbb{E}_q[\ln \eta/(1-\eta)] = \boldsymbol{\Psi}(h_1) - \boldsymbol{\Psi}(h_2)$, and $\mathbb{E}_q[\psi_n = k] = r_{nk}$. The expectation of $\mathbb{E}_q\left[(x_{nj} - \nu_j - \rho_{jk}\mu_{jk})^2\right]$ can be rearranged such that terms that do not involve $\rho_{jk}$ are added to the Constant term. The remaining terms are $-2x_{nj}\rho_{jk}m_{jk} + 2\rho_{jk}m_{jk}f_j + \rho_{jk}(m_{jk}^2 + s_{jk}^2)$. Substituting these closed form expressions back into the update equation yields

$$
\begin{aligned}
q^*(\boldsymbol{\rho}) \propto \exp\Bigg\{ & \sum_{j=1}^{J}\sum_{k=1}^{K} \rho_{jk}\left(-\frac{1}{2}\ln 2\pi - \frac{1}{2}[\ln v_{jk} - \boldsymbol{\Psi}(u_{jk})] - \frac{1}{2}[\ln b_j - \boldsymbol{\Psi}(a_j)] - \frac{u_{jk}a_j}{2v_{jk}b_j}(m_{jk}^2 + s_{jk}^2) + \boldsymbol{\Psi}(h_1) - \boldsymbol{\Psi}(h_2)\right) \\
& + \sum_{n=1}^{N} r_{nk}\left[-\frac{a_j}{2b_j}\left(-2x_{nj}\rho_{jk}m_{jk} + 2\rho_{jk}m_{jk}f_j + \rho_{jk}(m_{jk}^2 + s_{jk}^2)\right)\right]\Bigg\}.
\end{aligned}
$$

Next, using some algebra to rearrange terms and using the fact that $N_k = \sum_{n=1}^{N} r_{nk}$ and $\widehat{x}_{jk} = \sum_{i=1}^{N} r_{nk}x_{nj}$, we get the following expression

$$
\begin{aligned}
q^*(\boldsymbol{\rho}) \propto \exp\Bigg\{ & \sum_{j=1}^{J}\sum_{k=1}^{K} \rho_{jk}\Bigg(-\frac{1}{2}\ln 2\pi - \frac{1}{2}[\ln v_{jk} - \boldsymbol{\Psi}(u_{jk})] - \frac{1}{2}[\ln b_j - \boldsymbol{\Psi}(a_j)] - \frac{u_{jk}a_j}{2v_{jk}b_j}(m_{jk}^2 + s_{jk}^2) + \boldsymbol{\Psi}(h_1) - \boldsymbol{\Psi}(h_2) \\
& + \left(\frac{a_j}{b_j}\right)\widehat{x}_{jk}m_{jk} - N_k\left(\frac{a_j}{b_j}\right)m_{jk}f_j - \frac{a_j}{2b_j}N_k(m_{jk}^2 + s_{jk}^2)\Bigg)\Bigg\} \\
\propto \exp\Bigg\{ & \sum_{j=1}^{J}\sum_{k=1}^{K} \rho_{jk}\Bigg(-\frac{1}{2}\ln 2\pi - \frac{1}{2}[\ln v_{jk} - \boldsymbol{\Psi}(u_{jk})] - \frac{1}{2}[\ln b_j - \boldsymbol{\Psi}(a_j)] + \boldsymbol{\Psi}(h_1) - \boldsymbol{\Psi}(h_2) \\
& - \frac{a_j}{2b_j}\left(\frac{u_{jk}}{v_{jk}} + N_k\right)(m_{jk}^2 + s_{jk}^2) + \frac{a_j}{b_j}m_{jk}[\widehat{x}_{jk} - N_k f_j]\Bigg)\Bigg\}.
\end{aligned}
$$

In Section 2.2.1 under Eq. (29), we will show that

$$
\left(\frac{a_j}{b_j}\right)\widehat{x}_{jk} - N_k f_j = m_{jk}\frac{a_j}{b_j}\left(\frac{u_{jk}}{v_{jk}} + N_k\right) = \frac{m_{jk}}{s_{jk}^2}
$$

which is the reciprocal index of dispersion for the variational approximation of $\mu_{jk}$ (see Cox and Lewis [30]). Likewise, we will also show that $(a_j/b_j)[(u_{jk}/v_{jk} + N_k)] = 1/s_{jk}^2$ which is the precision of variational approximate distribution for $\mu_{jk}$. Substituting both of these values and simplifying yields

$$q^*(\boldsymbol{\rho}) \propto \exp\Bigg\{ \sum_{j=1}^{J}\sum_{k=1}^{K} \rho_{jk} \Bigg( -\frac{1}{2}\ln 2\pi - \frac{1}{2}[\ln v_{jk} - \boldsymbol{\Psi}(u_{jk})] - \frac{1}{2}[\ln b_j - \boldsymbol{\Psi}(a_j)] + \boldsymbol{\Psi}(h_1) - \boldsymbol{\Psi}(h_2)$$

$$- \frac{1}{2s_{jk}^2}(m_{jk}^2 + s_{jk}^2) + \frac{m_{jk}^2}{s_{jk}^2} \Bigg) \Bigg\}$$

$$\propto \exp\Bigg\{ \sum_{j=1}^{J}\sum_{k=1}^{K} \rho_{jk} \Bigg( -\frac{1}{2}\ln 2\pi - \frac{1}{2}[\ln v_{jk} - \boldsymbol{\Psi}(u_{jk})] - \frac{1}{2}[\ln b_j - \boldsymbol{\Psi}(a_j)] + \boldsymbol{\Psi}(h_1) - \boldsymbol{\Psi}(h_2) + \frac{m_{jk}^2}{2s_{jk}^2} - \frac{1}{2} \Bigg) \Bigg\}$$

Note that $q^*(\boldsymbol{\rho})$ takes the form of a Bernoulli distribution. To make this clearer, let $\ln \widetilde{y}_{jk} = -(1/2)\ln 2\pi - (1/2)[\ln v_{jk} - \boldsymbol{\Psi}(u_{jk})] - (1/2)[\ln b_j - \boldsymbol{\Psi}(a_j)] + \boldsymbol{\Psi}(h_1) - \boldsymbol{\Psi}(h_2) + m_{jk}^2/2s_{jk}^2 - 1/2$ be an un-normalized probability. Then, we get the following relationship

$$q^*(\boldsymbol{\rho}) \propto \exp\left\{ \sum_{j=1}^{J}\sum_{k=1}^{K} \ln \widetilde{y}_{jk} \right\}$$

$$\propto \prod_{k=1}^{K}\prod_{j=1}^{J} \text{Bernoulli}\left(\text{Sigmoid}\left\{ \ln \widetilde{y}_{jk} \right\}\right)$$

$$\propto \prod_{k=1}^{K}\prod_{j=1}^{J} \text{Bernoulli}\left(y_{jk}\right)$$

where, by applying a sigmoid (i.e., the standard logistic function), we arrive at the update

$$y_{jk} = \text{Sigmoid}\left\{ -\frac{1}{2}\ln 2\pi - \frac{1}{2}[\ln v_{jk} - \boldsymbol{\Psi}(u_{jk})] - \frac{1}{2}[\ln b_j - \boldsymbol{\Psi}(a_j)] + \boldsymbol{\Psi}(h_1) - \boldsymbol{\Psi}(h_2) + \frac{1}{2}\frac{m_{jk}^2}{s_{jk}^2} - \frac{1}{2} \right\}. \tag{27}$$

Here, $y_{jk} \approx \Pr[\rho_{jk} = 1 \mid \mathbf{X}, \boldsymbol{\Theta}, \eta]$ denotes the approximate posterior inclusion probability (PIP) for each gene. Note that this updated form for the probabilities can be effectively seen as the prior log odds $\eta/(1-\eta)$ being updated by the log Bayes factor for the alternative hypothesis that the $j$-th gene is playing a role in driving the assignment of cells to the $k$-th cluster[5]. Taken together, $\ln(y_{jk}/(1-y_{jk}))$ represents the posterior odds of choosing the alternative hypothesis over the null hypothesis (i.e., that the $j$-th gene does not affect the formation of a given cluster).

##### 2.1.2 Variational Update to $\boldsymbol{r}_{nk}$

Recall that $r_{nk}$ is the (approximate) probability that the $n$-th cell belongs to the $k$-th cluster. Similar to what we did in Section 2.1.1, we use the joint log-likelihood in Eq. (11) and the optimization framework in Eq. (26). By take the expectation with respect to all terms that involve the variable $\psi_n$, we get the following simplified expression

$$q^*(\boldsymbol{\psi}) \propto \exp\left\{\mathbb{E}_{q(\boldsymbol{\Theta}\setminus\boldsymbol{\psi})}\left[\sum_{n=1}^{N}\sum_{k=1}^{K}[\psi_n = k]\left(\ln \pi_k - \frac{J}{2}\ln 2\pi - \frac{1}{2}\sum_{j=1}^{J}\ln\sigma_j^2 - \sum_{j=1}^{J}\frac{1}{2\sigma_j^2}\left(x_{nj} - \nu_j - \rho_{jk}\mu_{jk}\right)^2\right)\right]\right\}$$

where we omit all the terms Eq. (11) that do not involve $\psi_n$. Once again, we can distribute the expectation operator and replace the terms involving the other parameters in the model with closed-form expressions. Namely, $\mathbb{E}_q\left[\ln \pi_k\right] = \boldsymbol{\Psi}(d_k) - \boldsymbol{\Psi}\left(\sum_k d_k\right)$; $\mathbb{E}_q\left[\ln\sigma_j^2\right] = \ln b_j - \boldsymbol{\Psi}(a_j)$, and $\mathbb{E}_q\left[1/\sigma_j^2\right] = a_j/b_j$. The form of the final expectation term is $\mathbb{E}_q\left[\left(x_{nj} - \nu_j - \rho_{jk}\mu_{jk}\right)^2\right] = x_{nj}^2 - 2x_{nj}(y_{jk}m_{jk} + f_j) + 2y_{jk}m_{jk}f_j + y_{jk}(m_{jk}^2 + s_{jk}^2) + (f_j^2 + t_j^2)$. Substituting these terms back into the expression, we get an un-normalized probability

$$\begin{aligned}
q^*(\boldsymbol{\psi}) \propto \exp&\left\{\sum_{n=1}^{N}\sum_{k=1}^{K}[\psi_n = k]\left(\left[\boldsymbol{\Psi}(d_k) - \boldsymbol{\Psi}\left(\sum_k d_k\right)\right] - \frac{J}{2}\ln 2\pi - \frac{1}{2}\sum_{j=1}^{J}[\ln b_j - \boldsymbol{\Psi}(a_j)]\right.\right.\\
&\left.\left. - \sum_{j=1}^{J}\frac{a_j}{2b_j}\left(x_{nj}^2 - 2x_{nj}(y_{jk}m_{jk} + f_j) + 2y_{jk}m_{jk}f_j + y_{jk}(m_{jk}^2 + s_{jk}^2) + (f_j^2 + t_j^2)\right)\right]\right\}\\
\propto \exp&\left\{\sum_{n=1}^{N}\sum_{k=1}^{K}[\psi_n = k)\ln\widetilde{r}_{nk}\right\}
\end{aligned}$$

where $\widetilde{r}_{nk}$ takes the form

$$\begin{aligned}
\widetilde{r}_{nk} = &\left[\boldsymbol{\Psi}(d_k) - \boldsymbol{\Psi}\left(\sum_k d_k\right)\right] - \frac{J}{2}\ln 2\pi - \frac{1}{2}\sum_{j=1}^{J}[\ln b_j - \boldsymbol{\Psi}(a_j)] - \sum_{j=1}^{J}\frac{a_j}{2b_j}\left(x_{nj}^2 - 2x_{nj}(y_{jk}m_{jk} + f_j)\right.\\
&\left.+2y_{jk}m_{jk}f_j + y_{jk}(m_{jk}^2 + s_{jk}^2) + (f_j^2 + t_j^2)\right).
\end{aligned}$$

Here, it is clear that $q^*(\boldsymbol{\psi})$ takes the form of a categorical distribution where

$$q^*(\boldsymbol{\psi}) \propto \exp\left\{\sum_{j=1}^{N}\sum_{k=1}^{K}\ln\widetilde{r}_{nk}\right\}$$

$$\propto \prod_{n=1}^{N} \prod_{k=1}^{K} \text{Categorical}\left(\text{Softmax}\left\{ \ln \widetilde{r}_{nk} \right\}\right)$$

$$\propto \prod_{n=1}^{N} \prod_{k=1}^{K} \text{Categorical}\left(r_{nk}\right).$$

To convert this value to the unit scale, we take the softmax which normalizes it with respect to the other clusters. This yields the following closed-form update

$$r_{nk} = \text{Softmax}(\widetilde{r}_{nk}) = \frac{\exp\{\widetilde{r}_{nk}\}}{\sum_{k'=1}^{K} \exp\{\widetilde{r}_{nk'}\}}, \tag{28}$$

where $r_{nk} \approx \Pr[\psi_n = k \,|\, \mathbf{X}, \mathbf{\Theta}]$. Importantly, this update provides us with insight into how the model clusters different cells. These cell-specific updates are of the same form as posterior probabilities in traditional nonparametric models. Note, however, that our method makes an important modification: instead of assuming that each gene has an equal contribution to a cell clustering assignment, the contribution of each gene is weighted by its importance in that cluster. This is captured by the approximate posterior inclusion probability for each gene given by $y_{jk}$ in Eq. (27).

#### 2.2 Step #2: Variational M-Step

In the M-step of the algorithm, we update the other parameters conditioned on the probabilities learned in the E-Step in Section 2.1. The intuition behind this approach is simple: cells and genes that are more likely to arise in the $k$-th cluster will have more influence on the parameter update of $k$-th cluster. We assume that the likelihood that the $n$-th cell contributes to updating the $k$-th cluster parameters is proportional to the cluster membership probability $r_{nk}$ that we calculated in Section 2.1.2. Similarly, we update $j$-th gene's parameters in the $k$-th cluster in proportion to its cluster inclusion probability $y_{jk}$ that we calculated in Section 2.1.1. We still follow the previously described procedure; namely, for a given variable in $\boldsymbol{\theta}_\ell \in \{\boldsymbol{\mu}, \boldsymbol{\nu}, \boldsymbol{\sigma}^2, \boldsymbol{\lambda}, \boldsymbol{\pi}, \eta, \boldsymbol{\chi}\}$, $(i)$ we use the joint log-likehood in Eq. (11) and the optimal update procedure of Eq. (26), $(ii)$ apply the expectation operator to all other variables that interact with $\boldsymbol{\theta}_\ell$, and $(iii)$ simplify the expression, keeping only the terms that depend on $\boldsymbol{\theta}_\ell$. The expression that results from this procedure is the variational updates that maximize the lower bound ELBO in Eq. (21). As we repeat this procedure, we arrive at the closed-form updates detailed below.

##### 2.2.1 Variational Update to $\boldsymbol{m}_{jk}$ and $\boldsymbol{s}_{jk}^2$

Recall from Eq. (13) that $m_{jk}$ and $s_{jk}^2$ are the respective mean and variance of the variational distribution that approximates $\mu_{jk}$. To find the variational updates for these terms, we let $\boldsymbol{\theta}_\ell = \boldsymbol{\mu}$ and find the following expression for the algorithmic updates

$$q^*(\boldsymbol{\mu}) \propto \exp\left\{\mathbb{E}_{q(\boldsymbol{\Theta}\setminus\boldsymbol{\mu})}\left[\sum_{j=1}^{J}\sum_{k=1}^{K} -\left(\frac{1}{2\lambda_{jk}\sigma_{jk}^2}\right)\rho_{jk}\mu_{jk}^2 + \sum_{n=1}^{N}[\psi_n = k]\left[-\frac{1}{2\sigma_j^2}\left(x_{nj} - \nu_j - \rho_{jk}\mu_{jk}\right)^2\right)\right]\right\}.$$

Distributing the expectation we get the following closed-form relationships: $\mathbb{E}_q[\rho_{jk}] = y_{jk}$; $\mathbb{E}_q[1/\lambda_{jk}] = u_{jk}/v_{jk}$; $\mathbb{E}_q\left[1/\sigma_j^2\right] = a_j/b_j$; and $\mathbb{E}_q[\psi_n = k] = r_{nk}$. The result of $\mathbb{E}_q\left[\left(x_{nj} - \nu_j - \rho_{jk}\mu_{jk}\right)^2\right]$ can be rearranged such that many of the terms do not involve $\mu_{jk}$ and, thus, are dropped—the remaining terms are $-2x_{nj}y_{jk}\mu_{jk} + 2y_{jk}\mu_{jk}f_j + y_{jk}\mu_{jk}^2$. Substituting these back into the equation above yields the following

$$q^*(\boldsymbol{\mu}) \propto \exp\left\{\sum_{j=1}^{J}\sum_{k=1}^{K} -\left(\frac{a_j u_{jk}}{2b_j v_{jk}}\right) y_{jk}\mu_{jk}^2 + \sum_{n=1}^{N} r_{nk}\left[-\frac{a_j}{2b_j}\left(-2x_{nj}y_{jk}\mu_{jk} + 2y_{jk}\mu_{jk}f_j + y_{jk}\mu_{jk}^2\right)\right]\right\}.$$

Next, we can use the fact that $N_k = \sum_{n=1}^{N} r_{nk}$ and $\widehat{x}_{jk} = \sum_{i=1}^{N} r_{nk}x_{nj}$ and combine like-terms to get

$$q^*(\boldsymbol{\mu}) \propto \exp\left\{\sum_{j=1}^{J}\sum_{k=1}^{K} -\frac{a_j}{2b_j}\left(\frac{u_{jk}}{v_{jk}} + N_k\right) y_{jk}\mu_{jk}^2 + \frac{a_j}{b_j}\left[\widehat{x}_{jk} - f_j N_k\right] y_{jk}\mu_{jk}\right\}.$$

This results in a Gaussian quadratic, which becomes clear when we complete the square and simplify

$$\begin{aligned}
q^*(\boldsymbol{\mu}) &\propto \exp\left\{\sum_{j=1}^{J}\sum_{k=1}^{K} y_{jk}\left[-\frac{a_j}{2b_j}\left(\frac{u_{jk}}{v_{jk}} + N_k\right)\left(\mu_{jk} - \frac{\widehat{x}_{jk} - f_j N_k}{u_{jk}/v_{jk} + N_k}\right)^2\right]\right\} \\
&\propto \exp\left\{\sum_{j=1}^{J}\sum_{k=1}^{K} y_{jk}\ln\mathcal{N}\left(\frac{\widehat{x}_{jk} - f_j N_k}{u_{jk}/v_{jk} + N_k}, \left[\frac{a_j}{b_j}\left(\frac{u_{jk}}{v_{jk}} + N_k\right)\right]^{-1}\right)\right\} \\
&\propto \prod_{k=1}^{K}\prod_{j=1}^{J}\left[\mathcal{N}\left(\frac{\widehat{x}_{jk} - f_j N_k}{u_{jk}/v_{jk} + N_k}, \left[\frac{a_j}{b_j}\left(\frac{u_{jk}}{v_{jk}} + N_k\right)\right]^{-1}\right)\right]^{y_{jk}} \\
&\propto \prod_{k=1}^{K}\prod_{j=1}^{J}\left[\mathcal{N}\left(m_{jk}, s_{jk}^2\right)\right]^{y_{jk}}.
\end{aligned}$$

This expression implies that the variational updates for $m_{jk}$ and $s_{jk}^2$ are the following

$$m_{jk} = \frac{\widehat{x}_{jk} - f_j N_k}{u_{jk}/v_{jk} + N_k}, \quad s_{jk}^2 = \left[\frac{a_j}{b_j}\left(\frac{u_{jk}}{v_{jk}} + N_k\right)\right]^{-1} \tag{29}$$

From Eq. (29), we can derive the equivalence relationships $(a_j/b_j)(u_{jk}/v_{jk} + N_k) = 1/s_{jk}^2$ and $(a_j/b_j)(\widehat{x}_{jk} - f_j N_k) = (a_j/b_j)(u_{jk}/v_{jk} + N_k)\, m_{jk} = m_{jk}/s_{jk}^2$ which can used to simplify other derivations (e.g., see Section 2.1.1). We would like to point out that the form for $s_{jk}^2$ is the posterior variance of the additive effect of $\mu_{jk}$ for a single-variable linear model. Furthermore, $m_{jk}$ can be interpreted as the residual expression of the $j$-th gene in the $k$-th cluster when accounting for the mean baseline expression of the $j$-th gene, $f_j$. Both $m_{jk}$ and $s_{jk}^2$ play an important role in both $(i)$ determining what clusters can be detected and $(ii)$ the types of phenotypic shifts within a cluster that can be detected by the model. For example, in Eq. (27), the term $m_{jk}^2/s_{jk}^2$ is the same as the classical signal to noise ratio statistic used in information theory. This value is minimized when $m_{jk} \to 0$ or $s_{jk}^2 \to \infty$. Because of the structure of the updates in Eq. (29), $m_{jk} \to 0$ only when the $N_k f_j$ is close to $\widehat{x}_{jk}$, the weighted cluster-wise expression of the $j$-th gene in the $k$-th cluster $j$. This makes the accuracy of $f_j$ paramount to the power of NCLUSION to perform variable selection. Similarly, the estimation of the slab's precision $(u_{jk}/v_{jk})^{-1}$ is of equal importance as it can prevent $s_{jk}^2 \to 0$ which can drive the inclusion probability to 1 and generate excessive false positives. During model development, we found it necessary to allow each gene in each cluster to have its own independent slab distribution in order to prevent $s_{jk}^2 \to 0$.

##### 2.2.2 Variational Update to $\boldsymbol{f}_j$ and $\boldsymbol{t}_j^2$.

Next, to find the variational updates for $f_j$ and $t_j^2$, we let $\boldsymbol{\theta}_\ell = \boldsymbol{\nu}$. This yields the following expression for the update equation

$$q^*(\boldsymbol{\nu}) \propto \exp\left\{\mathbb{E}_{q(\boldsymbol{\Theta}\setminus\boldsymbol{\nu})}\left[\sum_{j=1}^{J} -\frac{1}{2\tau^2}(\nu_j - \omega)^2 + \sum_{n=1}^{N}\sum_{k=1}^{K}[\psi_n = k]\left[-\frac{1}{2\sigma_j^2}\left(x_{nj} - \nu_j - \rho_{jk}\mu_{jk}\right)^2\right]\right]\right\}.$$

Distributing the expectation operator throughout the above equation yields the following closed-form expressions: $\mathbb{E}_q[\rho_{jk}] = y_{jk}$; $\mathbb{E}_q\left[1/\sigma_j^2\right] = a_j/b_j$; and $\mathbb{E}_q[\psi_n = k] = r_{nk}$. Similar to the other updates, $\mathbb{E}_q\left[\left(x_{nj} - \nu_j - \rho_{jk}\mu_{jk}\right)^2\right]$ can be rearranged such that the terms that do not involve $\nu_j$ are ignored, with the remaining terms being $-2x_{nj}\nu_j + 2y_{jk}m_{jk}\nu_j + \nu_j^2$. Substituting these expectations back into the optimization function yields

$$q^*(\boldsymbol{\nu}) \propto \exp\left\{\sum_{j=1}^{J} -\frac{1}{2\tau^2}(\nu_j^2 - 2\nu_j\omega) + \sum_{n=1}^{N}\sum_{k=1}^{K} r_{nk}\left[-\frac{a_j}{2b_j}\left(-2x_{nj}\nu_j + 2y_{jk}m_{jk}\nu_j + \nu_j^2\right)\right]\right\}.$$

Next, we use the fact that $N_k = \sum_{n=1}^{N} r_{nk}$ and $\widehat{x}_{jk} = \sum_{i=1}^{N} r_{nk}x_{nj}$ to simplify the expression further where

$$q^*(\boldsymbol{\nu}) \propto \exp\left\{\sum_{j=1}^{J} -\frac{1}{2}\left(\frac{1}{\tau^2} + \frac{a_j}{b_j}\sum_{k=1}^{K} N_k\right)\nu_j^2 + \left(\frac{\omega}{\tau^2} + \frac{a_j}{b_j}\sum_{k=1}^{K}\left[\widehat{x}_{jk} - N_k y_{jk} m_{jk}\right]\right)\nu_j\right\}.$$

This results in a Gaussian quadratic where we again complete the square and simplify

$$\begin{aligned}
q^*(\boldsymbol{\nu}) &\propto \exp\left\{\sum_{j=1}^{J} -\frac{1}{2}\left(\frac{1}{\tau^2} + \frac{a_j}{b_j}\sum_{k=1}^{K} N_k\right)\left(\nu_j - \frac{(\omega/\tau^2) + \sum_{k=1}^{K}(a_j/b_j)\left[\widehat{x}_{jk} - N_k y_{jk} m_{jk}\right]}{1/\tau^2 + (a_j/b_j)\sum_{k=1}^{K} N_k}\right)^2\right\} \\
&\propto \exp\left\{\sum_{j=1}^{J} \ln \mathcal{N}\left(\frac{\omega/\tau^2 + (a_j/b_j)\sum_{k=1}^{K}\left[\widehat{x}_{jk} - N_k y_{jk} m_{jk}\right]}{1/\tau^2 + (a_j/b_j)\sum_{k=1}^{K} N_k}, \left[\frac{1}{\tau^2} + \frac{a_j}{b_j}\sum_{k=1}^{K} N_k\right]^{-1}\right)\right\} \\
&\propto \prod_{j=1}^{J} \mathcal{N}\left(\frac{\omega/\tau^2 + (a_j/b_j)\sum_{k=1}^{K}\left[\widehat{x}_{jk} - N_k y_{jk} m_{jk}\right]}{1/\tau^2 + (a_j/b_j)\sum_{k=1}^{K} N_k}, \left[\frac{1}{\tau^2} + \frac{a_j}{b_j}\sum_{k=1}^{K} N_k\right]^{-1}\right) \\
&\propto \prod_{j=1}^{J} \mathcal{N}\left(f_j, t_j^2\right)
\end{aligned}$$

This expression implies that the variational updates for $f_j$ and $t_j^2$ are the following

$$f_j = \frac{\omega/\tau^2 + (a_j/b_j)\sum_{k=1}^{K}\left[\widehat{x}_{jk} - N_k y_{jk} m_{jk}\right]}{1/\tau^2 + (a_j/b_j)\sum_{k=1}^{K} N_k}, \quad t_j^2 = \left[\frac{1}{\tau^2} + \frac{a_j}{b_j}\sum_{k=1}^{K} N_k\right]^{-1} \tag{30}$$

Similar to Eq. (29), in Eq. (30) we see that the model estimates the $f_j$ as the "residual" expression not captured by the phenotypic shift term, $m_{jk}$. An important distinction here is that the "residual" expression is across all $K$ clusters in Eq. (30). Therefore, one can alternatively view $f_j$ as the total expression not captured across all clusters. The other terms in the numerator $\omega$ and $\tau^2$ are the prior mean and variance of the baseline global mean expression. These also influence the estimates of $f_j$. If one has strong belief about the baseline global mean expression (such as the sample mean or sample median), these assumptions can be encoded into the model by setting $\omega$ to a specific values and fixing $\tau^2$

to be very small (e.g., $\tau^2 = 1 \times 10^{-12}$). This will encourage $f_j$ to be tightly centered around the prior $\omega$. In this work, we choose to let the data almost completely inform the baseline global mean expression. Therefore, we set $\tau^2 = 1 \times 10^{12}$, which reduces the impact of $\omega$ on the estimate.

##### 2.2.3 Variational Update to $\boldsymbol{a}_j$ and $\boldsymbol{b}_j$

In the specification of NCLUSION, $\sigma_j^2$ is the global variance of the $j$-th gene that is common across all clusters. Using Eq. (16), we find the variational updates for $a_j$ and $b_j$ by setting $\boldsymbol{\theta}_\ell = \boldsymbol{\sigma}^2$. This yields the following function for the update equation

$$q^*(\boldsymbol{\sigma}^2) \propto \exp\Bigg\{\mathbb{E}_{q(\boldsymbol{\Theta}\backslash\boldsymbol{\sigma}^2)}\left[\sum_{j=1}^{J} -(\xi_1 - 1)\ln\sigma_j^2 - \frac{\xi_2}{\sigma_j^2} + \sum_{k=1}^{K}\left(-\frac{1}{2}\rho_{jk}\ln\sigma_j^2 - \frac{1}{2\sigma_j^2\lambda_{jk}}\rho_{jk}\mu_{jk}^2\right)\right.$$
$$\left.\left. + \sum_{n=1}^{N}\sum_{k=1}^{K}[\psi_n = k]\left(-\frac{1}{2}\ln\sigma_j^2 - \frac{1}{2\sigma_j^2}\left[(x_{nj} - \nu_j - \rho_{jk}\mu_{jk})^2\right]\right)\right]\right\}.$$

When taking the expectation of each term (with respect to the other parameters in the model), we get the following closed-form expressions: $\mathbb{E}_q[\rho_{jk}] = y_{jk}$; $\mathbb{E}_q[1/\lambda_{jk}] = u_{jk}/v_{jk}$; $\mathbb{E}_q\left[\mu_{jk}^2\right] = m_{jk}^2 + s_{jk}^2$; and $\mathbb{E}_q[\psi_n = k] = r_{nk}$. The form of the final expectation term is $\mathbb{E}_q\left[(x_{nj} - \nu_j - \rho_{jk}\mu_{jk})^2\right] = x_{nj}^2 - 2x_{nj}(y_{jk}m_{jk} + f_j) + 2y_{jk}m_{jk}f_j + y_{jk}(m_{jk}^2 + s_{jk}^2) + (f_j^2 + t_j^2)$. Substituting these values into the expression above yields the following

$$q^*(\boldsymbol{\sigma}^2) \propto \exp\Bigg\{\sum_{j=1}^{J} -(\xi_1 - 1)\ln\sigma_j^2 - \frac{\xi_2}{\sigma_j^2} + \sum_{k=1}^{K}\left[-\frac{1}{2}y_{jk}\ln\sigma_j^2 - \frac{u_{jk}}{2\sigma_j^2 v_{jk}}y_{jk}\left(m_{jk}^2 + s_{jk}^2\right)\right]$$
$$+ \sum_{n=1}^{N}\sum_{k=1}^{K} r_{nk}\Bigg( - \frac{1}{2}\ln\sigma_j^2 - \frac{1}{2\sigma_j^2}\left[x_{nj}^2 - 2x_{nj}(y_{jk}m_{jk} + f_j) + 2y_{jk}m_{jk}f_j\right.$$
$$\left. + y_{jk}(m_{jk}^2 + s_{jk}^2) + (f_j^2 + t_j^2)\right]\Bigg)\Bigg\}.$$

Next, we leverage the fact that $N_k = \sum_{n=1}^{N} r_{nk}$, $\widehat{x}_{jk}^2 = \sum_{i=1}^{N} r_{nk}x_{nj}^2$ , and $\widehat{x}_{jk} = \sum_{i=1}^{N} r_{nk}x_{nj}$ and combine like-terms to get the following the expression,

$$q^*(\boldsymbol{\sigma}^2) \propto \exp\Bigg\{\sum_{j=1}^{J} -\left(\xi_1 + \frac{1}{2}\sum_{k=1}^{K}(N_k + y_{jk}) - 1\right)\ln\sigma_j^2 - \Bigg[\xi_2 + \frac{1}{2}\sum_{k=1}^{K}\frac{u_{jk}}{v_{jk}}y_{jk}\left(m_{jk}^2 + s_{jk}^2\right)$$
$$+ \frac{1}{2}\sum_{k=1}^{K}\widehat{x}_{jk}^2 - 2\widehat{x}_{jk}(y_{jk}m_{jk} + f_j) + N_k(y_{jk}m_{jk} + f_j)^2 + N_k(y_{jk}s_{jk}^2 + t_j^2)\Bigg]\frac{1}{\sigma_j^2}\Bigg\}.$$

This takes the form of an inverse-gamma distribution where

$$q^*(\boldsymbol{\sigma}^2) \propto \prod_{j=1}^{J} \text{Inv-Gamma}\Bigg(\xi_1 + \frac{1}{2}\sum_{k=1}^{K}(N_k + y_{jk}), \xi_2 + \frac{1}{2}\sum_{k=1}^{K} \frac{u_{jk}}{v_{jk}} y_{jk}\left(m_{jk}^2 + s_{jk}^2\right) + \widehat{x}_{jk}^2 - 2\widehat{x}_{jk}(y_{jk}m_{jk} + f_j)$$

$$+ N_k(y_{jk}m_{jk} + f_j)^2 + N_k(y_{jk}s_{jk}^2 + t_j^2)\Bigg)$$

$$\propto \prod_{j=1}^{J} \text{Inv-Gamma}(a_j, b_j)$$

where the variational updates for $a_j$ and $b_j$ are the following

$$a_j = \xi_1 + \frac{1}{2}\sum_{k=1}^{K}(N_k + y_{jk}) \tag{31}$$

$$\begin{aligned} b_j = \xi_2 + \frac{1}{2}\sum_{k=1}^{K} \frac{u_{jk}}{v_{jk}} y_{jk}\left(m_{jk}^2 + s_{jk}^2\right) + \widehat{x}_{jk}^2 - 2\widehat{x}_{jk}(y_{jk}m_{jk} + f_j) \\ + N_k(y_{jk}m_{jk} + f_j)^2 + N_k(y_{jk}s_{jk}^2 + t_j^2). \end{aligned} \tag{32}$$

##### 2.2.4 Variational Update to $\boldsymbol{u}_{jk}$ and $\boldsymbol{v}_{jk}$

The term $\lambda_{jk}$ is the $k$-th cluster-specific variance scaling component of the slab distribution for the $j$-th gene. Recall from Eq. (15) that $u_{jk}$ and $v_{jk}$ are the shape and rate parameters of the variational distribution that approximates $\lambda_{jk}$. By letting $\boldsymbol{\theta}_\ell = \boldsymbol{\lambda}$, we get the follow function to optimize

$$q^*(\boldsymbol{\lambda}) \propto \exp\left\{\mathbb{E}_{q(\boldsymbol{\Theta}\setminus\boldsymbol{\lambda})}\left[\sum_{j=1}^{J}\sum_{k=1}^{K} -(\kappa_1 - 1)\ln\lambda_{jk} - \frac{\kappa_2}{\lambda_{jk}} - \frac{1}{2}\rho_{jk}\ln\lambda_{jk} - \frac{1}{2\lambda_{jk}\sigma_{jk}^2}\rho_{jk}\mu_{jk}^2\right]\right\}.$$

We replace the parameters $\rho_{jk}$, $\sigma_j^2$, and $\mu_{jk}$ with the closed-form expression expectations taken with respect to their variational distributions. Namely, we have $\mathbb{E}_q\left[\rho_{jk}\right] = y_{jk}$, $\mathbb{E}_q\left[1/\sigma_j^2\right] = a_j/b_j$, and $\mathbb{E}_q\left[\mu_{jk}^2\right] = m_{jk}^2 + s_{jk}^2$. Substituting these values into the function above and simplifying yields

$$q^*(\boldsymbol{\lambda}) \propto \exp\left\{\sum_{j=1}^{J}\sum_{k=1}^{K} -\left(\kappa_1 + \frac{1}{2}y_{jk} - 1\right)\ln\lambda_{jk} - \left(\kappa_2 + \frac{a_j}{2b_j}y_{jk}(m_{jk}^2 + s_{jk}^2)\right)\frac{1}{\lambda_{jk}}\right\}.$$

Similar to the last section, this also represents an inverse-gamma distribution of the form

$$q^*(\boldsymbol{\lambda}) \propto \prod_{k=1}^{K}\prod_{j=1}^{J} \text{Inv-Gamma}\left(\kappa_1 + \frac{1}{2}y_{jk}, \kappa_2 + \frac{a_j}{2b_j}y_{jk}(m_{jk}^2 + s_{jk}^2)\right)$$
$$\propto \prod_{k=1}^{K}\prod_{j=1}^{J} \text{Inv-Gamma}(u_{jk}, v_{jk})$$

where the variational updates for $u_{jk}$ and $v_{jk}$ are the following

$$u_{jk} = \kappa_1 + \frac{1}{2}y_{jk}, \quad v_{jk} = \kappa_2 + \frac{a_j}{2b_j}y_{jk}(m_{jk}^2 + s_{jk}^2). \tag{33}$$

##### 2.2.5 Variational Update to $\boldsymbol{h}_1$ and $\boldsymbol{h}_2$

In the NCLUSION model specification, the term $\eta$ denotes the prior probability that the expression of $j$-th gene deviates from the global baseline expression in any given cluster. Looking at Eq. (17), $h_1$ and $h_2$ are the shape parameters of the variational Beta distribution that approximates $\eta$. To find the variational updates for $h_1$ and $h_2$, we let $\boldsymbol{\theta}_\ell = \eta$. This yields the following expression for the optimization

$$q^*(\eta) \propto \exp\left\{\mathbb{E}_{q(\boldsymbol{\Theta}\setminus\eta)}\left[\sum_{k=1}^{K}\sum_{j=1}^{J}\rho_{jk}\ln\eta + (1-\rho_{jk})\ln(1-\eta) + (\varphi_1 - 1)\ln\eta + (\varphi_2 - 1)\ln(1-\eta)\right]\right\}.$$

When taking the expectation of the terms involving $\rho_{jk}$, we see that $\mathbb{E}_q\left[\rho_{jk}\right] = y_{jk}$ and $\mathbb{E}_q\left[(1-\rho_{jk})\right] = (1-g_{1k})g_{2k}/\left[g_{1k}g_{2k} + (1-g_{1k})g_{2k}\right] = (1-y_{jk})$. Substituting these quantities into the optimization function gives the following

$$q^*(\eta) \propto \exp\left\{\left(\varphi_1 + \sum_{k=1}^{K}\sum_{j=1}^{J}y_{jk} - 1\right)\ln\eta + \left(\varphi_2 + JK - \sum_{k=1}^{K}\sum_{j=1}^{J}y_{jk} - 1\right)\ln(1-\eta)\right\}.$$

Note that this is a Beta distribution of the form

$$q^*(\eta) \propto \eta^{\left(\varphi_1 + \sum_{k=1}^{K}\sum_{j=1}^{J}y_{jk} - 1\right)} \times \ln(1-\eta)^{\left(\varphi_2 + JK - \sum_{k=1}^{K}\sum_{j=1}^{J}y_{jk} - 1\right)}$$
$$\propto \text{Beta}\left(\varphi_1 + \sum_{k=1}^{K}\sum_{j=1}^{J}y_{jk}, \varphi_2 + JK - \sum_{k=1}^{K}\sum_{j=1}^{J}y_{jk}\right)$$
$$\propto \text{Beta}\left(h_1, h_2\right)$$

where the variational updates for $h_1$ and $h_2$ are the following

$$h_1 = \varphi_1 + \sum_{k=1}^{K}\sum_{j=1}^{J} y_{jk}, \quad h_2 = \varphi_2 + JK - \sum_{k=1}^{K}\sum_{j=1}^{J} y_{jk}. \tag{34}$$

##### 2.2.6 Variational Update to $\boldsymbol{d}_k$

The $\pi_k$ in the mixture model represents the marginal probability that any cell will belong to the $k$-th cluster. Recall from Eq. (19) that the parameter $d_k$ is the analog to these normal mixture weights. By setting $\boldsymbol{\theta}_\ell = \boldsymbol{\pi}$, we get the following function to optimize over

$$q^*(\boldsymbol{\pi}) \propto \exp\left\{\mathbb{E}_{q(\boldsymbol{\Theta}\setminus\boldsymbol{\pi})}\left[\sum_{k=1}^{K}\left(\alpha_0\chi_k\prod_{l=1}^{k-1}[(1-\chi_l)] - 1 + \sum_{n=1}^{N}[\psi_n = k]\right)\ln\pi_k\right]\right\}.$$

Next, we can take the expectation of the terms involving the parameters $\chi_k$ and $\psi_n$ with respect to their approximating distribtutions which yields: $\mathbb{E}_q[\chi_k] = g_{1k}g_{2k}/[g_{1k}g_{2k} + (1-g_{1k})g_{2k}] = g_{1k}$, $\mathbb{E}_q[(1-\chi_k)] = 1 - \{g_{1k}g_{2k}/[g_{1k}g_{2k} + (1-g_{1k})g_{2k}]\} = (1-g_{1k})$, and $\mathbb{E}_q[\psi_n = k] = r_{nk}$. Substituting these expectation back into the function above and using the fact that $N_k = \sum_{n=1}^{N} r_{nk}$ yields the expression

$$q^*(\boldsymbol{\pi}) \propto \exp\left\{\sum_{k=1}^{K}\left(\alpha_0 g_{1k}\prod_{l=1}^{k-1}(1-g_{1l}) + N_k - 1\right)\ln\pi_k\right\}$$

We observe that $q^*(\boldsymbol{\pi})$ is a Dirichlet distribution of the form

$$\begin{aligned}
q^*(\boldsymbol{\pi}) &\propto \prod_{k=1}^{K}\pi_k^{\left(\alpha_0\times g_{1k}\times\prod_{l=1}^{k-1}(1-g_{1l})+N_k-1\right)} \\
&\propto \prod_{k=1}^{K}\pi_k^{(d_k-1)} \\
&\propto \mathrm{Dir}(d_1,\ldots,d_K)
\end{aligned}$$

where $d_k \in \{d_1, ..., d_K\}$ is updated using the posterior cluster membership probabilities estimated in the E-step of the algorithm via

$$d_k = \alpha_0 \times g_{1k} \times \prod_{l=1}^{k-1}(1-g_{1l}) + N_k. \tag{35}$$

Note that this update takes a very similar form to what is used for Dirichlet distributed random variables in classic Bayesian methods where the concentration parameters are updated by adding the counts for all the new observations in each of the $K$-categories (see Gelman et al.[2] for more discussion).

##### 2.2.7 Variational Update to $g_{1k}$ and $g_{2k}$

The variational parameters $g_{1k}$ and $g_{2k}$ are used to approximate the distribution for $\chi_k$ which is the latent variable that ultimately influences the total number of clusters that end up being used in the model. Theoretically, we could still use a proper beta distribution for $\boldsymbol{\chi}_k$, as it is still in the exponential family, but this has been shown to lead to a non-conjugate relationship with the variational distribution for $\pi_k$[11]. This is mainly due to the intractable expectation $\mathbb{E}_{q(\chi)}[\ln\Gamma(\sum_k \alpha_0\beta_k) - \ln\prod_k \Gamma(\alpha_0\beta_k)]$ which is used to compute the lower bound in Eq. (21). The lack of conjugacy also prevents us from updating $g_{1k}$ and $g_{2k}$ using the previously outlined procedure. Instead, following an alternative strategy by Hughes et al.[11], we chose to update $g_{1k}$ and $g_{2k}$ using stochastic gradient descent. To derive the objective function we used, we first modified Eq. (21) by placing a lower bound on this expectation

$$\ln\left[\frac{\Gamma(\sum_k \alpha_0\beta_k)}{\prod_k \Gamma(\alpha_0\beta_k)}\right] \geq K\,\ln\alpha_0 + \sum_{k=1}^{K}\ln\chi_k + \sum_{k=1}^{K}(K+1-k)\,\ln(1-\chi_k).$$

We refer the readers to Hughes et al.[11] for the proof that shows the above to be a valid bound for all $\alpha_0 > 0$. Replacing this surrogate bound within likelihood allows us to compute the expectation

$$\begin{aligned}
\mathbb{E}_{q(\chi)}\left\{\ln\left[\frac{\Gamma(\sum_k \alpha_0\beta_k)}{\prod_k \Gamma(\alpha_0\beta_k)}\right]\right\} &\geq \mathbb{E}_{q(\chi)}\left[K\,\ln\alpha_0 + \sum_{k=1}^{K}\ln\chi_k + \sum_{k=1}^{K}(K+1-k)\,\ln(1-\chi_k)\right] \\
&\geq K\,\ln\alpha_0 + \sum_{k=1}^{K}\mathbb{E}_{q(\chi)}\left[\ln\chi_k\right] + \sum_{k=1}^{K}(K+1-k)\,\mathbb{E}_{q(\chi)}\left[\ln(1-\chi_k)\right] \\
&\geq K\,\ln\alpha_0 + \sum_{k=1}^{K}\left[\boldsymbol{\Psi}(g_{1k}g_{2k}) - \boldsymbol{\Psi}(g_{2k})\right] + \sum_{k=1}^{K}(K+1-k)\left[\boldsymbol{\Psi}((1-g_{1k})g_{2k}) - \boldsymbol{\Psi}(g_{2k})\right].
\end{aligned}$$

In the NCLUSION software, we estimate $g_{1k}$ and $g_{2k}$ via a stochastic gradient descent optimization procedure where we use gradients to facilitate faster convergence to a local optima given the current estimate of all other parameters. The objective function for this step substitutes the surrogate bound above in the overall lower bound in Eq. (21) to create

$$
\begin{aligned}
\mathcal{L}^*(g_{1k}, g_{2k}) = &\sum_{k=1}^{K} \ln\left[\frac{\Gamma(g_{1k}g_{2k})\Gamma((1-g_{1k})g_{2k})}{\Gamma(g_{1k}g_{2k} + (1-g_{1k})g_{2k})}\right] + \sum_{k=1}^{K}(2 - g_{1k}g_{2k})\left[\boldsymbol{\Psi}(g_{1k}g_{2k}) - \boldsymbol{\Psi}(g_{2k})\right] \\
&+ \sum_{k=1}^{K}\left[(K+1-k) + \gamma_0 - (1-g_{1k})g_{2k}\right]\left[\boldsymbol{\Psi}((1-g_{1k})g_{2k}) - \boldsymbol{\Psi}(g_{2k})\right] \\
&+ \sum_{k=1}^{K+1}\alpha_0\left[\frac{g_{1k}g_{2k}}{g_{1k}g_{2k} + (1-g_{1k})g_{2k}}\prod_{l=1}^{k-1}\frac{(1-g_{1l})g_{2l}}{g_{1l}g_{2l} + (1-g_{1l})g_{2l}}\right]\left[\boldsymbol{\Psi}(d_k) - \boldsymbol{\Psi}\left(\sum_k d_k\right)\right].
\end{aligned}
\tag{36}
$$

As a result, the constrained optimization problem to solve for $g_{1k}$ and $g_{2k}$ takes on the joint form

$$
\left(\widehat{g}_k, \widehat{h}_k\right) = \underset{g_{1k}, g_{2k}}{\arg\max}\, \mathcal{L}^*(g_{1k}, g_{2k})
$$

subject to $0 < g_{1k} < 1$ and $g_{2k} > 0$ for $k = 1, \ldots, K$ cluster components. In this work, we use the gradients supplied in Hughes et al. [11] as optimal values for these parameters where, for every $l$-th entry $g_{1l}$, we have the following

$$
\begin{aligned}
\frac{\partial \mathcal{L}^*(g_{1k}, g_{2k})}{\partial g_{1l}} = &-g_{2l}\left[(K+1-k) + \gamma_0 - (1-g_{1l})g_{2l}\right]\boldsymbol{\Psi}'((1-g_{1l})g_{2l}) \\
&+ g_{2l}(2 - g_{1l}g_{2l})\boldsymbol{\Psi}'(g_{1l}g_{2l}) + \alpha_0\sum_{k=1}^{K}\Delta_{kl}\left[\boldsymbol{\Psi}(d_k) - \boldsymbol{\Psi}\left(\sum_k d_k\right)\right].
\end{aligned}
\tag{37}
$$

Similarly, for every $l$-th entry $g_{2l}$, we have the following

$$
\begin{aligned}
\frac{\partial \mathcal{L}^*(g_{1k}, g_{2k})}{\partial g_{2l}} = &\left[(K+1-k) + \gamma_0 - (1-g_{1l})g_{2l}\right]\left[(1-g_{1l})\boldsymbol{\Psi}'((1-g_{1l})g_{2l}) - \boldsymbol{\Psi}'(g_{2l})\right] \\
&+ (2 - g_{1l}g_{2l})\left[g_{1l}\boldsymbol{\Psi}'(g_{1l}g_{2l}) - \boldsymbol{\Psi}'(g_{2l})\right].
\end{aligned}
\tag{38}
$$

In both of the equations above, $\boldsymbol{\Psi}'(\cdot)$ is the first derivative of the digamma function and $\boldsymbol{\Delta}$ is defined as a $K \times (K+1)$ matrix of partial derivatives of $\mathbb{E}_{q(\chi)}[\beta_k]$ with respect to $g_{1l}$ such that

$$
\Delta_{kl} = \frac{\partial \mathbb{E}_{q(\chi)}[\beta_k]}{\partial g_{1l}} = \begin{cases} \mathbb{E}_{q(\chi)}[\beta_k]/(1-g_{1l}) & \text{if} \quad l < k \\ \mathbb{E}_{q(\chi)}[\beta_k]/g_{1l} & \text{if} \quad l = k \\ 0 & \text{if} \quad l > k \end{cases}
\tag{39}
$$

where it is straightforward to show that

$$\mathbb{E}_{q(\chi)}[\beta_k] = \left[ \frac{g_{1k}g_{2k}}{g_{1k}g_{2k} + (1-g_{1k})g_{2k}} \prod_{l=1}^{k-1} \frac{(1-g_{1l})g_{2l}}{g_{1l}g_{2l} + (1-g_{1l})g_{2l}} \right] = g_{1k} \prod_{l=1}^{k-1} (1-g_{1l}). \tag{40}$$

### 3 Full Derivation of the Evidence Lower Bound

In this section, we provide the full derivation for the evidence lower bound in Eq. (21). Here, we begin by partitioning the likelihood into the following equation:

$$\begin{aligned}
\mathcal{L}(\boldsymbol{\Theta}) &= \mathbb{E}_{q(\boldsymbol{\Theta})}\left[\ln p(\boldsymbol{\Theta}, \mathbf{X})\right] - \mathbb{E}_{q(\boldsymbol{\Theta})}\left[\ln q(\boldsymbol{\Theta})\right] \\
&= \mathbb{E}_{q(\boldsymbol{\Theta})}\left[\ln p(\mathbf{X} \mid \boldsymbol{\Theta}) + \ln p(\boldsymbol{\Theta}) - \ln q(\boldsymbol{\Theta})\right] \\
&= \mathbb{E}_{q(\boldsymbol{\Theta})} \Bigg\{ \ln p(\mathbf{X} \mid \boldsymbol{\Theta}) + \ln \Bigg[ p(\boldsymbol{\mu} \mid \boldsymbol{\rho}, \boldsymbol{\sigma}^2, \boldsymbol{\lambda}) p(\boldsymbol{\nu} \mid \omega, \tau^2) p(\boldsymbol{\rho} \mid \eta) p(\boldsymbol{\sigma^2} \mid \xi_1, \xi_2) p(\boldsymbol{\lambda} \mid \kappa_1, \kappa_2) \\
&\qquad p(\eta \mid \varphi_1, \varphi_2) p(\boldsymbol{\psi} \mid \boldsymbol{\pi}) p(\boldsymbol{\pi} \mid , \boldsymbol{\chi}, \alpha_0) p(\boldsymbol{\chi} \mid \gamma_0) \Bigg] \\
&\qquad - \ln \Bigg[ q(\boldsymbol{\mu} \mid \boldsymbol{m_\mu}, \boldsymbol{s^2_\mu}) q(\boldsymbol{\nu} \mid \boldsymbol{m_\nu}, \boldsymbol{s^2_\nu}) q(\boldsymbol{\rho} \mid \boldsymbol{y}) q(\boldsymbol{\sigma}^2 \mid \boldsymbol{a}, \boldsymbol{b}) q(\boldsymbol{\lambda} \mid \boldsymbol{u}, \boldsymbol{v}) \\
&\qquad q(\eta \mid h_1, h_2) q(\boldsymbol{\psi} \mid \boldsymbol{r}) q(\boldsymbol{\pi} \mid \boldsymbol{d}) q(\boldsymbol{\chi} \mid \boldsymbol{g_1}, \boldsymbol{g_2}) \Bigg] \Bigg\} \\
&= \mathbb{E}_{q(\boldsymbol{\Theta})} \left\{ \ln p(\mathbf{X} \mid \boldsymbol{\Theta}) + \ln \left[ \frac{p(\boldsymbol{\mu} \mid \boldsymbol{\rho}, \boldsymbol{\sigma}^2, \boldsymbol{\lambda}) p(\boldsymbol{\nu} \mid \omega, \tau^2) p(\boldsymbol{\rho} \mid \eta) p(\boldsymbol{\sigma}^2 \mid \xi_1, \xi_2) p(\boldsymbol{\lambda} \mid \kappa_1, \kappa_2) p(\eta \mid \varphi_1, \varphi_2)}{q(\boldsymbol{\mu} \mid \boldsymbol{m_\mu}, \boldsymbol{s^2_\mu}) q(\boldsymbol{\nu} \mid \boldsymbol{m_\nu}, \boldsymbol{s^2_\nu}) q(\boldsymbol{\sigma}^2 \mid \boldsymbol{a}, \boldsymbol{b}) q(\boldsymbol{\lambda} \mid \boldsymbol{u}, \boldsymbol{v}) q(\eta \mid h_1, h_2)} \right] \right\} \\
&\quad - \mathbb{E}_{q(\boldsymbol{\Theta})}\left[\ln q(\boldsymbol{\rho} \mid \boldsymbol{y})\right] + \mathbb{E}_{q(\boldsymbol{\Theta})} \left\{ \ln \left[ \frac{p(\boldsymbol{\psi} \mid \boldsymbol{\pi}) p(\boldsymbol{\pi} \mid , \boldsymbol{\chi}, \alpha_0)}{q(\boldsymbol{\pi} \mid \boldsymbol{d})} \right] \right\} - \mathbb{E}_{q(\boldsymbol{\Theta})}[\ln q(\boldsymbol{\psi} \mid \boldsymbol{r})] + \mathbb{E}_{q(\boldsymbol{\Theta})} \left\{ \ln \left[ \frac{p(\boldsymbol{\chi} \mid \gamma_0)}{q(\boldsymbol{\chi} \mid \boldsymbol{g_1}, \boldsymbol{g_2})} \right] \right\} \\
&= \mathcal{L}^{\text{Data}} + \mathcal{H}(\boldsymbol{\rho}) + \mathcal{L}^{\text{HDP}} + \mathcal{H}(\boldsymbol{\psi}) + \mathcal{L}^{\boldsymbol{\chi}}.
\end{aligned} \tag{41}$$

Below we further expand each component of the above equation.

#### 3.1 Derivation of the Data Likelihood

To begin, we will derive the part of the lower bound that relies on the data. Consequently, it also contains the sparsity promoting prior specification that facilitates variable selection. We can expand this expression and, using the linearity of expectations, distribute the expected value into the expression. Using the hierarchical model in Eqs. (1)-(4) and the corresponding variational families in Eqs. (12)-(20), we can write the following

$$
\begin{aligned}
\mathcal{L}^{\text{Data}} &= \mathbb{E}_{q(\boldsymbol{\Theta})}\left\{\ln p(\mathbf{X}\,|\,\boldsymbol{\Theta}) + \ln\left[\frac{p(\boldsymbol{\mu}\,|\,\boldsymbol{\rho},\boldsymbol{\sigma}^2,\boldsymbol{\lambda})p(\boldsymbol{\nu}\,|\,\omega,\tau^2)p(\boldsymbol{\rho}\,|\,\eta)p(\boldsymbol{\sigma}^2\,|\,\xi_1,\xi_2)p(\boldsymbol{\lambda}\,|\,\kappa_1,\kappa_2)p(\eta\,|\,\varphi_1,\varphi_2)}{q(\boldsymbol{\mu}\,|\,\boldsymbol{m_\mu},\boldsymbol{s^2_\mu})q(\boldsymbol{\nu}\,|\,\boldsymbol{m_\nu},\boldsymbol{s^2_\nu})q(\boldsymbol{\sigma}^2\,|\,\boldsymbol{a},\boldsymbol{b})q(\boldsymbol{\lambda}\,|\,\boldsymbol{u},\boldsymbol{v})q(\eta\,|\,h_1,h_2)}\right]\right\}\\
&= \mathbb{E}_{q(\boldsymbol{\Theta})}[\ln p(\mathbf{X}\,|\,\boldsymbol{\Theta})] + \mathbb{E}_{q(\boldsymbol{\Theta})}[\ln p(\boldsymbol{\mu}\,|\,\boldsymbol{\rho},\boldsymbol{\sigma}^2,\boldsymbol{\lambda})] + \mathbb{E}_{q(\boldsymbol{\Theta})}[\ln p(\boldsymbol{\nu}\,|\,\omega,\tau^2)] + \mathbb{E}_{q(\boldsymbol{\Theta})}[\ln p(\boldsymbol{\rho}\,|\,\eta)]\\
&+ \mathbb{E}_{q(\boldsymbol{\Theta})}[\ln p(\boldsymbol{\sigma}^2\,|\,\xi_1,\xi_2)] + \mathbb{E}_{q(\boldsymbol{\Theta})}[\ln p(\boldsymbol{\lambda}\,|\,\kappa_1,\kappa_2)] + \mathbb{E}_{q(\boldsymbol{\Theta})}[\ln p(\eta\,|\,\varphi_1,\varphi_2)]\\
&- \mathbb{E}_{q(\boldsymbol{\Theta})}[\ln q(\boldsymbol{\mu}\,|\,\boldsymbol{m_\mu},\boldsymbol{s^2_\mu})] - \mathbb{E}_{q(\boldsymbol{\Theta})}[\ln q(\boldsymbol{\nu}\,|\,\boldsymbol{m_\nu},\boldsymbol{s^2_\nu})] - \mathbb{E}_{q(\boldsymbol{\Theta})}[\ln q(\boldsymbol{\sigma}^2\,|\,\boldsymbol{a},\boldsymbol{b})]\\
&- \mathbb{E}_{q(\boldsymbol{\Theta})}[\ln q(\boldsymbol{\lambda}\,|\,\boldsymbol{u},\boldsymbol{v})] - \mathbb{E}_{q(\boldsymbol{\Theta})}[\ln q(\eta\,|\,h_1,h_2)]\\
&= \mathbb{E}_{q(\boldsymbol{\Theta})}\left\{\sum_{n=1}^{N}\sum_{j=1}^{J}\sum_{k=1}^{K}[\psi_n = k]\left[-\frac{1}{2}\ln 2\pi - \frac{1}{2}\ln\sigma_j^2 - \frac{1}{2\sigma_j^2}(x_{nj} - \nu_j - \rho_{jk}\mu_{jk})^2\right]\right\}\\
&+ \mathbb{E}_{q(\boldsymbol{\Theta})}\left\{\sum_{j=1}^{J}\sum_{k=1}^{K}\rho_{jk}\left(-\frac{1}{2}\ln 2\pi - \frac{1}{2}\ln\lambda_{jk} - \frac{1}{2}\ln\sigma_j^2 - \frac{1}{2\lambda_{jk}\sigma_j^2}\mu_{jk}^2\right)\right\}\\
&+ \mathbb{E}_{q(\boldsymbol{\Theta})}\left\{\sum_{j=1}^{J} -\frac{1}{2}\ln 2\pi - \frac{1}{2}\ln\tau^2 - \frac{1}{2\tau^2}(\nu_j - \omega)^2\right\} + \mathbb{E}_{q(\boldsymbol{\Theta})}\left\{\sum_{j=1}^{J}\sum_{k=1}^{K}\rho_{jk}\ln\eta + (1-\rho_{jk})\,\ln(1-\eta)\right\}\\
&+ \mathbb{E}_{q(\boldsymbol{\Theta})}\left\{\sum_{j=1}^{J}\ln\frac{\xi_2^{\xi_1}}{\Gamma(\xi_1)} - (\xi_1 - 1)\ln\sigma_j^2 - \frac{\xi_2}{\sigma_j^2}\right\}\\
&+ \mathbb{E}_{q(\boldsymbol{\Theta})}\left\{\sum_{j=1}^{J}\sum_{k=1}^{K}\ln\frac{\kappa_2^{\kappa_1}}{\Gamma(\kappa_1)} - (\kappa_1 - 1)\ln\lambda_{jk} - \frac{\kappa_2}{\lambda_{jk}}\right\}\\
&+ \mathbb{E}_{q(\boldsymbol{\Theta})}\left\{-\ln\frac{\Gamma(\varphi_1)\Gamma(\varphi_2)}{\Gamma(\varphi_1+\varphi_2)} + (\varphi_1 - 1)\ln\eta + (\varphi_2 - 1)\ln(1-\eta)\right\} \qquad (42)\\
&+ \mathbb{E}_{q(\boldsymbol{\Theta})}\left\{\sum_{j=1}^{J}\sum_{k=1}^{K}\rho_{jk}\left[\frac{1}{2}\ln 2\pi + \frac{1}{2}\ln s_{jk}^2 + \frac{1}{2s_{jk}^2}(\mu_{jk} - m_{jk})^2\right]\right\}\\
&+ \mathbb{E}_{q(\boldsymbol{\Theta})}\left\{\sum_{j=1}^{J}\frac{1}{2}\ln 2\pi + \frac{1}{2}\ln t_j^2 + \frac{1}{2t_j^2}\left(\nu_j - f_j\right)^2\right\}\\
&+ \mathbb{E}_{q(\boldsymbol{\Theta})}\left\{\sum_{j=1}^{J} -\ln\frac{b_j^{a_j}}{\Gamma(a_j)} + (a_j - 1)\ln\sigma_j^2 + \frac{b_j}{\sigma_j^2}\right\}\\
&+ \mathbb{E}_{q(\boldsymbol{\Theta})}\left\{\sum_{j=1}^{J}\sum_{k=1}^{K} -\ln\frac{v_{jk}^{u_{jk}}}{\Gamma(u_{jk})} + (u_{jk} - 1)\ln\lambda_{jk} + \frac{v_{jk}}{\lambda_{jk}}\right\}\\
&+ \mathbb{E}_{q(\boldsymbol{\Theta})}\left\{\ln\frac{\Gamma(h_1)\Gamma(h_2)}{\Gamma(h_1+h_2)} - (h_1 - 1)\ln\eta - (h_2 - 1)\ln(1-\eta)\right\}.
\end{aligned}
$$

We can now distribute the expectation throughout the expression and group together similar terms to get the following

$$\begin{aligned}
\mathcal{L}^{\text{Data}} &= \sum_{n=1}^{N}\sum_{j=1}^{J}\sum_{k=1}^{K}\mathbb{E}_{q(\boldsymbol{\Theta})}[\psi_n = k]\left\{-\frac{1}{2}\ln 2\pi - \frac{1}{2}\mathbb{E}_{q(\boldsymbol{\Theta})}\left[\ln\sigma_j^2\right] - \mathbb{E}_{q(\boldsymbol{\Theta})}\left[\frac{1}{2\sigma_j^2}\right]\mathbb{E}_{q(\boldsymbol{\Theta})}\left[(x_{nj} - \nu_j - \rho_{jk}\mu_{jk})^2\right]\right\} \\
&+ \sum_{j=1}^{J}\sum_{k=1}^{K}\mathbb{E}_{q(\boldsymbol{\Theta})}\left[\rho_{jk}\right]\left(-\frac{1}{2}\ln 2\pi - \frac{1}{2}\mathbb{E}_{q(\boldsymbol{\Theta})}\left[\ln\lambda_{jk}\right] - \frac{1}{2}\mathbb{E}_{q(\boldsymbol{\Theta})}\left[\ln\sigma_j^2\right] - \mathbb{E}_{q(\boldsymbol{\Theta})}\left[\frac{1}{2\lambda_{jk}\sigma_j^2}\right]\mathbb{E}_{q(\boldsymbol{\Theta})}\left[\mu_{jk}^2\right]\right) \\
&+ \sum_{j=1}^{J}\sum_{k=1}^{K}\mathbb{E}_{q(\boldsymbol{\Theta})}\left[\rho_{jk}\right]\left[\frac{1}{2}\ln 2\pi + \frac{1}{2}\ln s_{jk}^2 + \frac{1}{2s_{jk}^2}\mathbb{E}_{q(\boldsymbol{\Theta})}\left[(\mu_{jk} - m_{jk})^2\right]\right] \\
&+ \sum_{j=1}^{J} -\frac{1}{2\tau^2}\mathbb{E}_{q(\boldsymbol{\Theta})}\left[(\nu_j - \omega)^2\right] + \frac{1}{2}\ln t_j^2 + \frac{1}{2t_j^2}\mathbb{E}_{q(\boldsymbol{\Theta})}\left[(\nu_j - f_j)^2\right] \\
&+ \sum_{j=1}^{J}\sum_{k=1}^{K}\mathbb{E}_{q(\boldsymbol{\Theta})}\left[\rho_{jk}\right]\mathbb{E}_{q(\boldsymbol{\Theta})}\left[\ln\eta\right] + \mathbb{E}_{q(\boldsymbol{\Theta})}\left[(1 - \rho_{jk})\right]\mathbb{E}_{q(\boldsymbol{\Theta})}\left[\ln(1 - \eta)\right] \\
&- \sum_{j=1}^{J}(\xi_1 - 1)\mathbb{E}_{q(\boldsymbol{\Theta})}\left[\ln\sigma_j^2\right] - \xi_2\mathbb{E}_{q(\boldsymbol{\Theta})}\left[\frac{1}{\sigma_j^2}\right] - \ln\frac{b_j^{a_j}}{\Gamma(a_j)} + (a_j - 1)\mathbb{E}_{q(\boldsymbol{\Theta})}\left[\ln\sigma_j^2\right] + b_j\mathbb{E}_{q(\boldsymbol{\Theta})}\left[\frac{1}{\sigma_j^2}\right] \\
&+ \sum_{j=1}^{J}\sum_{k=1}^{K} -(\kappa_1 - 1)\mathbb{E}_{q(\boldsymbol{\Theta})}\left[\ln\lambda_{jk}\right] - \kappa_2\mathbb{E}_{q(\boldsymbol{\Theta})}\left[\frac{1}{\lambda_{jk}}\right] - \ln\frac{v_{jk}^{u_{jk}}}{\Gamma(u_{jk})} + (u_{jk} - 1)\mathbb{E}_{q(\boldsymbol{\Theta})}\left[\ln\lambda_{jk}\right] + v_{jk}\mathbb{E}_{q(\boldsymbol{\Theta})}\left[\frac{1}{\lambda_{jk}}\right] \\
&+ (\varphi_1 - 1)\mathbb{E}_{q(\boldsymbol{\Theta})}\left[\ln\eta\right] + (\varphi_2 - 1)\mathbb{E}_{q(\boldsymbol{\Theta})}\left[\ln(1 - \eta)\right] - (h_1 - 1)\mathbb{E}_{q(\boldsymbol{\Theta})}\left[\ln\eta\right] - (h_2 - 1)\mathbb{E}_{q(\boldsymbol{\Theta})}\left[\ln(1 - \eta)\right] \\
&+ J\ln\frac{\xi_2^{\xi_1}}{\Gamma(\xi_1)} - \ln\frac{\Gamma(\varphi_1)\Gamma(\varphi_2)}{\Gamma(\varphi_1 + \varphi_2)} - \frac{J}{2}\ln 2\pi + \frac{J}{2}\ln 2\pi - \frac{J}{2}\ln\tau^2 + KJ\ln\frac{\kappa_2^{\kappa_1}}{\Gamma(\kappa_1)} + \ln\frac{\Gamma(h_1)\Gamma(h_2)}{\Gamma(h_1 + h_2)}.
\end{aligned} \tag{43}$$

Here, we can expand the squared expectations. Beginning with the first one, we have

$$\begin{aligned}
\mathbb{E}_{q(\boldsymbol{\Theta})}\left[(x_{nj} - \nu_j - \rho_{jk}\mu_{jk})^2\right] &= x_{nj}^2 - 2x_{nj}\mathbb{E}_{q(\boldsymbol{\Theta})}[\rho_{jk}]\mathbb{E}_{q(\boldsymbol{\Theta})}[\mu_{jk}] - 2x_{nj}\mathbb{E}_{q(\boldsymbol{\Theta})}[\nu_j] \\
&+ 2\mathbb{E}_{q(\boldsymbol{\Theta})}[\rho_{jk}]\mathbb{E}_{q(\boldsymbol{\Theta})}[\mu_{jk}]\mathbb{E}_{q(\boldsymbol{\Theta})}[\nu_j] + \mathbb{E}_{q(\boldsymbol{\Theta})}[\rho_{jk}]\mathbb{E}_{q(\boldsymbol{\Theta})}[\mu_{jk}^2] + \mathbb{E}_{q(\boldsymbol{\Theta})}[\nu_j^2] \\
&= x_{nj}^2 - 2x_{nj}(y_{jk}m_{jk} + f_j) + 2y_{jk}m_{jk}f_j + y_{jk}(m_{jk}^2 + s_{jk}^2) + (f_j^2 + t_j^2) \\
&= x_{nj}^2 - 2x_{nj}\mathbb{E}_{q(\boldsymbol{\Theta})}[(\rho_{jk}\mu_{jk} + \nu_j)] + \mathbb{E}_{q(\boldsymbol{\Theta})}[(\rho_{jk}\mu_{jk} + \nu_j)^2]
\end{aligned}$$

where $\mathbb{E}_{q(\boldsymbol{\Theta})}[\rho_{jk}] = y_{jk}$; $\mathbb{E}_{q(\boldsymbol{\Theta})}[\mu_{jk}] = m_{jk}$; $\mathbb{E}_{q(\boldsymbol{\Theta})}[\nu_j] = f_j$; $\mathbb{E}_{q(\boldsymbol{\Theta})}[\mu_{jk}^2] = (m_{jk}^2 + s_{jk}^2)$; and $\mathbb{E}_{q(\boldsymbol{\Theta})}[\nu_j^2] = (f_j^2 + t_j^2)$. Next, we can expand the squared expectations containing $\nu_j$ and its prior mean where$\omega$

$$\mathbb{E}_{q(\boldsymbol{\Theta})}\left[(\nu_j - \omega)^2\right] = \mathbb{E}_{q(\boldsymbol{\Theta})}[\nu_j^2] - 2\mathbb{E}_{q(\boldsymbol{\Theta})}[\nu_j]\omega + \omega^2$$

$$= (f_j^2 + t_j^2) - 2f_j\omega + \omega^2.$$

We can similarly simplify the following

$$\begin{aligned}\mathbb{E}_{q(\boldsymbol{\Theta})}\left[(\mu_{jk} - m_{jk})^2\right] &= \mathbb{E}_{q(\boldsymbol{\Theta})}[\mu_{jk}^2] - 2m_{jk}\mathbb{E}_{q(\boldsymbol{\Theta})}[\mu_{jk}] + m_{jk}^2\\ &= (m_{jk}^2 + s_{jk}^2) - 2m_{jk}^2 + m_{jk}^2\\ &= s_{jk}^2 - m_{jk}^2 + m_{jk}^2\\ &= s_{jk}^2.\end{aligned}$$

We simplify the final squared expectations with the following

$$\begin{aligned}\mathbb{E}_{q(\boldsymbol{\Theta})}\left[(\nu_j - f_j)^2\right] &= \mathbb{E}_{q(\boldsymbol{\Theta})}[\nu_j^2] - 2f_j\mathbb{E}_{q(\boldsymbol{\Theta})}[\nu_j] + f_j^2\\ &= (f_j^2 + t_j^2) - 2f_j^2 + f_j^2\\ &= t_j^2 - f_j^2 + f_j^2\\ &= t_j^2.\end{aligned}$$

Next, we substitute the squared expectations terms back into Eq. (43). Furthermore, we also replace the expectations of the following using the corresponding variational distribution assumptions: $\mathbb{E}_q\left[\rho_{jk}\right] = y_{jk}$; $\mathbb{E}_q\left[\ln\sigma_j^2\right] = \ln b_j - \boldsymbol{\Psi}(a_j)$; $\mathbb{E}_q\left[\ln\lambda_{jk}\right] = \ln v_{jk} - \boldsymbol{\Psi}(u_{jk})$; $\mathbb{E}_q\left[\ln\eta\right] = \boldsymbol{\Psi}\left(h_1\right) - \boldsymbol{\Psi}\left(h_1 + h_2\right)$; $\mathbb{E}_q\left[\ln(1-\eta)\right] = \boldsymbol{\Psi}\left(h_2\right) - \boldsymbol{\Psi}\left(h_1 + h_2\right)$; $\mathbb{E}_q\left[1/\lambda_{jk}\right] = u_{jk}/v_{jk}$; $\mathbb{E}_q\left[1/\sigma_j^2\right] = a_j/b_j$; $\mathbb{E}_{q(\boldsymbol{\Theta})}[\psi_n = k] = r_{nk}$; and $E_{q(\boldsymbol{\Theta})}[\rho_{jk}] = y_{jk}$. This yields the following simplified expression

$$\begin{aligned}\mathcal{L}^{\text{Data}} = \sum_{n=1}^{N}\sum_{j=1}^{J}\sum_{k=1}^{K} r_{nk}\Bigg\{ &-\frac{1}{2}\ln 2\pi - \frac{1}{2}\left[\ln b_j - \boldsymbol{\Psi}(a_j)\right]\\ &- \frac{a_j}{2b_j}\left[x_{nj}^2 - 2x_{nj}\mathbb{E}_{q(\boldsymbol{\Theta})}[(\rho_{jk}\mu_{jk} + \nu_j)] + \mathbb{E}_{q(\boldsymbol{\Theta})}[(\rho_{jk}\mu_{jk} + \nu_j)^2]\right]\Bigg\}\\ &+ \sum_{j=1}^{J}\sum_{k=1}^{K} y_{jk}\left(\frac{1}{2}\ln s_{jk}^2 + \frac{1}{2} - \frac{1}{2}\left[\ln v_{jk} - \boldsymbol{\Psi}(u_{jk})\right] - \frac{1}{2}\left[\ln b_j - \boldsymbol{\Psi}(a_j)\right] - \frac{1}{2}\left[\frac{u_{jk}}{v_{jk}}\right]\left[\frac{a_j}{b_j}\right]\mathbb{E}_{q(\boldsymbol{\Theta})}\left[\mu_{jk}^2\right]\right)\\ &+ \frac{J}{2} + \sum_{j=1}^{J} -\frac{1}{2\tau^2}\left[\mathbb{E}_{q(\boldsymbol{\Theta})}[\nu_j^2] - 2\mathbb{E}_{q(\boldsymbol{\Theta})}[\nu_j]\omega + \omega^2\right] + \frac{1}{2}\ln t_j^2\end{aligned}$$

$$+\left(\varphi_1 - h_1 + \sum_{j=1}^{J}\sum_{k=1}^{K} y_{jk}\right)\left[\boldsymbol{\Psi}\left(h_1\right) - \boldsymbol{\Psi}\left(h_1 + h_2\right)\right]$$
$$+\left(\varphi_2 - h_2 + jk - \sum_{j=1}^{J}\sum_{k=1}^{K} y_{jk}\right)\left[\boldsymbol{\Psi}\left(h_2\right) - \boldsymbol{\Psi}\left(h_1 + h_2\right)\right]$$
$$-\sum_{j=1}^{J}(\xi_1 - a_j)\left[\ln b_j - \boldsymbol{\Psi}(a_j)\right] - (\xi_2 - b_j)\left[\frac{a_j}{b_j}\right] - \ln\frac{b_j^{a_j}}{\Gamma(a_j)}$$
$$+\sum_{j=1}^{J}\sum_{k=1}^{K} -(\kappa_1 - u_{jk})\left[\ln v_{jk} - \boldsymbol{\Psi}(u_{jk})\right] - (\kappa_2 - v_{jk})\left[\frac{u_{jk}}{v_{jk}}\right] - \ln\frac{v_{jk}^{u_{jk}}}{\Gamma(u_{jk})}$$
$$+ J\ln\frac{\xi_2^{\xi_1}}{\Gamma(\xi_1)} - \ln\frac{\Gamma(\varphi_1)\Gamma(\varphi_2)}{\Gamma(\varphi_1+\varphi_2)} - \frac{J}{2}\ln\tau^2 + KJ\ln\frac{\kappa_2^{\kappa_1}}{\Gamma(\kappa_1)} + \ln\frac{\Gamma(h_1)\Gamma(h_2)}{\Gamma(h_1+h_2)}.$$

We can simplify this expression even further. We do this by first recognizing that the sum over the sample index $N$ only affects the values of $r_{nk}$ and $x_{nj}$. By defining the following summary statistics, $N_k = \sum_{i=1}^{N} r_{nk}$, $\widehat{x}_{ij} = \sum_{i=1}^{N} r_{nk}x_{nj}$, and $\widehat{x}_{ij}^2 = \sum_{i=1}^{N} r_{nk}x_{nj}^2$ the inner expectation gives us the reduced expression

$$\mathcal{L}^{\text{Data}} = \sum_{j=1}^{J}\sum_{k=1}^{K}\left\{-\frac{N_k}{2}\ln 2\pi - \frac{N_k}{2}\left[\ln b_j - \boldsymbol{\Psi}(a_j)\right] - \frac{1}{2}\left[\frac{a_j}{b_j}\right]\left[\widehat{x}_{jk}^2 - 2\widehat{x}_{jk}\mathbb{E}_{q(\boldsymbol{\Theta})}[(\rho_{jk}\mu_{jk}+\nu_j)] + N_k\mathbb{E}_{q(\boldsymbol{\Theta})}[(\rho_{jk}\mu_{jk}+\nu_j)^2]\right]\right\}$$
$$+\sum_{j=1}^{J}\sum_{k=1}^{K} y_{jk}\left(\frac{1}{2}\ln s_{jk}^2 + \frac{1}{2} - \frac{1}{2}\left[\ln v_{jk} - \boldsymbol{\Psi}(u_{jk})\right] - \frac{1}{2}\left[\ln b_j - \boldsymbol{\Psi}(a_j)\right] - \frac{1}{2}\left[\frac{u_{jk}}{v_{jk}}\right]\left[\frac{a_j}{b_j}\right]\mathbb{E}_{q(\boldsymbol{\Theta})}\left[\mu_{jk}^2\right]\right)$$
$$+\sum_{j=1}^{J} -\frac{1}{2\tau^2}\left[\mathbb{E}_{q(\boldsymbol{\Theta})}[\nu_j^2] - 2\mathbb{E}_{q(\boldsymbol{\Theta})}[\nu_j]\omega\right] + \frac{1}{2}\ln t_j^2$$
$$+\left(\varphi_1 - h_1 + \sum_{j=1}^{J}\sum_{k=1}^{K} y_{jk}\right)\left[\boldsymbol{\Psi}\left(h_1\right) - \boldsymbol{\Psi}\left(h_1+h_2\right)\right] + \left(\varphi_2 - h_2 + jk - \sum_{j=1}^{J}\sum_{k=1}^{K} y_{jk}\right)\left[\boldsymbol{\Psi}\left(h_2\right) - \boldsymbol{\Psi}\left(h_1+h_2\right)\right]$$
$$-\sum_{j=1}^{J}(\xi_1 - a_j)\left[\ln b_j - \boldsymbol{\Psi}(a_j)\right] - (\xi_2 - b_j)\left[\frac{a_j}{b_j}\right] - \ln\frac{b_j^{a_j}}{\Gamma(a_j)}$$
$$+\sum_{j=1}^{J}\sum_{k=1}^{K} -(\kappa_1 - u_{jk})\left[\ln v_{jk} - \boldsymbol{\Psi}(u_{jk})\right] - (\kappa_2 - v_{jk})\left[\frac{u_{jk}}{v_{jk}}\right] - \ln\frac{v_{jk}^{u_{jk}}}{\Gamma(u_{jk})}$$
$$+ J\ln\frac{\xi_2^{\xi_1}}{\Gamma(\xi_1)} - \ln\frac{\Gamma(\varphi_1)\Gamma(\varphi_2)}{\Gamma(\varphi_1+\varphi_2)} - \frac{J}{2}\ln\tau^2 + KJ\ln\frac{\kappa_2^{\kappa_1}}{\Gamma(\kappa_1)} + \ln\frac{\Gamma(h_1)\Gamma(h_2)}{\Gamma(h_1+h_2)} + \frac{J}{2} - \frac{J\omega^2}{2\tau^2}.$$

As a final step, we collect the expected values of the log variances terms of the prior and rewrite the terms to to save space so that

$$
\begin{aligned}
\mathcal{L}^{\text{Data}} &= \sum_{j=1}^{J}\sum_{k=1}^{K}\left\{-\frac{N_k}{2}\ln 2\pi - \frac{N_k}{2}\left[\ln b_j - \boldsymbol{\Psi}(a_j)\right] - \frac{1}{2}\left[\frac{a_j}{b_j}\right]\left[\widehat{x}_{jk}^2 - 2\widehat{x}_{jk}\mathbb{E}_{q(\boldsymbol{\Theta})}[(\rho_{jk}\mu_{jk}+\nu_j)] + N_k\mathbb{E}_{q(\boldsymbol{\Theta})}[(\rho_{jk}\mu_{jk}+\nu_j)^2]\right]\right\} \\
&+ \sum_{j=1}^{J}\sum_{k=1}^{K} - \left(\kappa_1 + \frac{1}{2}y_{jk} - u_{jk}\right)\left[\ln v_{jk} - \boldsymbol{\Psi}(u_{jk})\right] - \kappa_2\left[\frac{u_{jk}}{v_{jk}}\right] - \frac{1}{2}y_{jk}\left[\frac{u_{jk}}{v_{jk}}\right]\left[\frac{a_j}{b_j}\right]\mathbb{E}_{q(\boldsymbol{\Theta})}\left[\mu_{jk}^2\right] + \frac{1}{2}y_{jk} + \frac{1}{2}y_{jk}\ln s_{jk}^2 \\
&+ \sum_{j=1}^{J} - \ln\frac{b_j^{a_j}}{\Gamma(a_j)} + a_j + \sum_{j=1}^{J}\sum_{k=1}^{K} - \ln\frac{v_{jk}^{u_{jk}}}{\Gamma(u_{jk})} + u_{jk} \\
&+ \sum_{j=1}^{J} - \left(\xi_1 + \frac{1}{2}\sum_{k=1}^{K} y_{jk} - a_j\right)\left[\ln b_j - \boldsymbol{\Psi}(a_j)\right] - \xi_2\left[\frac{a_j}{b_j}\right] - \frac{1}{2\tau^2}\left[\mathbb{E}_{q(\boldsymbol{\Theta})}[\nu_j^2] - 2\mathbb{E}_{q(\boldsymbol{\Theta})}[\nu_j]\omega\right] + \frac{1}{2}\ln\tau^2 \\
&+ \left(\varphi_1 - h_1 + \sum_{j=1}^{J}\sum_{k=1}^{K} y_{jk}\right)\left[\boldsymbol{\Psi}(h_1) - \boldsymbol{\Psi}(h_1+h_2)\right] + \left(\varphi_2 - h_2 + jk - \sum_{j=1}^{J}\sum_{k=1}^{K} y_{jk}\right)\left[\boldsymbol{\Psi}(h_2) - \boldsymbol{\Psi}(h_1+h_2)\right] \\
&- \ln\frac{\Gamma(\varphi_1)\Gamma(\varphi_2)}{\Gamma(\varphi_1+\varphi_2)} + \ln\frac{\Gamma(h_1)\Gamma(h_2)}{\Gamma(h_1+h_2)} + KJ\ln\frac{\kappa_2^{\kappa_1}}{\Gamma(\kappa_1)} + J\ln\frac{\xi_2^{\xi_1}}{\Gamma(\xi_1)} - \frac{J}{2}\ln\tau^2 - \frac{J\omega^2}{2\tau^2} + \frac{J}{2}.
\end{aligned}
$$

which results in the first portion of the lower bound in Eq. (21).

#### 3.2 Derivation of the Gene Importance Entropy for the Likelihood

Next, we will expand the component of the lower bound that represents the entropy of the cluster-specific, gene importance weights. Recall that in Eqs. (14) and (41) we defined

$$
\begin{aligned}
\mathcal{H}(\boldsymbol{\rho}) &= -\mathbb{E}_{q(\boldsymbol{\Theta})}[\ln q(\boldsymbol{\rho}\,|\,\boldsymbol{y})] \\
&= -\mathbb{E}_{q(\boldsymbol{\Theta})}\left[\sum_{j=1}^{J}\sum_{k=1}^{K}\rho_{jk}\ln y_{jk} + (1-\rho_{jk})\,\ln(1-y_{jk})\right].
\end{aligned}
\tag{44}
$$

As we did in the previous subsection, we can distribute the expectation throughout the expression and use the fact that $E_{q(\boldsymbol{\Theta})}[\rho_{jk}] = y_{jk}$ to get the following simplified form

$$
\mathcal{H}(\boldsymbol{\rho}) = \sum_{j=1}^{J}\sum_{k=1}^{K} -y_{jk}\ln y_{jk} - (1-y_{jk})\,\ln(1-y_{jk}) \tag{45}
$$

which results in the second portion of the lower bound in Eq. (21).

#### 3.3 Likelihood Corresponding to the Lower Level of the Dirichlet Process

Next, we will continue by expanding the partition of the lower bound that contains the lower level of the Dirichlet process and the categorical distribution prior that estimates cell-to-cluster assignment. Once again, using the hierarchical model in Eqs. (1)-(4) and the corresponding variational families in Eqs. (12)-(20), we can write the following from Eq. (41)

$$\begin{aligned}
\mathcal{L}^{\text{HDP}} &= \mathbb{E}_{q(\boldsymbol{\Theta})}\left\{\ln\left[\frac{p(\boldsymbol{\psi}\,|\,\boldsymbol{\pi})p(\boldsymbol{\pi}\,|\,,\boldsymbol{\chi},\alpha_0)}{q(\boldsymbol{\pi}\,|\,\boldsymbol{d})}\right]\right\} \\
&= \mathbb{E}_{q(\boldsymbol{\Theta})}[\ln p(\boldsymbol{\psi}\,|\,\boldsymbol{\pi})] + \mathbb{E}_{q(\boldsymbol{\Theta})}[\ln p(\boldsymbol{\pi}\,|\,,\boldsymbol{\chi},\alpha_0)] - \mathbb{E}_{q(\boldsymbol{\Theta})}[\ln q(\boldsymbol{\pi}\,|\,\boldsymbol{d})] \\
&= \mathbb{E}_{q(\boldsymbol{\Theta})}\left[\sum_{i=1}^{N}\sum_{k=1}^{K+1}[\psi_n = k]\ln\pi_k\right] + \mathbb{E}_{q(\boldsymbol{\Theta})}\left\{\ln\left[\frac{\Gamma(\sum_k \alpha_0\beta_k)}{\prod_k\Gamma(\alpha_0\beta_k)}\right]\sum_{k=1}^{K+1}(\alpha_0\beta_k - 1)\ln\pi_k\right\} \\
&\quad - \mathbb{E}_{q(\boldsymbol{\Theta})}\left\{\ln\left[\frac{\Gamma(\sum_k d_k)}{\prod_k\Gamma(d_k)}\right]\sum_{k=1}^{K+1}(d_k - 1)\ln\pi_k\right\}
\end{aligned} \tag{46}$$

where $\Gamma(\cdot)$ is used to denote the gamma function. We again distribute the expectations such that

$$\begin{aligned}
\mathcal{L}^{\text{HDP}} &= \sum_{i=1}^{N}\sum_{k=1}^{K+1}\mathbb{E}_{q(\boldsymbol{\Theta})}[\psi_n = k]\,\mathbb{E}_{q(\boldsymbol{\Theta})}[\ln\pi_k] + \mathbb{E}_{q(\boldsymbol{\Theta})}\left\{\ln\left[\frac{\Gamma(\sum_k \alpha_0\beta_k)}{\prod_k\Gamma(\alpha_0\beta_k)}\right]\right\} - \ln\left[\frac{\Gamma(\sum_k d_k)}{\prod_k\Gamma(d_k)}\right] \\
&\quad + \sum_{k=1}^{K+1}(\alpha_0\mathbb{E}_{q(\boldsymbol{\Theta})}[\beta_k] - 1)\,\mathbb{E}_{q(\boldsymbol{\Theta})}[\ln\pi_k] - (d_k - 1)\,\mathbb{E}_{q(\boldsymbol{\Theta})}[\ln\pi_k]
\end{aligned} \tag{47}$$

We can obtain the final simplified form of $\mathcal{L}^{\text{HDP}}$ by replacing each of the expectations with their closed-forms. Recall that $\mathbb{E}_{q(\boldsymbol{\Theta})}[\psi_n = k] = r_{nk}$ and $\mathbb{E}[\ln\pi_k] = \boldsymbol{\Psi}(d_k) - \boldsymbol{\Psi}(\sum_k d_k)$ from Eq. (19). Additionally, we found the form of $\mathbb{E}_{q(\boldsymbol{\Theta})}[\beta_k]$ in Eq. (40). Plugging these values into the above results in the following

$$\begin{aligned}
\mathcal{L}^{\text{HDP}} &= \sum_{i=1}^{N}\sum_{k=1}^{K+1} r_{nk}\left[\boldsymbol{\Psi}(d_k) - \boldsymbol{\Psi}\left(\sum_k d_k\right)\right] + \mathbb{E}_{q(\boldsymbol{\Theta})}\left\{\ln\left[\frac{\Gamma(\sum_k \alpha_0\beta_k)}{\prod_k\Gamma(\alpha_0\beta_k)}\right]\right\} - \ln\left[\frac{\Gamma(\sum_k d_k)}{\prod_k\Gamma(d_k)}\right] \\
&\quad + \sum_{k=1}^{K+1}\alpha_0\left[\frac{g_{1k}g_{2k}}{g_{1k}g_{2k} + (1-g_{1k})g_{2k}}\prod_{l=1}^{k-1}\frac{(1-g_{1l})g_{2l}}{g_{1l}g_{2l} + (1-g_{1l})g_{2l}} - d_k\right]\left[\boldsymbol{\Psi}(d_k) - \boldsymbol{\Psi}\left(\sum_k d_k\right)\right]
\end{aligned} \tag{48}$$

where $\boldsymbol{\Psi}(\cdot)$ is the digamma function. Note that we did not expand one term and this is because this expression does not have a closed-form. Instead, we use a surrogate bound from Hughes et al.[11] to approximate it (see the derivation of the updates for $g_{1k}$ and $g_{2k}$ in Section 2.2 for details). Altogether, this results in the third portion of the lower bound in Eq. (21).

##### 3.4 Derivation of the Cluster Assignment Entropy for the Likelihood

Next, we will expand the component of the lower bound that represents the entropy of the cluster assignment probabilities. Recall that in Eqs. (18) and (41) we defined

$$\mathcal{H}(\boldsymbol{\psi}) = -\mathbb{E}_{q(\boldsymbol{\Theta})}[\ln q(\boldsymbol{\psi} \,|\, \boldsymbol{r})] = -\mathbb{E}_{q(\boldsymbol{\Theta})}\left[\sum_{i=1}^{N}\sum_{k=1}^{K}[\psi_n = k]\,\ln r_{nk}\right]. \tag{49}$$

As we did in the subsections, we can distribute the expectation throughout the expression and use the fact that $E_{q(\boldsymbol{\Theta})}[\psi_n = k] = r_{nk}$ to get the following simplified form

$$\mathcal{H}(\boldsymbol{\psi}) = \sum_{i=1}^{N}\sum_{k=1}^{K} r_{nk}\,\ln r_{nk}. \tag{50}$$

which results in the fourth portion of the lower bound in Eq. (21).

##### 3.5 Likelihood Corresponding to the Upper Level of the Dirichlet Process

Finally, we can expand the partition of the lower bound that contains the upper level of the Dirichlet process. We follow the construction of Hughes et al.[11] and place a proper beta distribution as the variational approximating distribution for each $\chi_k$ in Eq. (20). This construction comes with the benefit of penalizing the allocation to empty clusters, thus allowing for faster convergence during model training. Working from the last component of Eq. (41), we get the following relationship

$$\begin{aligned}
\mathcal{L}^{\boldsymbol{\chi}} &= \mathbb{E}_{q(\boldsymbol{\Theta})}\left\{\ln\left[\frac{p(\boldsymbol{\chi} \,|\, \gamma_0)}{q(\boldsymbol{\chi} \,|\, \boldsymbol{g_1}, \boldsymbol{g_2})}\right]\right\} \\
&= \mathbb{E}_{q(\boldsymbol{\Theta})}[\ln p(\boldsymbol{\chi} \,|\, \gamma_0) - \ln q(\boldsymbol{\chi} \,|\, \boldsymbol{g_1}, \boldsymbol{g_2})] \\
&= \mathbb{E}_{q(\boldsymbol{\Theta})}\left[\sum_{k=1}^{K} -\ln\left(\frac{\Gamma(\gamma_0)\Gamma(1)}{\Gamma(\gamma_0+1)}\right) + (1-1)\,\ln\chi_k + (\gamma_0 - 1)\,\ln(1-\chi_k)\right] \\
&\quad - \mathbb{E}_{q(\boldsymbol{\Theta})}\left[\sum_{k=1}^{K} -\ln\left(\frac{\Gamma(g_{1k}g_{2k})\Gamma((1-g_{1k})g_{2k})}{\Gamma(g_{1k}g_{2k} + (1-g_{1k})g_{2k})}\right) + (g_{1k}g_{2k} - 1)\,\ln\chi_k + ((1-g_{1k})g_{2k} - 1)\,\ln(1-\chi_k)\right].
\end{aligned}$$

We can further simplify and reorder the summations of the beta priors such that they share the same summation over the number of cluster components $K$ where

$$\mathcal{L}^{\mathcal{X}} = \sum_{k=1}^{K} \left[ -\ln\left(\frac{\Gamma(\gamma_0)\Gamma(1)}{\Gamma(\gamma_0+1)}\right) + \ln\left(\frac{\Gamma(g_{1k}g_{2k})\Gamma((1-g_{1k})g_{2k})}{\Gamma(g_{1k}g_{2k}+(1-g_{1k})g_{2k})}\right)\right]$$

$$+ \sum_{k=1}^{K} \left\{ (\gamma_0 - 1)\,\mathbb{E}_{q(\boldsymbol{\Theta})}[\ln(1-\chi_k)] - (g_{1k}g_{2k}-1)\,\mathbb{E}_{q(\boldsymbol{\Theta})}[\ln\chi_k] + ((1-g_{1k})g_{2k}-1)\,\mathbb{E}_{q(\boldsymbol{\Theta})}[\ln(1-\chi_k)]\right\}$$

We can substitute $\mathbb{E}_{q(\boldsymbol{\Theta})}[\ln\chi_k] = \boldsymbol{\Psi}(g_{1k}g_{2k}) - \boldsymbol{\Psi}(g_{2k})$ and $\mathbb{E}_{q(\boldsymbol{\Theta})}[\ln(1-\chi_k)] = \boldsymbol{\Psi}((1-g_{1k})g_{2k}) - \boldsymbol{\Psi}(g_{2k})$. This results in the following simplified form

$$\begin{aligned} \mathcal{L}^{\mathcal{X}} &= -K\,\ln\left(\frac{\Gamma(\gamma_0)\Gamma(1)}{\Gamma(\gamma_0+1)}\right) + \sum_{k=1}^{K}\ln\left(\frac{\Gamma(g_{1k}g_{2k})\Gamma((1-g_{1k})g_{2k})}{\Gamma(g_{1k}g_{2k}+(1-g_{1k})g_{2k})}\right) \\ &\quad + \sum_{k=1}^{K}(\gamma_0-(1-g_{1k})g_{2k})\boldsymbol{\Psi}((1-g_{1k})g_{2k}) - \boldsymbol{\Psi}(g_{2k}) + (1-g_{1k}g_{2k})\boldsymbol{\Psi}(g_{1k}g_{2k})\boldsymbol{\Psi}(g_{2k}) \end{aligned} \tag{51}$$

which results in the fifth and final portion of the lower bound in Eq. (21).

### 4 Pseudocode for Variational EM Algorithm

**Algorithm 1** Nonparametric Clustering of Single Cell Populations (NCLUSION)

```
Input log-normalized single-cell RNA sequencing (scRNA-seq) expression data X.
Choose the number of maximum iterations T and tolerance parameter ϵ.
Choose the number of possible clusters K, and set the concentration hyper-parameters α_0 and γ_0.
Randomly initialize variational parameters {y_jk, r_nk, d_k, g_1k, g_2k, h_1, h_2, m_jk, s^2_jk, f_j, t^2_j, u_jk, v_jk, a_j, b_j}
    for i = 1, ..., N cells and j = 1, ..., J genes across K clusters, respectively.
Compute initial value of evidence lower bound (ELBO).
for i = 1 → T do
    Set ELBO = ELBO_new.
    Update {y_jk, r_nk}_{k=1}^K.                                                  ▷ E-Step
    Update {m_jk, s^2_jk, u_jk, v_jk, d_k, g_1k, g_2k}_{k=1}^K, f_j, t^2_j, h_1, h_2, a_j, and b_j.   ▷ M-Step
    Update lower bound ELBO_new.
    if ELBO_new − ELBO ≤ ϵ then
        Save ELBO = ELBO_new.
        Break
    end if
end for
Get K* ≤ K by dropping indices where the number of cells occupying the k-th cluster N_k = 0.
Compute the cluster assignment labels for each cell k*_n = max(r_n1, ..., r_nK*).
Compute marginal and adjusted posterior inclusion probabilities PIP(j; k) and PIP*(j; k).
Compute the effect size sign ESS(j; k) and strictly standardized mean difference SSMD(j; k) based
    on posterior mean μ̂_jk = y_jk m_jk and posterior precision 1/σ̂^2_j = a_j/b_j for each gene.
Return {k*_n, PIP(j; k), PIP*(j; k), ESS(j; k), SSMD(j; k)}.
```

### 5 Details on Comparisons between Clustering Algorithms

In the main text, we compare the performance of NCLUSION against five baseline methods. We give short descriptions of how we implement these approaches below.

#### 5.1 Competing Methods for Cluster Identification

To evaluate clustering accuracy for NCLUSION, we compared it to a wide range of both established and recently proposed unsupervised learning algorithms. Each method was applied to the same synthetic input data where we had 20 replicates per simulation scenario. Clustering performance was evaluated by the comparing inferred labels to known cell type annotations using the adjusted Rand index (ARI) and normalized mutual information (NMI) as evaluation metrics (see the main text for details).

**CIDR.** The "Clustering through Imputation and Dimensionality Reduction" (CIDR) approach was applied to raw count matrices. Dropout candidates were identified and removed, and dissimilarity-based clustering was performed on principal components determined via an internal heuristic specified by Lin et al.[31]. The final cluster labels were obtained via the `scCluster` function from CIDR package.

**SC3.** The SC3 algorithm was applied to log-normalized expression counts with an internal estimation of the optimal number of clusters found via the `sc3_estimate_k` function from Kiselev et al.[32]. Since our simulations consisted of datasets with more than 5,000 cells, a support vector machine (SVM)-based extension was used to scale clustering. Only unique genes were retained and default gene filtering was enabled.

**scCCESS+SIMLR.** To run the "single-cell Consensus Clusters of Encoded Subspaces" (scCCESS-SIMLR) approach, we first estimated the number of clusters with the `estimate_k` function in the software by searching over the default range of $K = 5$ to 15. Once this value was obtained, we then used the `SIMLR_Large_Scale` function to assign cells to each cluster following the suggested parameter values described in Yu et al.[33] and Wang et al.[34].

**scLCA.** To run the "single-cell latent cellular analysis" (scLCA), we followed the documentation in Cheng et al.[35]. As required by the corresponding software package, we ran experiments on scLCA using the raw expression counts of some preselected subset of highly variable genes. For the training set, we opted to use 10% of the total number of cells in the study instead of using the default 1000 cells. This greatly improved the performance of the method. We limited the training set size to 10% due to the significantly longer run times for larger proportions. We used the default max number of clusters ($K = 10$) and the remaining parameters were also used at their default values.

**KNN+Leiden.** Here, we used the K-nearest Neighbor (KNN) classifier built within the `scikit-learn` library[36] to construct a sparse adjacency matrix of k-neighbor connectivities (i.e., distances). We performed a small grid search with $K$ ranging from 2 to 25 to find the optimal number of neighbors. Optimal neighbors were determined using `scikit-learn`'s `GridSearchCV` function. Briefly, for each neighbor on the grid, this function performs M-fold cross-validation by training a classifier to predict the accuracy on the held-out set. In the text, we used M=5. The neighbors with the highest accuracy are selected and used to cluster cells with the Leiden algorithm in `scanpy`[37]. All of the remaining parameters in `scanpy` were set to their default values.

**Seurat.** As required by the Seurat software package[38–41], preprocessed single-cell RNA sequencing data was scaled such that the expression of each gene has zero mean and unit variance. All principal component analyses (PCA) were performed on some subset of pre-selected top highly-variable genes. We obtained results in the main text by running the software's nearest-neighbor graph construction and cluster assignment method following the package tutorial (see `https://satijalab.org/seurat/articles/pbmc3k_tutorial.html`).

**SOUP.** To implement the "semisoft clustering with pure cells" (SOUP) approach, we followed the software documentation from Zhu et al.[42] by using preprocessed single-cell RNA sequencing data as input. By following the vignette provided in the software package, we performed a small grid search with $K$ ranging from 2 to 5 to find the optimal number of neighbors. We found that increasing the upper bound on this grid led to prohibitively long run times and expensive use of computational resources. The remaining parameters were set to their default values.

**scDeepCluster.** The scDeepCluster approach by Tian et al.[43] is a deep-learning-based method which uses an autoencoder with self-training to refine clustering on latent representations. Here, raw expression counts were normalized, size-factor adjusted, and log-transformed before model training. An initial clustering estimate was obtained using the Louvain algorithm on the autoencoder's latent space. These estimates were used to initialize a deep clustering model trained with KL divergence minimization and a target distribution update strategy. Final cluster assignments were extracted after model convergence.
Lastly, we want to note that for graph-based methods (e.g., algorithms using Leiden and scDeepCluster), neighborhood graphs were computed using either ($i$) pre-specified hyperparameters according to software

default settings or ($ii$) optimized hyperparameters selected as part of the general software implementation.

#### 5.2 Competing Methods for Marker Gene Selection

To evaluate the ability of NCLUSION to detect the correct marker genes, we compared it to a wide range of gene selection algorithms. Across all simulation scenario, each method configured to select the top 500 most informative features unless otherwise specified. Detected genes were ranked according to each method-specific significance scores and performance was assessed by comparing the ranked gene lists to the known marker gene sets. The following marker gene selections algorithms were compared in our analyses:

**DUBStepR.** The "Determining the Underlying Basis using Stepwise Regression" (DUBStepR) framework by Ranjan et al.[44] identifies marker genes based on differential correlation patterns in the local structure of a PCA-derived cell neighborhood graph. Briefly, DUBStepR carries out feature selection by leveraging gene-gene relationships with a measure of inhomogeneity in transcriptomic space, termed the Density Index (DI). For each dataset, we applied the DUBStepR to log-normalized and scaled expression matrices after subsetting the data to a select number of highly variable genes. Where available, PCA coordinates from `scanpy` were reused as embeddings in the model to preserve consistency across workflows. Marker genes were ranked by these DI correlation range scores, with missing genes assigned NA (or zero) significance values to preserve completeness.

**FESTEM.** The "Feature Selection Through EM" (FESTEM) algorithm by Chen et al.[45] is an approach that is meant to identify informative genes from cell type distributions via a deterministic EM hypothesis testing framework. In practice, it runs as part of a two-step workflow where FESTEM first identifies candidate top genes. Then, in the second step, these top genes are used for clustering with Seurat. For each gene in turn, Festem performs a statistical test (an EM test for negative binomial distributions) to determine if its expression is homogeneously distributed (i.e., not a marker gene) or heterogeneously distributed (i.e., a marker gene). It then assigns a p-value based on a chi-squared distribution with three degrees of freedom. The p-values are adjusted for multiple comparisons using the Benjamini-Hochberg procedure, and the top genes are selected for clustering. It is important to note that FESTEM does not identify cluster-specific marker genes themselves, just candidate genes that are most likely to be.

In other words, it might identify that gene $A$ has a heterogenous distribution across different cells but cannot identify for which cell type(s) it is marker. Therefore, it does not provide the same resolution as NCLUSION. Still, we include it in our comparisons since our marker detection assessment was evaluated globally (i.e., across all clusters).

**singleCellHaystack.** The singleCellHaystack approach was applied using two configurations from Vandenbon and Diez[46]: one based on low-dimensional UMAP coordinates (`singleCellHaystack_UMAP`) and another using PCA embeddings (`singleCellHaystack_PCA`).Regardless of the strategy, this general approach uses Kullback–Leibler divergence to find genes that are expressed in subsets of cells that are non-randomly positioned in a multidimensional space. Here, the distribution of expression for cells from a dataset is compared to a reference distribution. From this comparison, the KL-divergence of each gene is calculated and compared with permuted data to evaluate statistical significance. Marker genes are then identified by taking the top ranked adjusted log p-values.

All competing methods were run in `R` using standardized wrappers that read input from H5AD-formatted AnnData objects and wrote ranked feature tables to structured output files. As previously mentioned, missing genes were assigned NA values and included in output to facilitate alignment across method outputs. Where relevant, consistent preprocessing (e.g., Seurat normalization, scaling, and PCA computation) was enforced. Marker detection was evaluated globally (i.e., across all clusters) using power (i.e., true positive rate), false discovery rate (FDR), and false positive rate (i.e., 1-Specificity) as metrics.

### 6 Description of Computing Resources

All timing experiments were performed using computing resources at Microsoft Research. This consisted of a 64-bit AMD EPYC 7742 64-Core Processor (x86_64 architecture) with 128 threads. The amount of memory was automatically requested and the total amount used varied based on the size of the dataset being analyzed. All other experiments were run on a 64-bit Intel(R) Xeon(R) Platinum 8268 24-Core CPU @ 2.90GHz (x86_64 architecture) with 48 threads. Across all datasets, 128 gigabytes (GB) of total memory was requested and the amount used varied based on the size of the dataset being analyzed.

### 7 Supplementary Figures

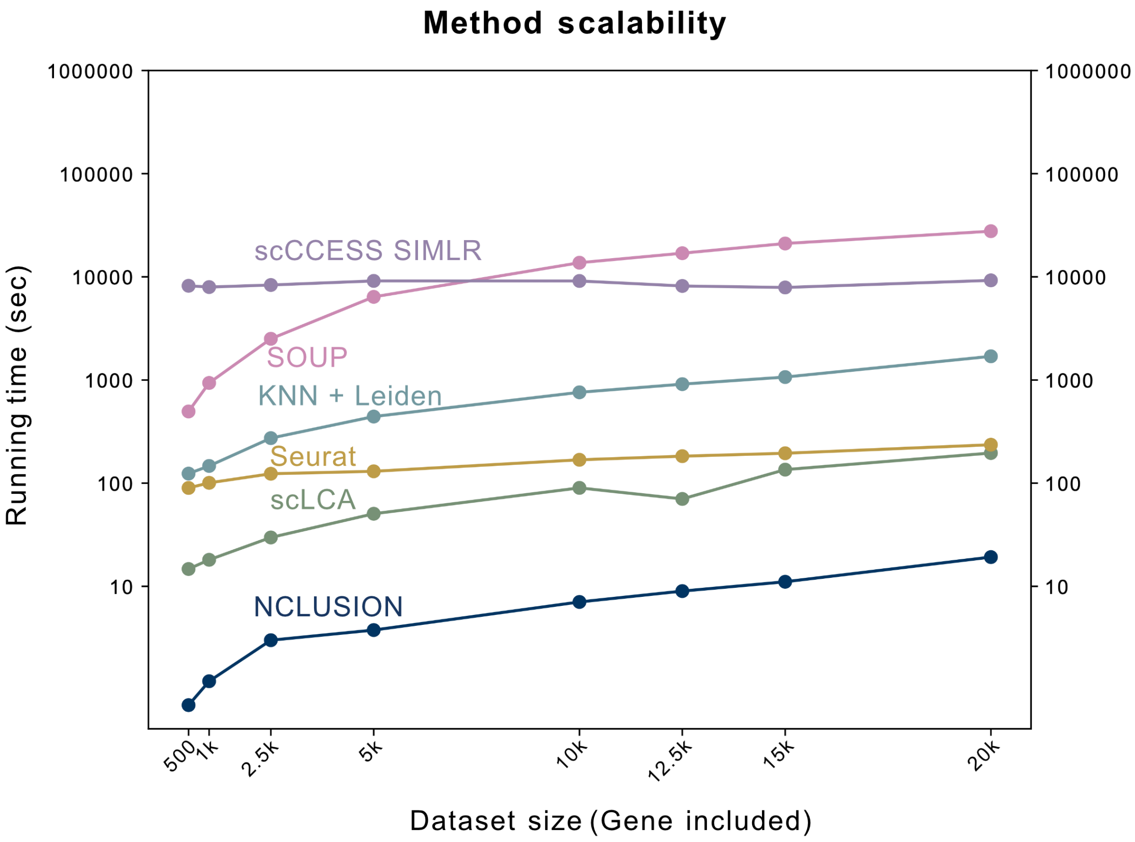

**Fig S1. Runtimes of NCLUSION and other clustering baselines on the BRAIN-LARGE data with a fixed (subsampled) set of $N = 31{,}000$ cells and a varying number of variables ranging from $J = 500$ to 20,000 genes.** The methods used as comparisons include: Seurat, scLCA, K-nearest neighbors followed by the Leiden clustering algorithm (KNN+Leiden), SOUP, and scCCESS-SIMLR. All runtimes are recorded in seconds.

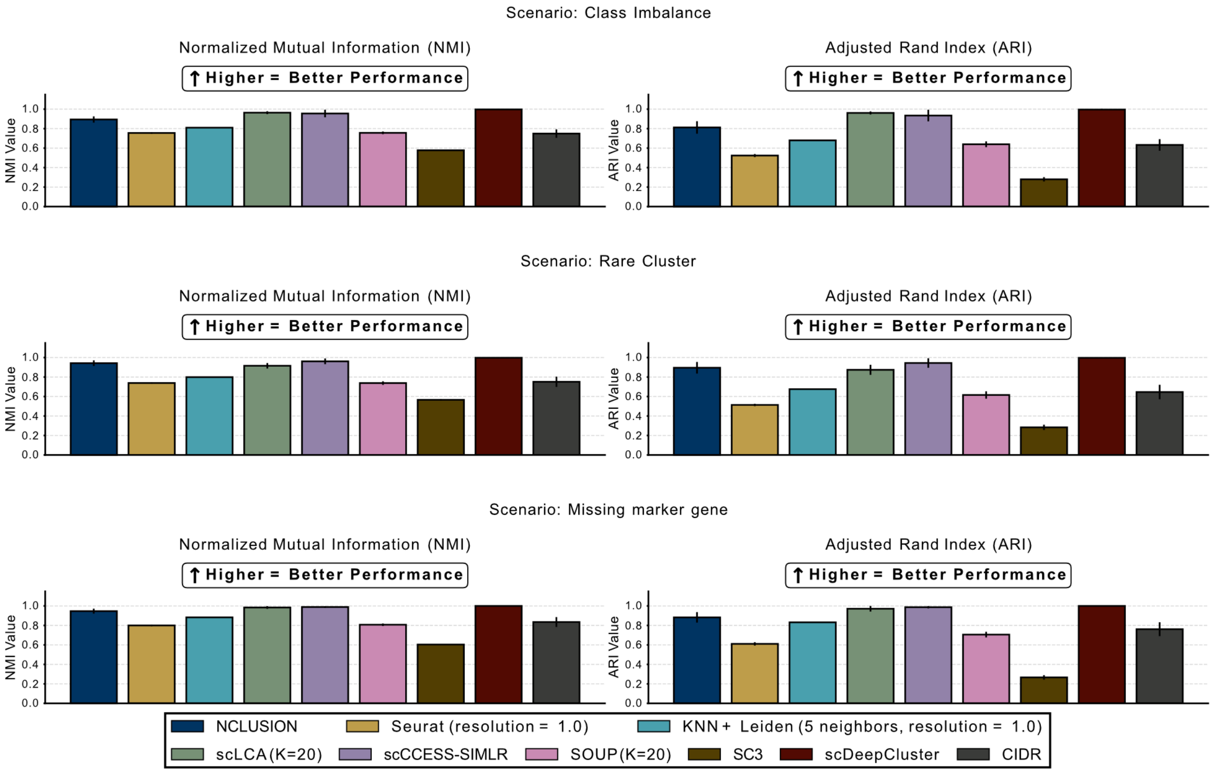

**Fig S2. Comparing NCLUSION and competing algorithms on performing clustering in a simulation study.** Depicted are results for Scenarios II-IV for the simulation study. Each simulated dataset comprised of $N = 10{,}000$ cells and $J = 1{,}000$ genes where we preserved realistic transcriptomic correlation structures through Gaussian copula modeling with `scDesign3`[47]. In Scenario II (top), we implemented an imbalanced cluster design where one small cluster had 200 cells and the other four larger clusters each had 2450 cells. In Scenario III (middle), there was also an imbalanced cluster design but a situation where one cluster had 20 (rare) cells and the other four larger clusters each had 2495 cells each. All clusters in Scenarios II and III had 50 unique marker genes each. Lastly, in Scenario IV (bottom), we generated balanced clusters of 2000 cells per cell type, but one cluster had only 20 marker genes while the other four clusters had 50 marker genes. Inferred clusters labels were compared to "true" annotations created during the simulation, where performance was measured according to (left) normalized mutual information (NMI) and (right) adjusted Rand index (ARI). We compare NCLUSION to Seurat, K-nearest neighbors followed by the Leiden clustering algorithm (KNN+Leiden), scLCA, scCCESS-SIMLR, SOUP, SC3, scDeepCluster, and CIDR. Results are based on 20 simulations per simulation scenario, with each bar plot representing the mean and the error bars covering a ± 95% confidence interval.

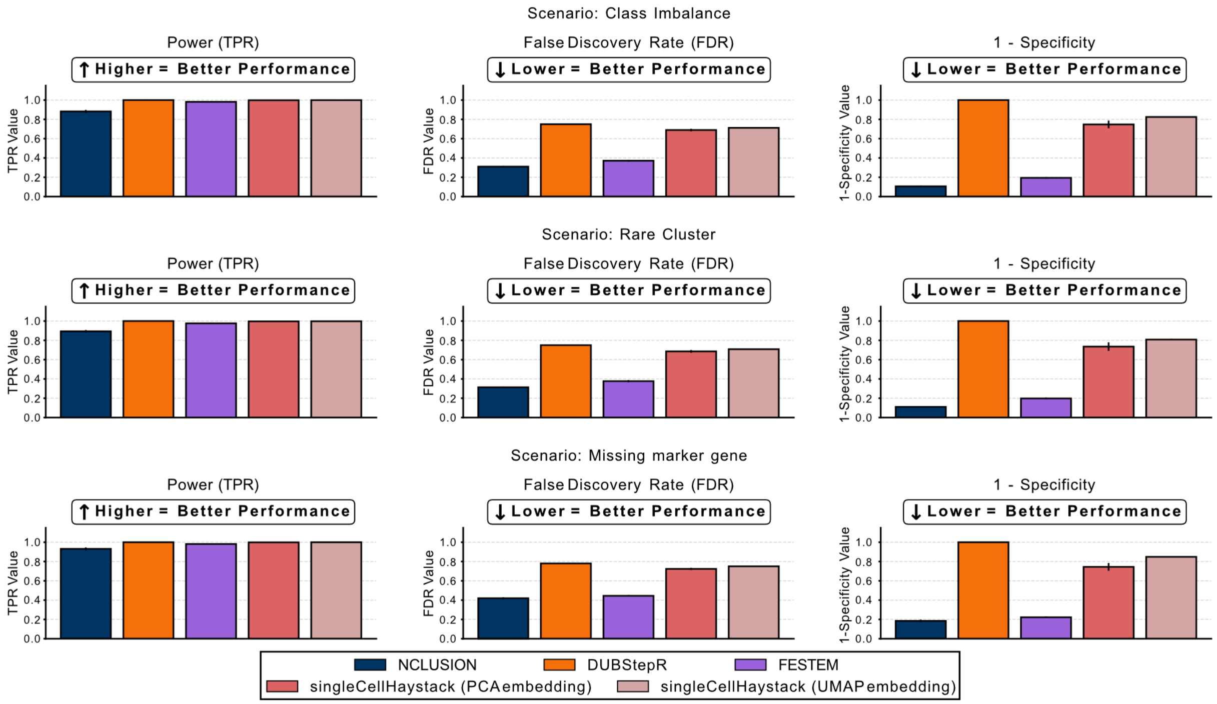

**Fig S3. Comparing NCLUSION and competing algorithms on performing gene selection in a simulation study.** Depicted are results for Scenarios II-IV for the simulation study. Each simulated dataset comprised of $N = 10{,}000$ cells and $J = 1{,}000$ genes where we preserved realistic transcriptomic correlation structures through Gaussian copula modeling with `scDesign3`[47]. In Scenario II (top), we implemented an imbalanced cluster design where one small cluster had 200 cells and the other four larger clusters each had 2450 cells. In Scenario III (middle), there was also an imbalanced cluster design but a situation where one cluster had 20 (rare) cells and the other four larger clusters each had 2495 cells each. All clusters in Scenarios II and III had 50 unique marker genes each. Lastly, in Scenario IV (bottom), we generated balanced clusters of 2000 cells per cell type, but one cluster had only 20 marker genes while the other four clusters had 50 marker genes. Assessment of marker gene selection was done on the global scale—meaning, we evaluated how well a method was able to detect a "true" causal gene without taking cluster assignment into account. This was due to the limitation of competing methods not being able to identify cluster-specific genes. Evaluations were done by measuring the true positive rate (TPR; or power), false discovery rate (FDR), and false positive rate (FPR; computed as 1-Specificity) for each approach. We compare NCLUSION to DUBStepR, FESTEM, singleCellHaystack (one version using a PCA embedding and the other using a UMAP embedding). Results are based on 20 simulations per simulation scenario, with each bar plot representing the mean and the error bars covering a ± 95% confidence interval.

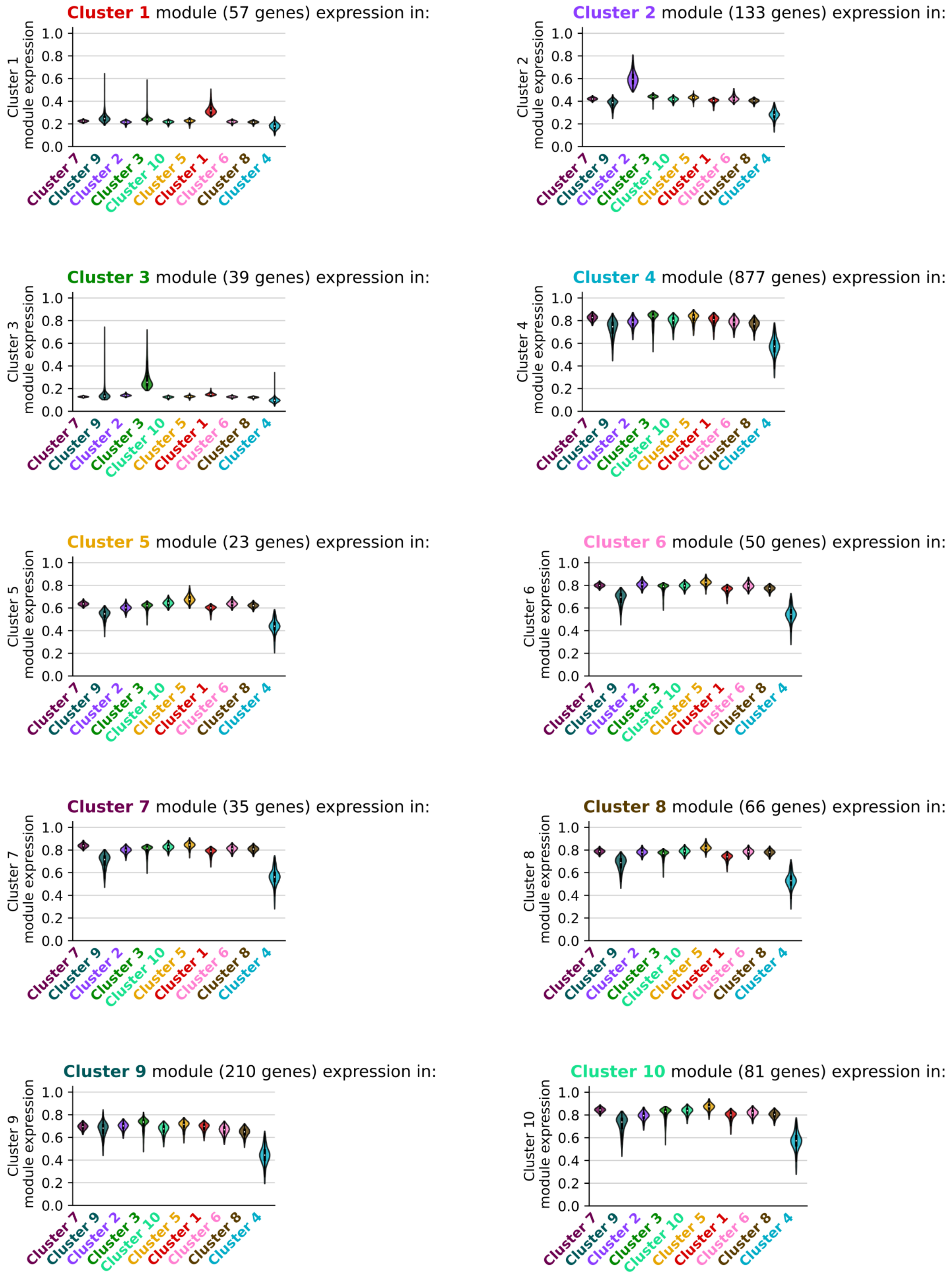

**Fig S4. Violin plots comparing the normalized expression of cluster-specific marker genes across clusters inferred by NCLUSION in the PBMC dataset ($N = 94{,}615$ cells).** Each panel represents an evaluation in a given inferred cluster. The objective of this analysis is to see that each gene module exhibits the highest expression within its respective cluster relative to others.

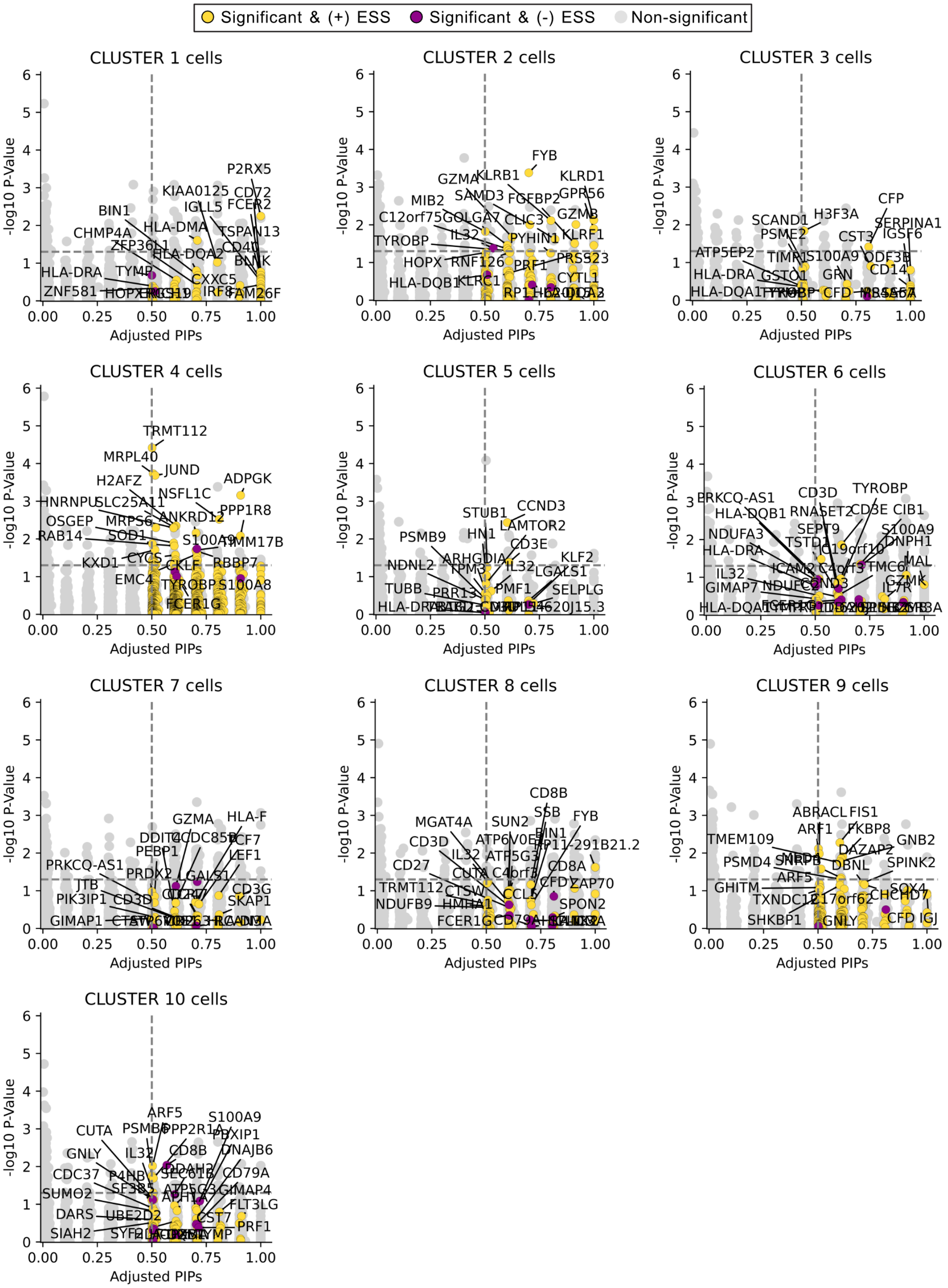

**Fig S5. Scatter plot comparing the marker genes identified using *post hoc* differential expression analysis with Seurat versus the variable selection approach with NCLUSION in the PBMC dataset ($N$ = 94,615 cells).** Here, our evaluation is based on NCLUSION's inferred cluster labels (depicted in each panel). The y-axis shows log-transformed $q$-values (i.e., adjusted $P$) for differentially expressed genes from Seurat found via a Wilcoxon rank sum test which is performed by doing a *post hoc* one-versus-all comparison for each cluster. The x-axis shows the adjusted posterior inclusion probabilities (PIPs) for each gene as computed by NCLUSION. All points in color are genes with PIP $\geq 0.5$. The specific color corresponds to the effect size sign (ESS) for each gene. Yellow points are genes with ESS = + (up-regulated), while purple points are genes with ESS = - (down-regulated). The vertical dashed line marks the median probability criterion[12]. The horizontal dashed line marks the Bonferroni-corrected threshold corresponding to significant $q$-values $\leq 0.05$. Genes found above the horizontal line and to the right of the vertical line are selected by both approaches; while, elements in the bottom right and top left quadrants are uniquely identified by NCLUSION and Seurat, respectively.

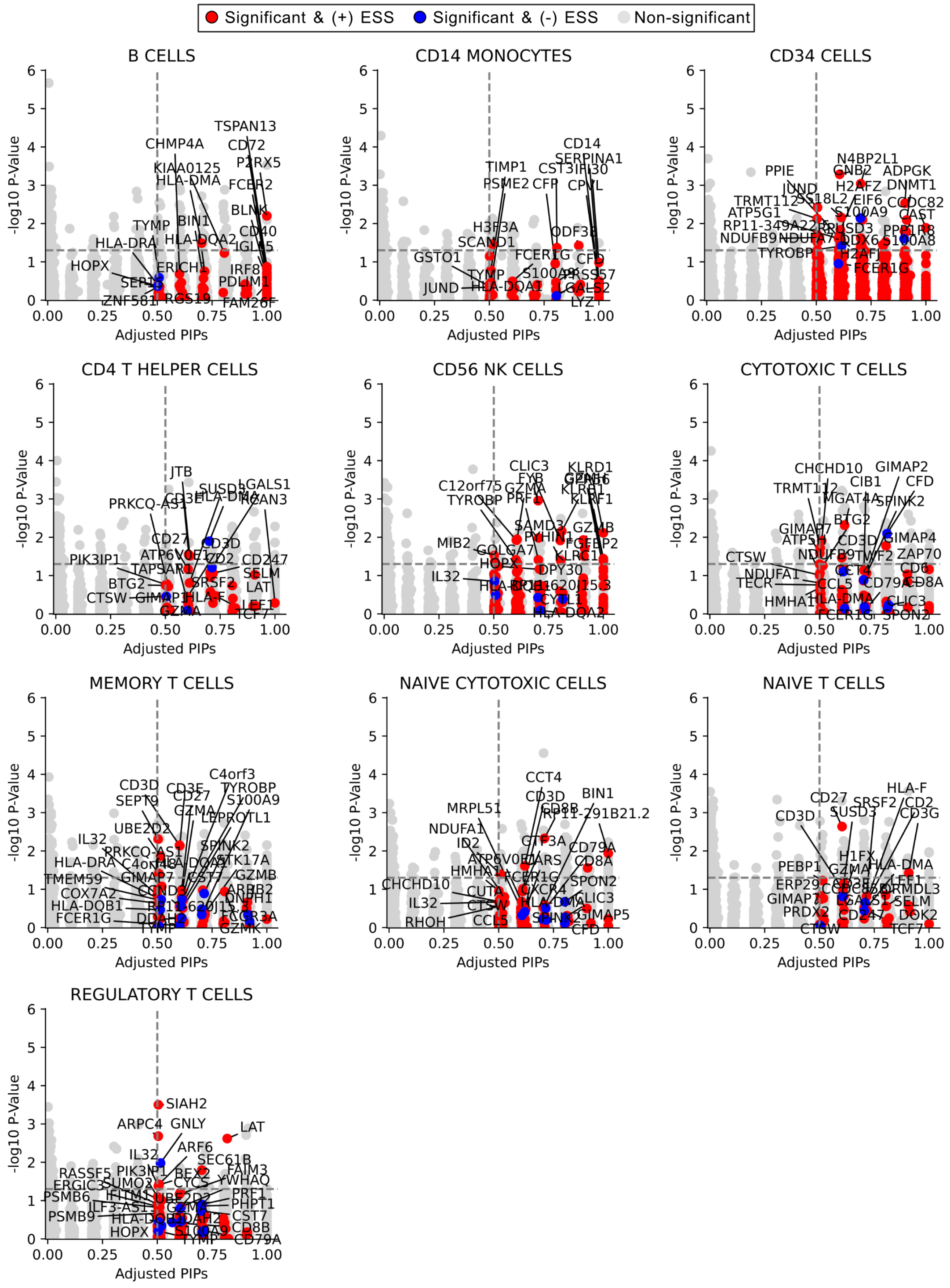

**Fig S6. Scatter plot comparing the marker genes identified using *post hoc* differential expression analysis with Seurat versus the variable selection approach with NCLUSION in the PBMC dataset ($N$ = 94,615 cells).** Here, we evaluate clusters according to the FACS-derived experimental annotations from Zheng et al.[48]. The y-axis shows log-transformed $q$-values (i.e., adjusted $P$) for differentially expressed genes from Seurat found via a Wilcoxon rank sum test which is performed by doing a *post hoc* one-versus-all comparison for each cluster. The x-axis shows the adjusted posterior inclusion probabilities (PIPs) for each gene as computed by NCLUSION. All points in color are genes with PIP $\geq 0.5$. The specific color corresponds to the effect size sign (ESS) for each gene. Red points are genes with ESS = + (up-regulated), while blue points are genes with ESS = - (down-regulated). The vertical dashed line marks the median probability criterion[12]. The horizontal dashed line marks the Bonferroni-corrected threshold corresponding to significant $q$-values $\leq 0.05$. Genes found above the horizontal line and to the right of the vertical line are selected by both approaches; while, elements in the bottom right and top left quadrants are uniquely identified by NCLUSION and Seurat, respectively.

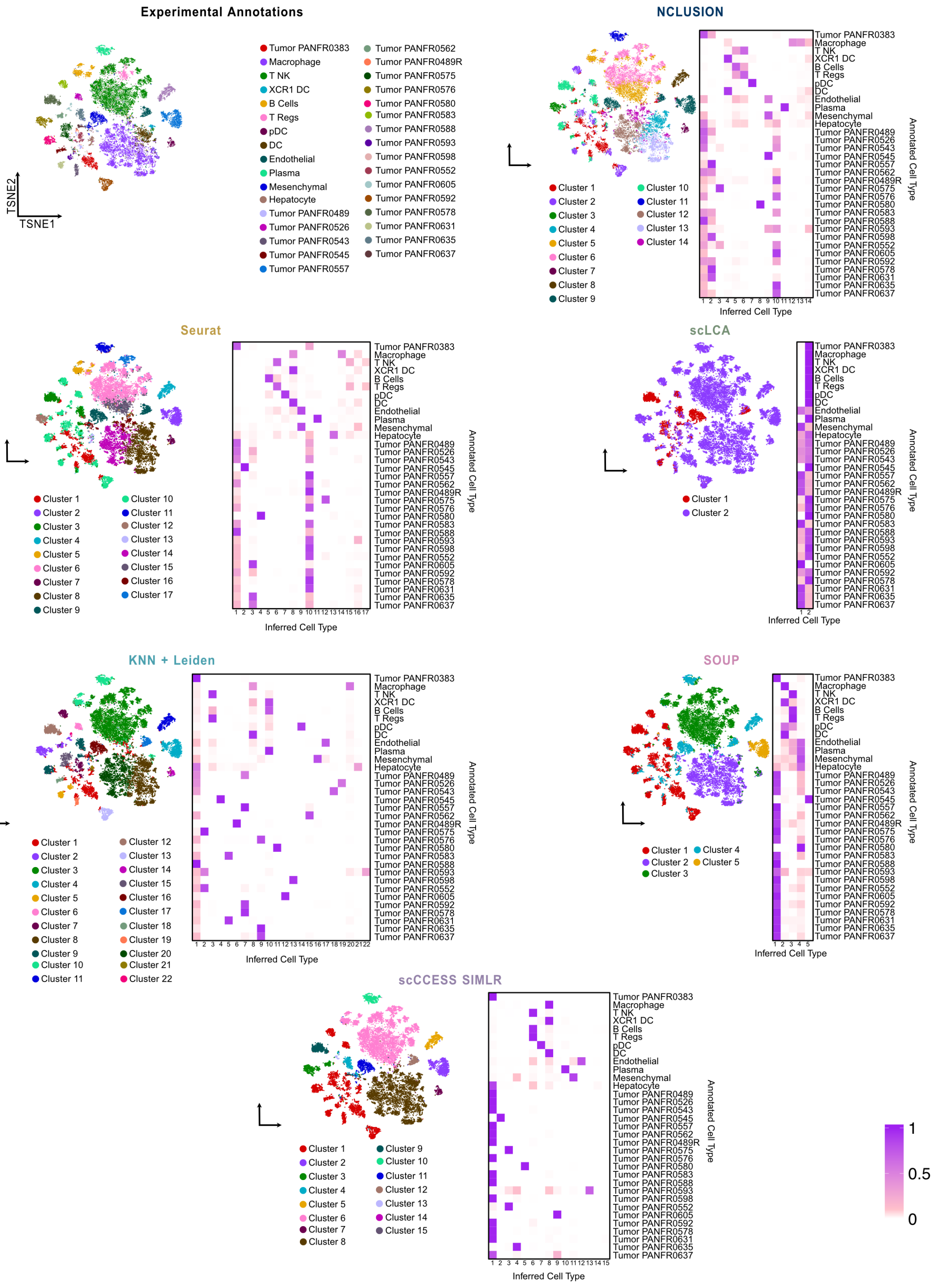

**Fig S7. Comparing the quality of clustering for NCLUSION and other baseline methods on the PDAC dataset ($N$ = 23,042 cells).** The first panel in the top left corner shows an overview of the cellular annotations found in the original study by Raghavan et al.[49]. The other panels depict results from running NCLUSION and other baseline methods on the same data. These baselines include: Seurat, scLCA, K-nearest neighbors followed by the Leiden clustering algorithm (KNN+Leiden), SOUP, and scCCESS-SIMLR, respectively. Here, we visualize the structure of the inferred clusters across all baselines using t-distributed stochastic neighbor embeddings (t-SNEs) on the left hand side of each panel and a contingency heat map showing the prevalence of each cell type across clusters on the right hand side.

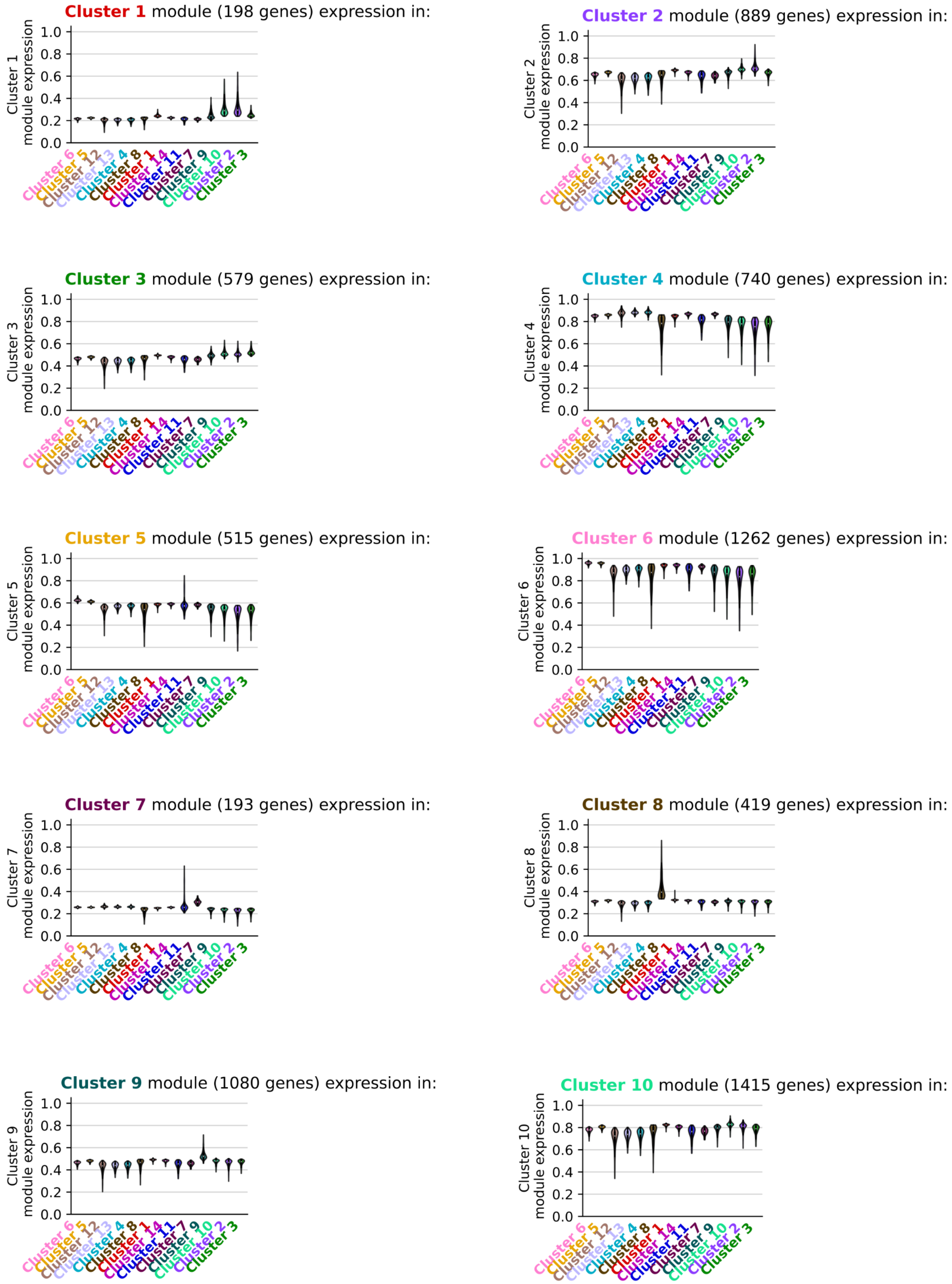

**Fig S8. Violin plots comparing the normalized expression of cluster-specific marker genes across clusters inferred by NCLUSION in the PDAC dataset ($N$ = 23,042 cells).** Each panel represents an evaluation in a given inferred cluster. The objective of this analysis is to see that each gene module exhibits the highest expression within its respective cluster relative to others. Results in this figure are continued in Fig S9 and they are related to results shown in Fig 5D-G.

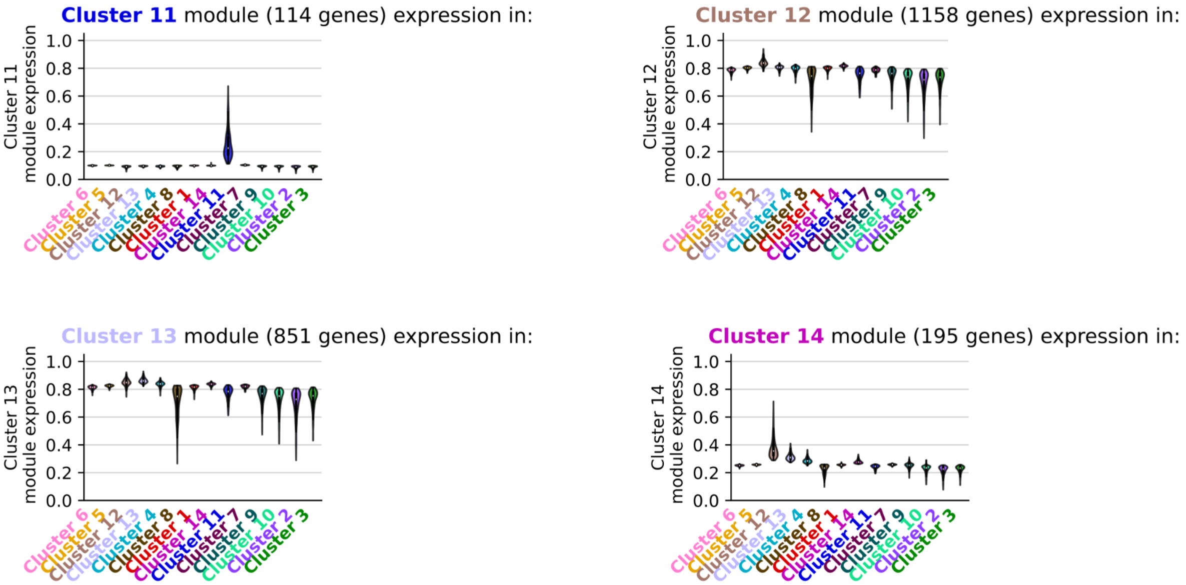

**Fig S9. Violin plots comparing the normalized expression of cluster-specific marker genes across clusters inferred by NCLUSION in the PDAC dataset ($N$ = 23,042 cells).** Each panel represents an evaluation in a given inferred cluster. The objective of this analysis is to see that each gene module exhibits the highest expression within its respective cluster relative to others. Results in this figure are an extension to the findings in Fig S8 and they are related to results shown in Fig 5D-G.

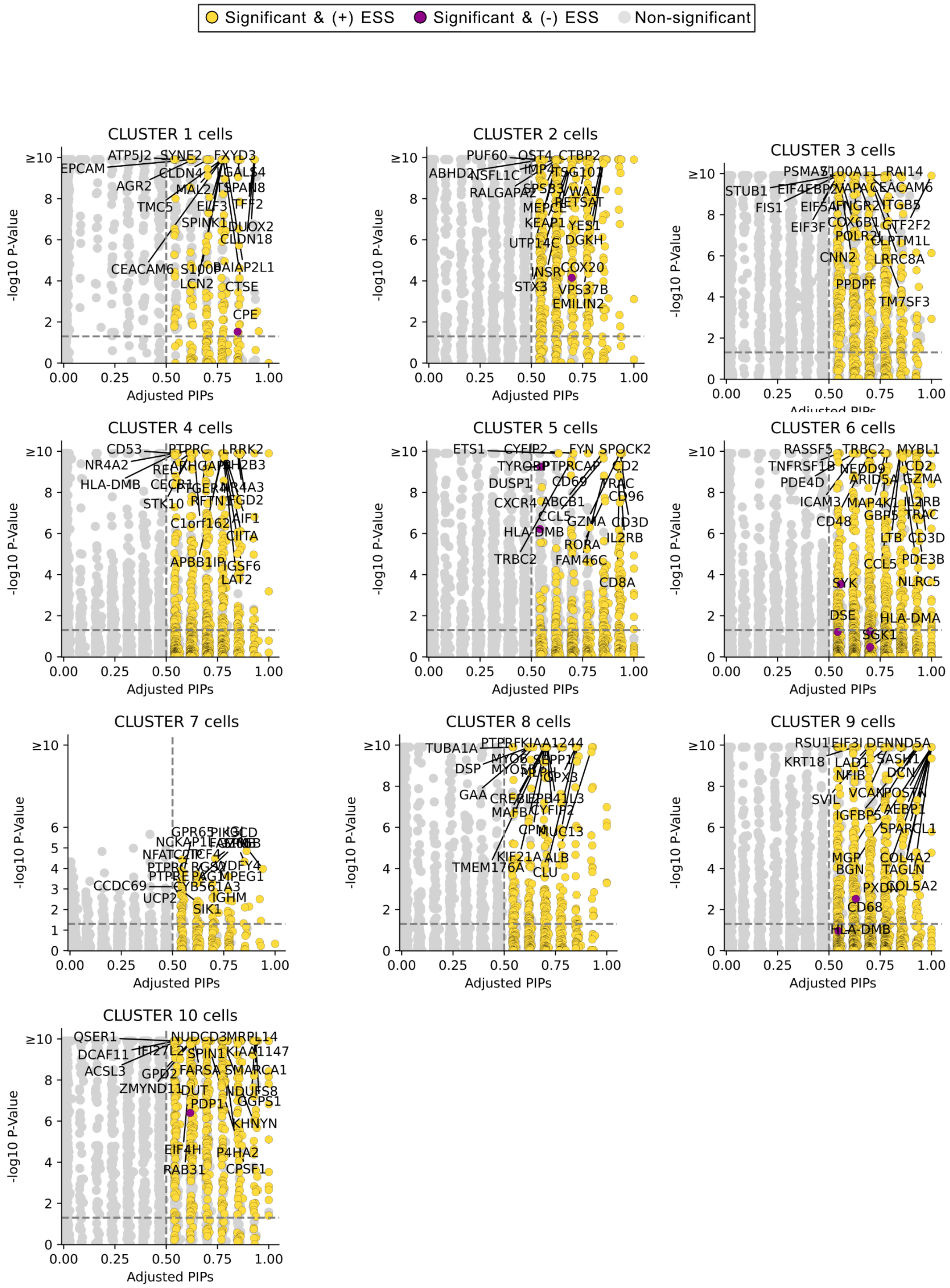

**Fig S10. Scatter plot comparing the marker genes identified using *post hoc* differential expression analysis with Seurat versus the variable selection approach with NCLUSION in the PDAC dataset ($N$ = 23,042 cells).** Here, our evaluation is based on NCLUSION's inferred cluster labels (depicted in each panel). The y-axis shows log-transformed $q$-values (i.e., adjusted $P$) for differentially expressed genes from Seurat found via a Wilcoxon rank sum test which is performed by doing a *post hoc* one-versus-all comparison for each cluster. The x-axis shows the adjusted posterior inclusion probabilities (PIPs) for each gene as computed by NCLUSION. All points in color are genes with PIP $\geq 0.5$. The specific color corresponds to the effect size sign (ESS) for each gene. Yellow points are genes with ESS = + (up-regulated), while purple points are genes with ESS = - (down-regulated). The vertical dashed line marks the median probability criterion[12]. The horizontal dashed line marks the Bonferroni-corrected threshold corresponding to significant $q$-values $\leq 0.05$. Genes found above the horizontal line and to the right of the vertical line are selected by both approaches; while, elements in the bottom right and top left quadrants are uniquely identified by NCLUSION and Seurat, respectively. Results in this figure are continued in Fig S11 and they are related to results shown in Fig 5D-G.

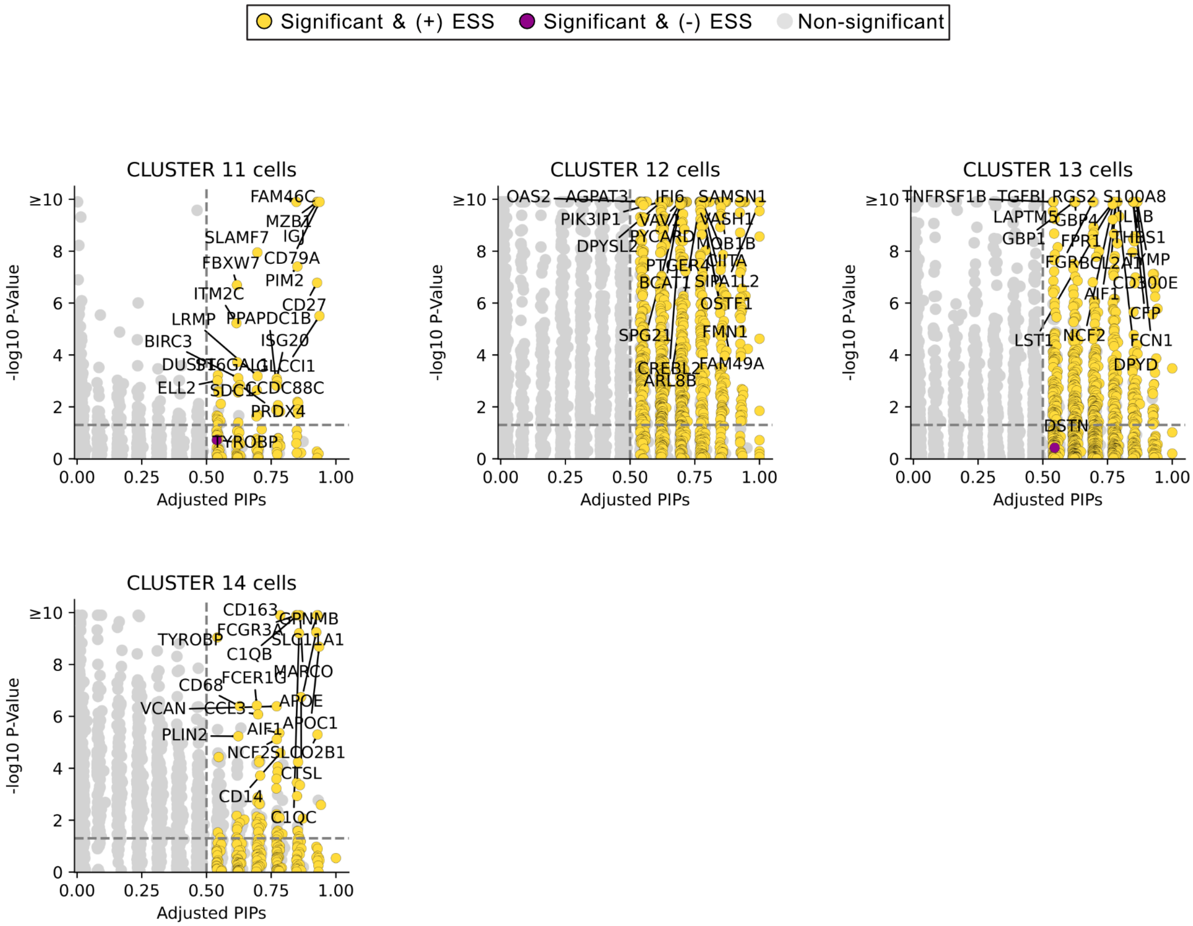

**Fig S11. Scatter plot comparing the marker genes identified using *post hoc* differential expression analysis with Seurat versus the variable selection approach with NCLUSION in the PDAC dataset ($N = 23{,}042$ cells).** Here, our evaluation is based on NCLUSION's inferred cluster labels (depicted in each panel). The y-axis shows log-transformed $q$-values (i.e., adjusted $P$) for differentially expressed genes from Seurat found via a Wilcoxon rank sum test which is performed by doing a *post hoc* one-versus-all comparison for each cluster. The x-axis shows the adjusted posterior inclusion probabilities (PIPs) for each gene as computed by NCLUSION. All points in color are genes with PIP $\geq 0.5$. The specific color corresponds to the effect size sign (ESS) for each gene. Yellow points are genes with ESS = + (up-regulated), while purple points are genes with ESS = - (down-regulated). The vertical dashed line marks the median probability criterion[12]. The horizontal dashed line marks the Bonferroni-corrected threshold corresponding to significant $q$-values $\leq 0.05$. Genes found above the horizontal line and to the right of the vertical line are selected by both approaches; while, elements in the bottom right and top left quadrants are uniquely identified by NCLUSION and Seurat, respectively. Results in this figure are an extension to the findings in Fig S10 and they are related to results shown in Fig 5D-G.

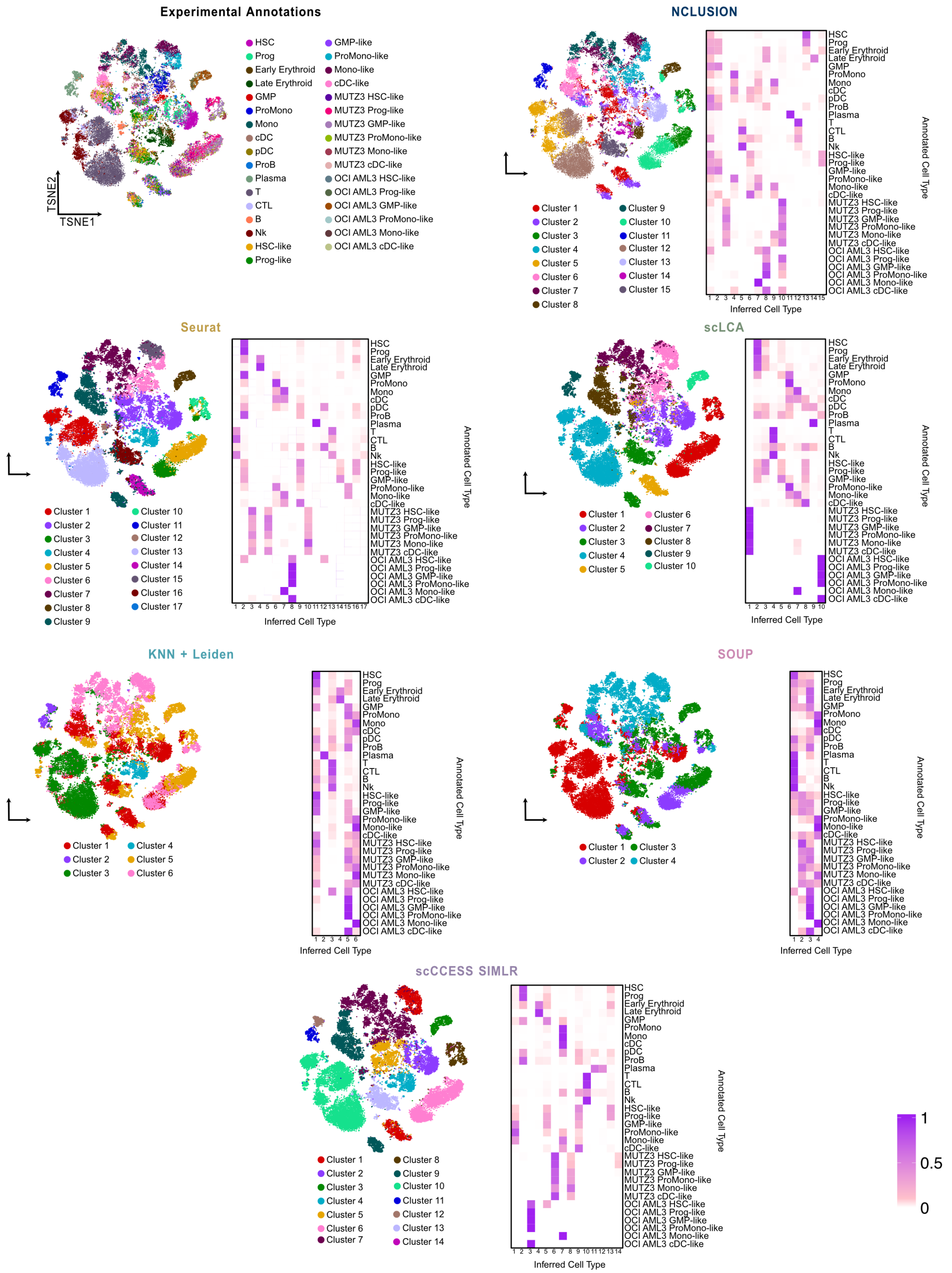

**Fig S12. Comparing the quality of clustering for NCLUSION and other baseline methods on the IMMUNE dataset ($N$ = 43,690 cells).** The first panel in the top left corner shows an overview of the cellular annotations found in the original study by van Galen et al.[50]. The other panels depict results from running NCLUSION and other baseline methods on the same data. These baselines include: Seurat, scLCA, K-nearest neighbors followed by the Leiden clustering algorithm (KNN+Leiden), SOUP, and scCCESS-SIMLR, respectively. Here, we visualize the structure of the inferred clusters across all baselines using t-distributed stochastic neighbor embeddings (t-SNEs) on the left hand side of each panel and a contingency heat map showing the prevalence of each cell type across clusters on the right hand side.

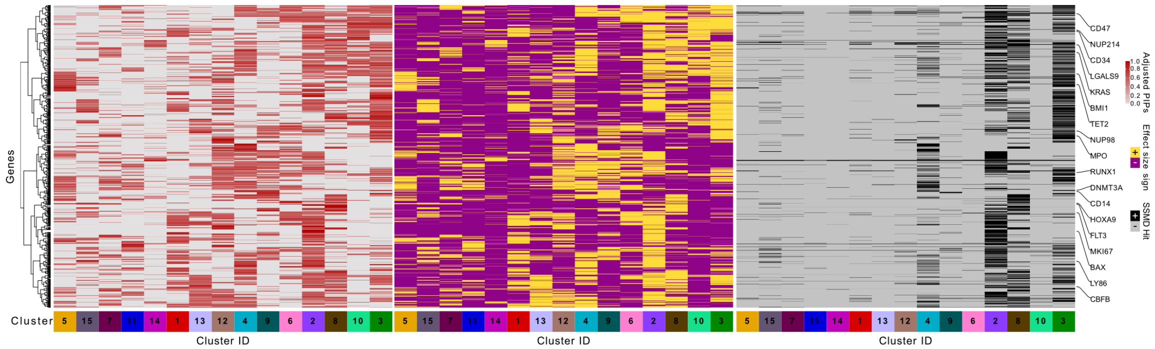

**Fig S13. Heat map of the significant genes as determined by (left) adjusted posterior inclusion probabilities (PIPs), (center) effect size sign (ESS), and (right) strictly standardized mean difference (SSMD) in each inferred cluster from NCLUSION for the AML dataset ($N$ = 43,690 cells). (Left)** Genes that are identified as "important" across many different clusters are seen as ubiquitous housekeeping variables rather than significant marker genes of unique cell types. Therefore, we down-weight the inclusion probabilities to proportionally penalize genes based on the number of clusters in which they appear. Here, significant genes are selected as those with adjusted PIPs greater than or equal to 0.5 which corresponds to the median probability criterion [12]. **(Center)** The ESS for each gene is computed by taking the sign of Cohen's $d$[13] between the expression of the $j$-th gene for cells in the $k$-th cluster and cells not in the $k$-th cluster. A positive ESS can be interpreted as the $j$-th gene being up-regulated in a given cluster. **(Right)** Significant genes are selected as those with SSMD greater than or equal to 0.15 which corresponds to the excluding genes with effect sizes that are weak [17]. In general, we observed that our criteria of cluster-specific marker genes was positively correlated with the expression of each gene.

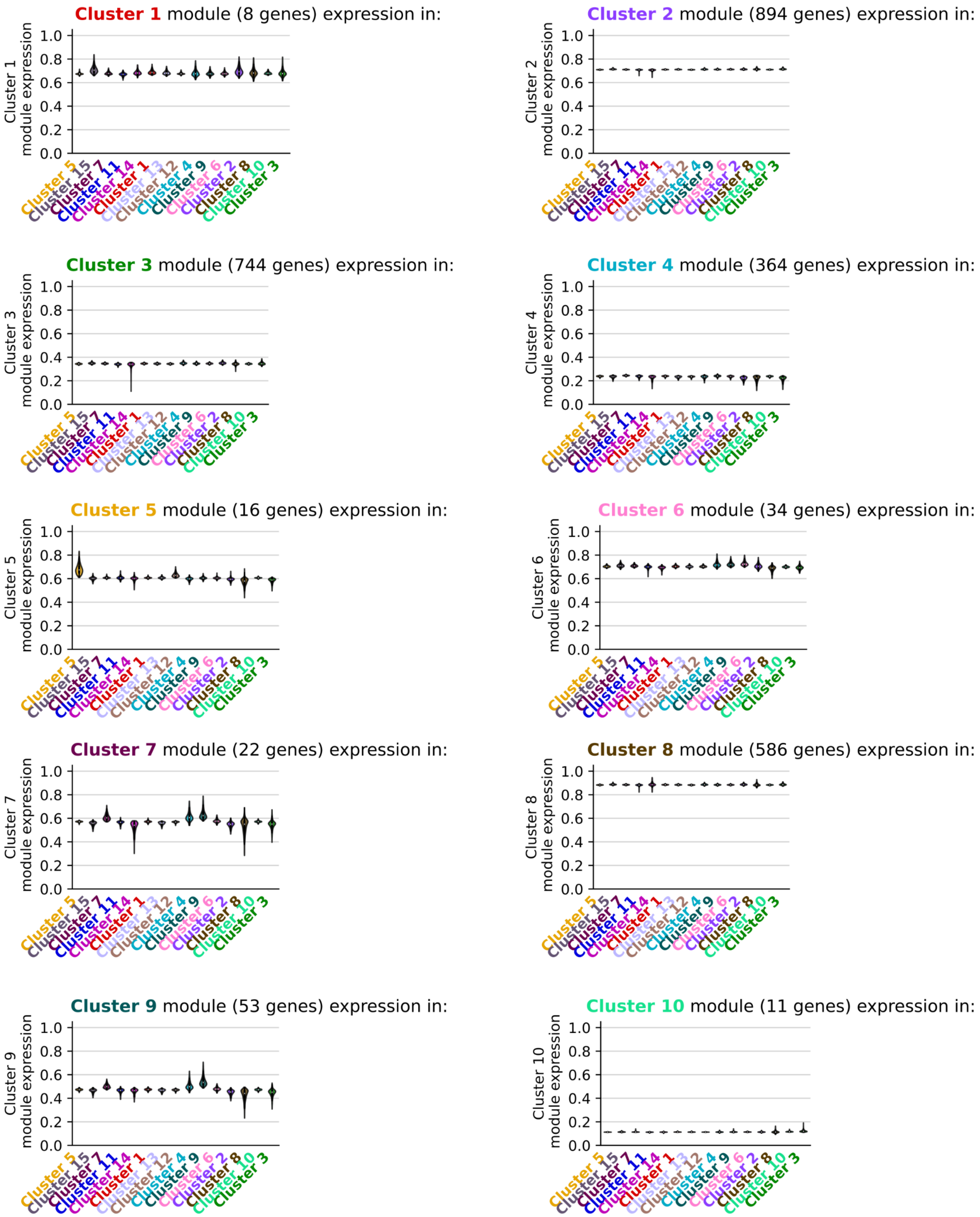

**Fig S14. Violin plots comparing the normalized expression of cluster-specific marker genes across clusters inferred by NCLUSION for the AML dataset ($N$ = 43,690 cells).** Each panel represents an evaluation in a given inferred cluster. The objective of this analysis is to see that each gene module exhibits the highest expression within its respective cluster relative to others. Results in this figure are continued in Fig S15.

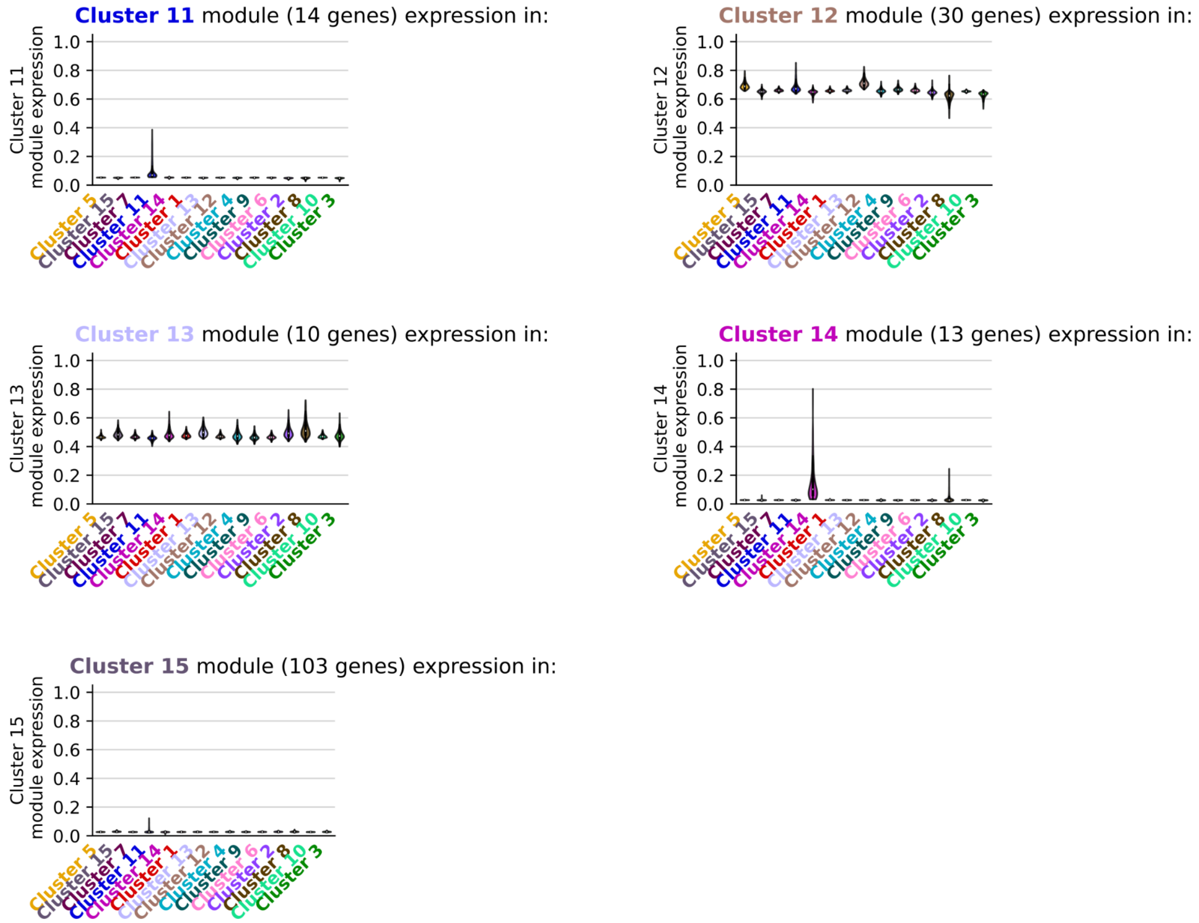

**Fig S15. Violin plots comparing the normalized expression of cluster-specific marker genes across clusters inferred by NCLUSION for the AML dataset ($N$ = 43,690 cells).** Each panel represents an evaluation in a given inferred cluster. The objective of this analysis is to see that each gene module exhibits the highest expression within its respective cluster relative to others. Results in this figure are an extension to violin plots in Fig S14.

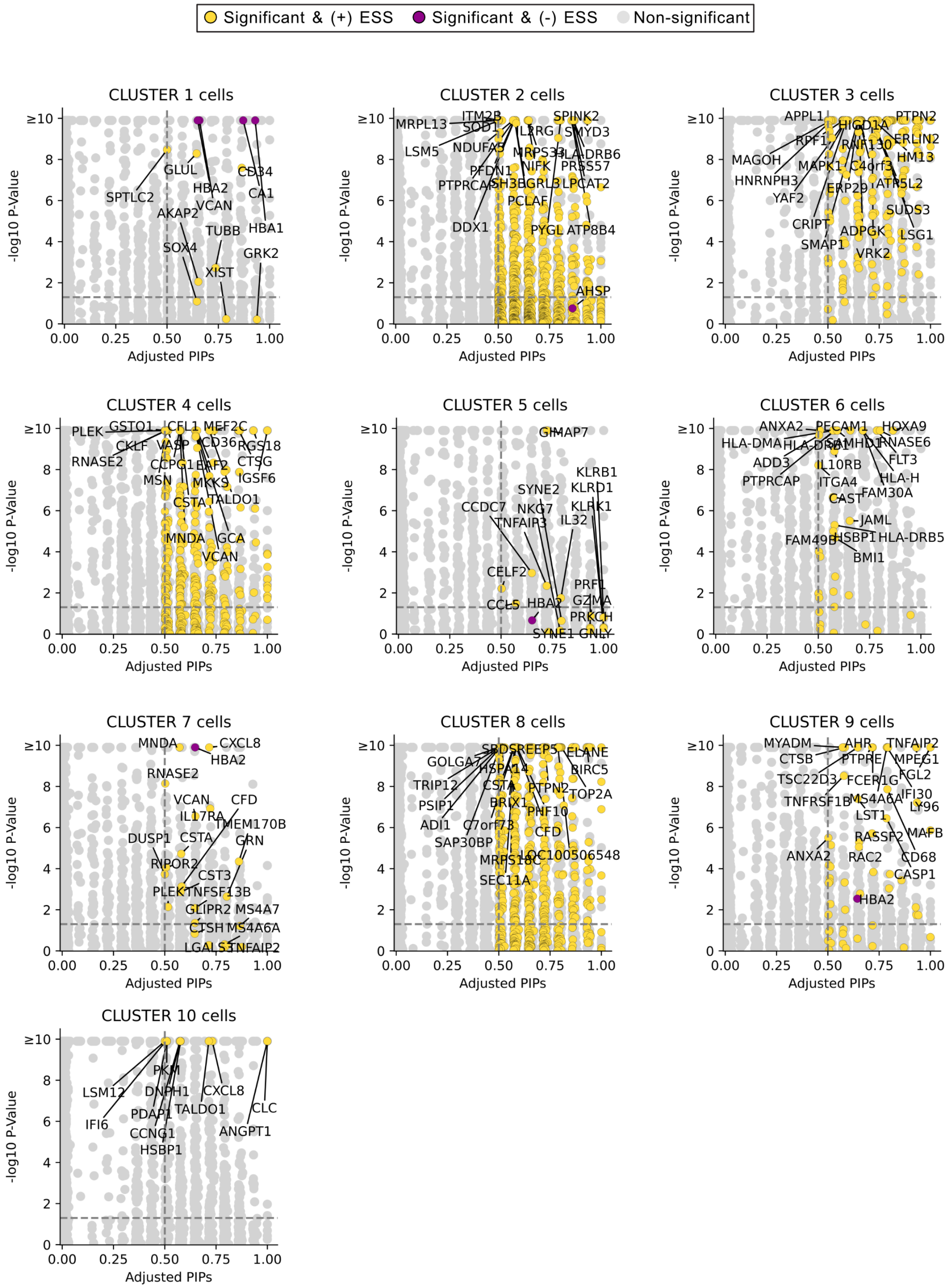

**Fig S16. Scatter plot comparing the marker genes identified using *post hoc* differential expression analysis with Seurat versus the variable selection approach with NCLUSION in the AML dataset ($N$ = 43,690 cells).** Here, our evaluation is based on NCLUSION's inferred cluster labels (depicted in each panel). The y-axis shows log-transformed $q$-values (i.e., adjusted $P$) for differentially expressed genes from Seurat found via a Wilcoxon rank sum test which is performed by doing a *post hoc* one-versus-all comparison for each cluster. The x-axis shows the adjusted posterior inclusion probabilities (PIPs) for each gene as computed by NCLUSION. All points in color are genes with PIP $\geq 0.5$. The specific color corresponds to the effect size sign (ESS) for each gene. Yellow points are genes with ESS = + (up-regulated), while purple points are genes with ESS = - (down-regulated). The vertical dashed line marks the median probability criterion[12]. The horizontal dashed line marks the Bonferroni-corrected threshold corresponding to significant $q$-values $\leq 0.05$. Genes found above the horizontal line and to the right of the vertical line are selected by both approaches; while, elements in the bottom right and top left quadrants are uniquely identified by NCLUSION and Seurat, respectively. Results in this figure are continued in Fig S17.

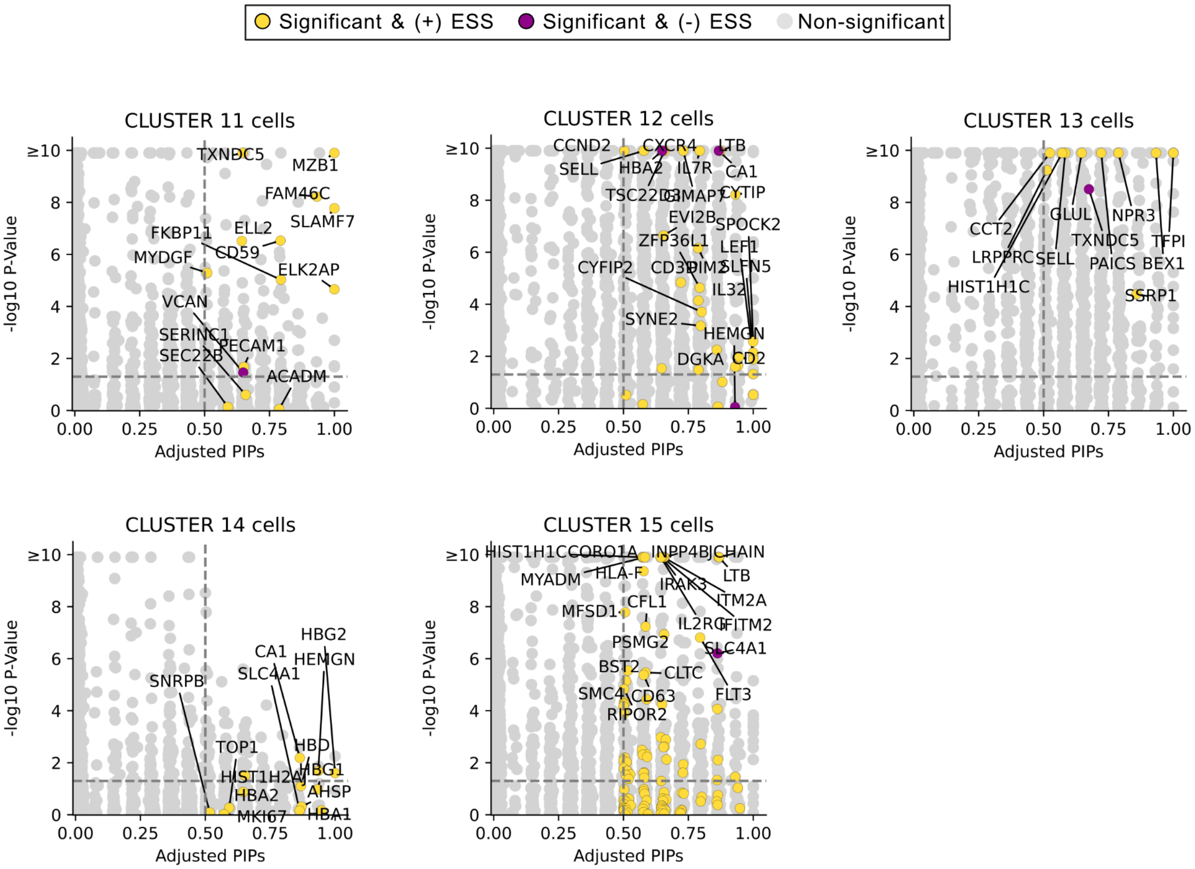

**Fig S17. Scatter plot comparing the marker genes identified using *post hoc* differential expression analysis with Seurat versus the variable selection approach with NCLUSION in the AML dataset ($N$ = 43,690 cells).** Here, our evaluation is based on NCLUSION's inferred cluster labels (depicted in each panel). The y-axis shows log-transformed $q$-values (i.e., adjusted $P$) for differentially expressed genes from Seurat found via a Wilcoxon rank sum test which is performed by doing a *post hoc* one-versus-all comparison for each cluster. The x-axis shows the adjusted posterior inclusion probabilities (PIPs) for each gene as computed by NCLUSION. All points in color are genes with PIP $\geq 0.5$. The specific color corresponds to the effect size sign (ESS) for each gene. Yellow points are genes with ESS = + (up-regulated), while purple points are genes with ESS = - (down-regulated). The vertical dashed line marks the median probability criterion[12]. The horizontal dashed line marks the Bonferroni-corrected threshold corresponding to significant $q$-values $\leq 0.05$. Genes found above the horizontal line and to the right of the vertical line are selected by both approaches; while, elements in the bottom right and top left quadrants are uniquely identified by NCLUSION and Seurat, respectively. Results in this figure are an extension of the scatter plots shown in Fig S16.

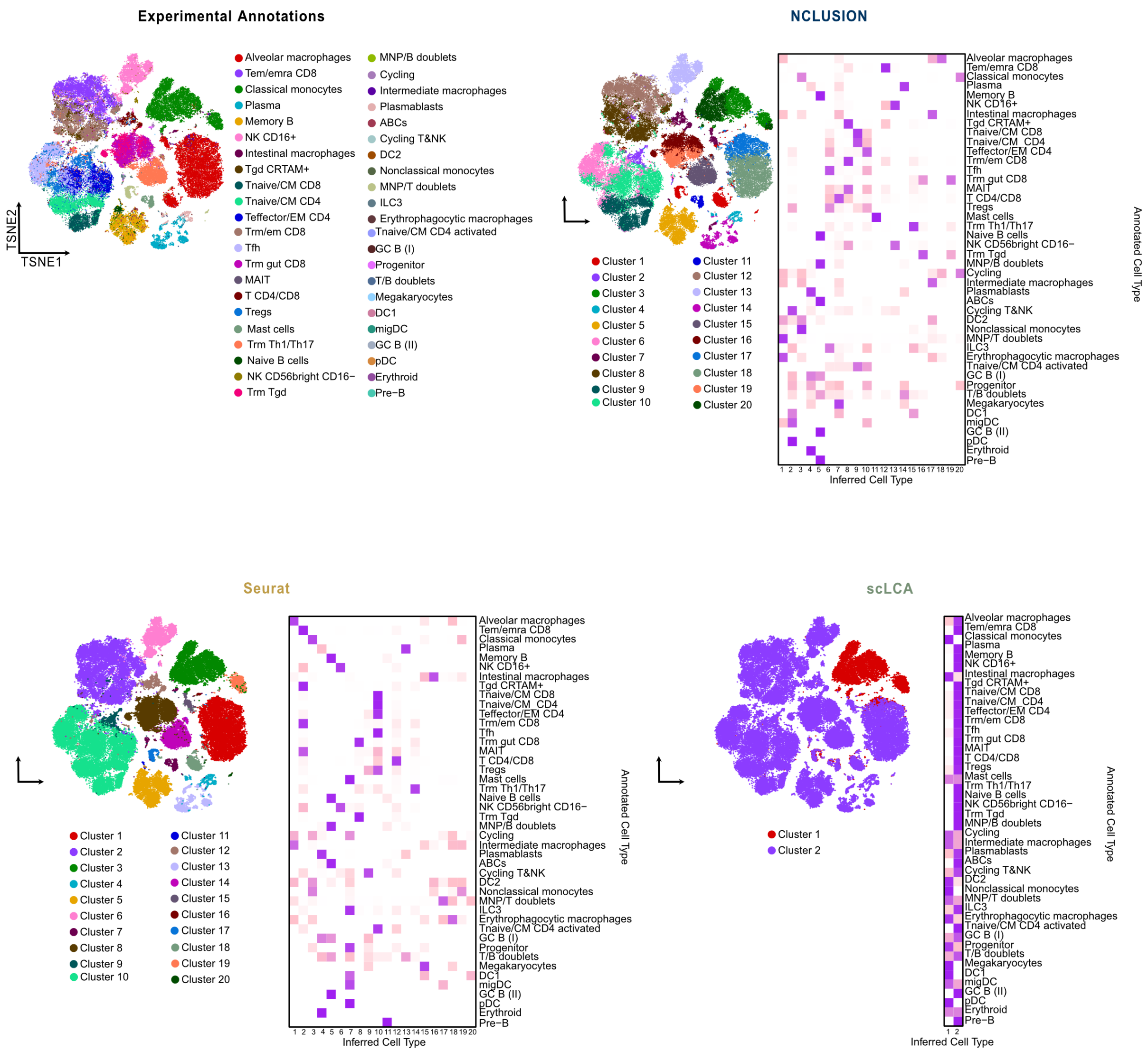

**Fig S18. Comparing the quality of clustering for NCLUSION and other baseline methods on the IMMUNE dataset ($N$ = 88,057 cells).** The first panel in the top left corner shows an overview of the cellular annotations found in the original study by Domínguez Conde et al.[51]. The other panels depict results from running NCLUSION and other baseline methods on the same data. These baselines include: Seurat, scLCA, K-nearest neighbors followed by the Leiden clustering algorithm (KNN+Leiden), SOUP, and scCCESS-SIMLR, respectively. Here, we visualize the structure of the inferred clusters across all baselines using t-distributed stochastic neighbor embeddings (t-SNEs) on the left hand side of each panel and a contingency heat map showing the prevalence of each cell type across clusters on the right hand side. The results in this image are continued in Fig S19.

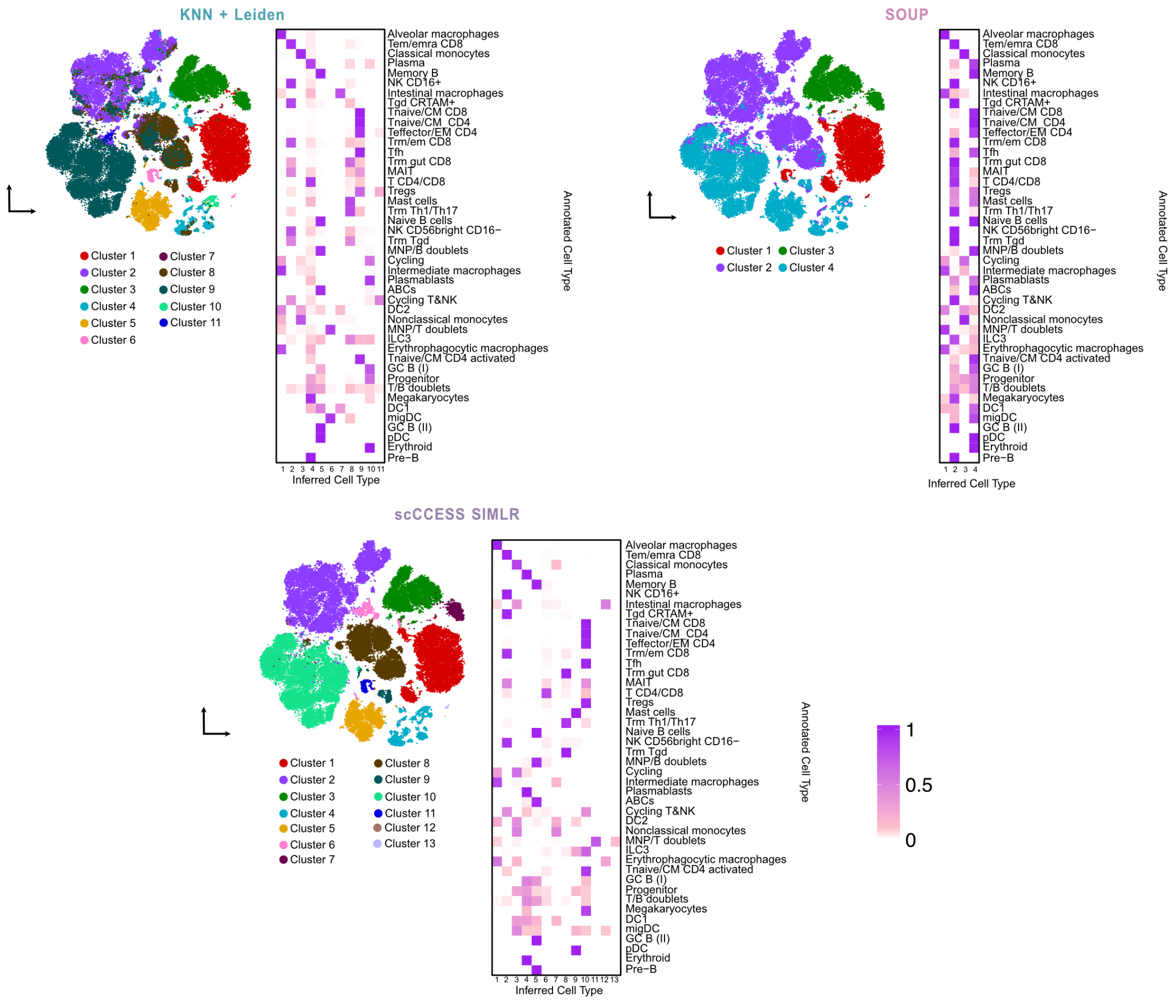

**Fig S19. Comparing the quality of clustering for NCLUSION and other baseline methods on the IMMUNE dataset ($N$ = 88,057 cells).** The first panel in the top left corner shows an overview of the cellular annotations found in the original study by Domínguez Conde et al. [51]. The other panels depict results from running NCLUSION and other baseline methods on the same data. These baselines include: Seurat, scLCA, K-nearest neighbors followed by the Leiden clustering algorithm (KNN+Leiden), SOUP, and scCCESS-SIMLR, respectively. Here, we visualize the structure of the inferred clusters across all baselines using t-distributed stochastic neighbor embeddings (t-SNEs) on the left hand side of each panel and a contingency heat map showing the prevalence of each cell type across clusters on the right hand side. The results in this image are a continuation of Fig S18.

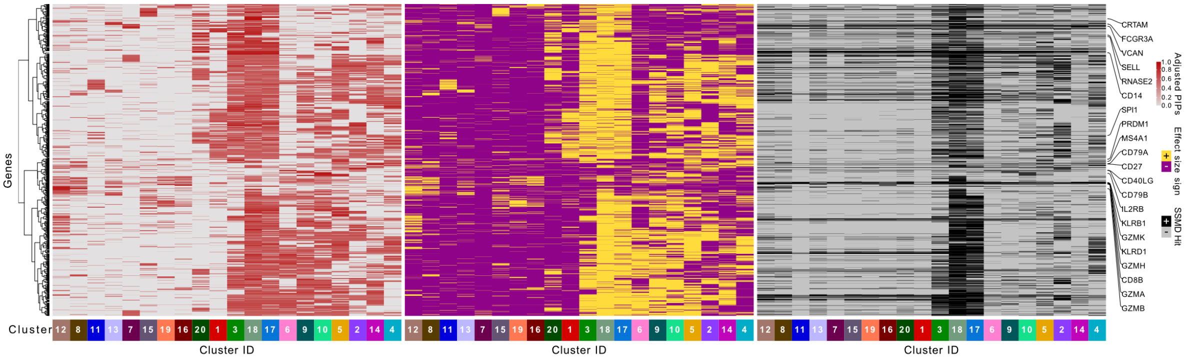

**Fig S20. Heat map of the significant genes as determined by (left) adjusted posterior inclusion probabilities (PIPs), (center) effect size sign (ESS), and (right) strictly standardized mean difference (SSMD) in each inferred cluster from NCLUSION for the IMMUNE dataset ($N$ = 88,057 cells). (Left)** Genes that are identified as "important" across many different clusters are seen as ubiquitous housekeeping variables rather than significant marker genes of unique cell types. Therefore, we down-weight the inclusion probabilities to proportionally penalize genes based on the number of clusters in which they appear. Here, significant genes are selected as those with adjusted PIPs greater than or equal to 0.5 which corresponds to the median probability criterion[12]. **(Center)** The ESS for each gene is computed by taking the sign of Cohen's $d$[13] between the expression of the $j$-th gene for cells in the $k$-th cluster and cells not in the $k$-th cluster. A positive ESS can be interpreted as the $j$-th gene being up-regulated in a given cluster. **(Right)** Significant genes are selected as those with SSMD greater than or equal to 0.15 which corresponds to the excluding genes with effect sizes that are weak[17]. In general, we observed that our criteria of cluster-specific marker genes was positively correlated with the expression of each gene.

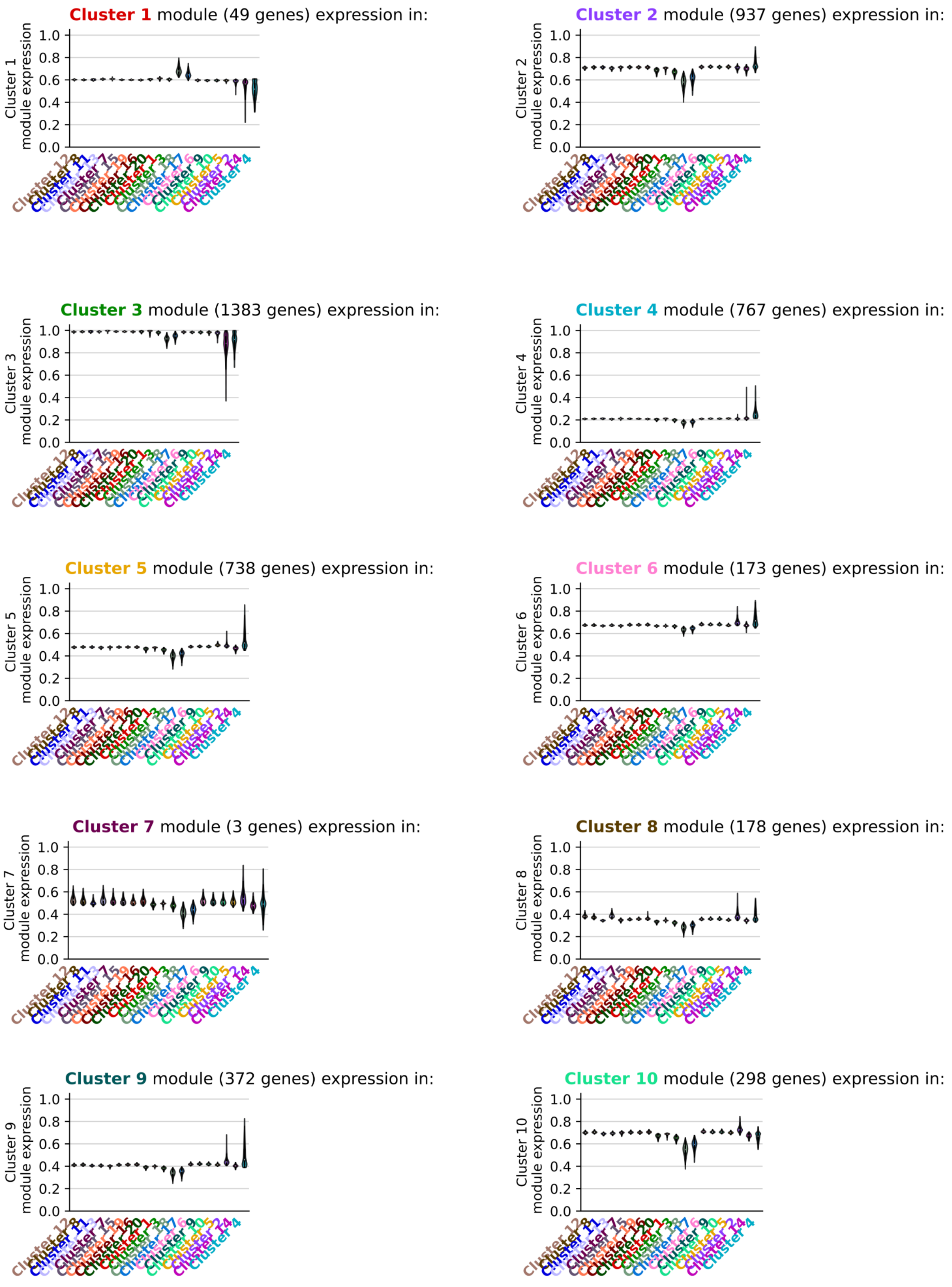

**Fig S21. Violin plots comparing the normalized expression of cluster-specific marker genes across clusters inferred by NCLUSION for the IMMUNE dataset ($N$ = 88,057 cells).** Each panel represents an evaluation in a given inferred cluster. The objective of this analysis is to see that each gene module exhibits the highest expression within its respective cluster relative to others. Results in this figure are continued in Fig S22.

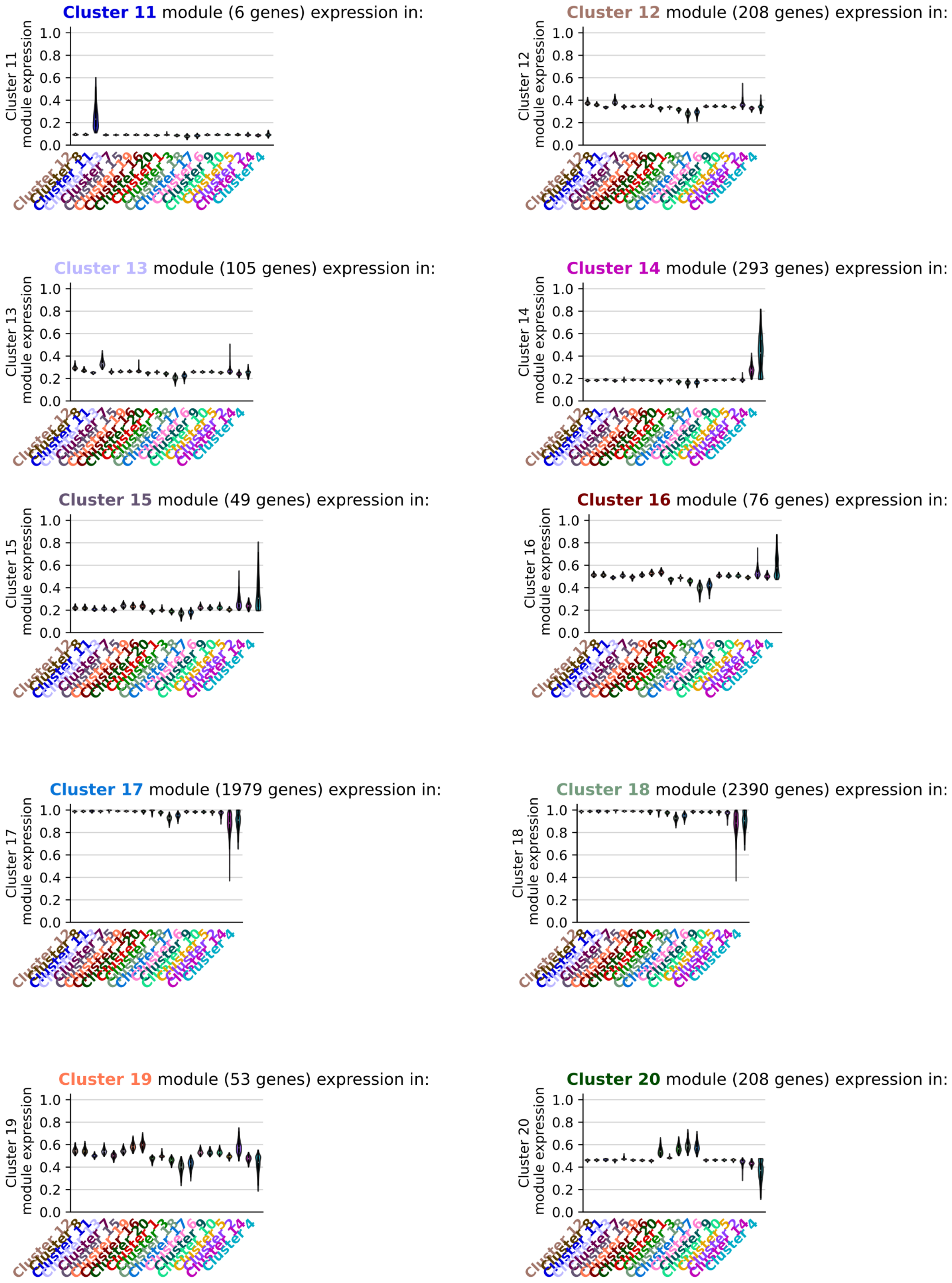

**Fig S22. Violin plots comparing the normalized expression of cluster-specific marker genes across clusters inferred by NCLUSION for the IMMUNE dataset ($N$ = 88,057 cells).** Each panel represents an evaluation in a given inferred cluster. The objective of this analysis is to see that each gene module exhibits the highest expression within its respective cluster relative to others. Results in this figure are an extension of the violin plots shown in Fig S21.

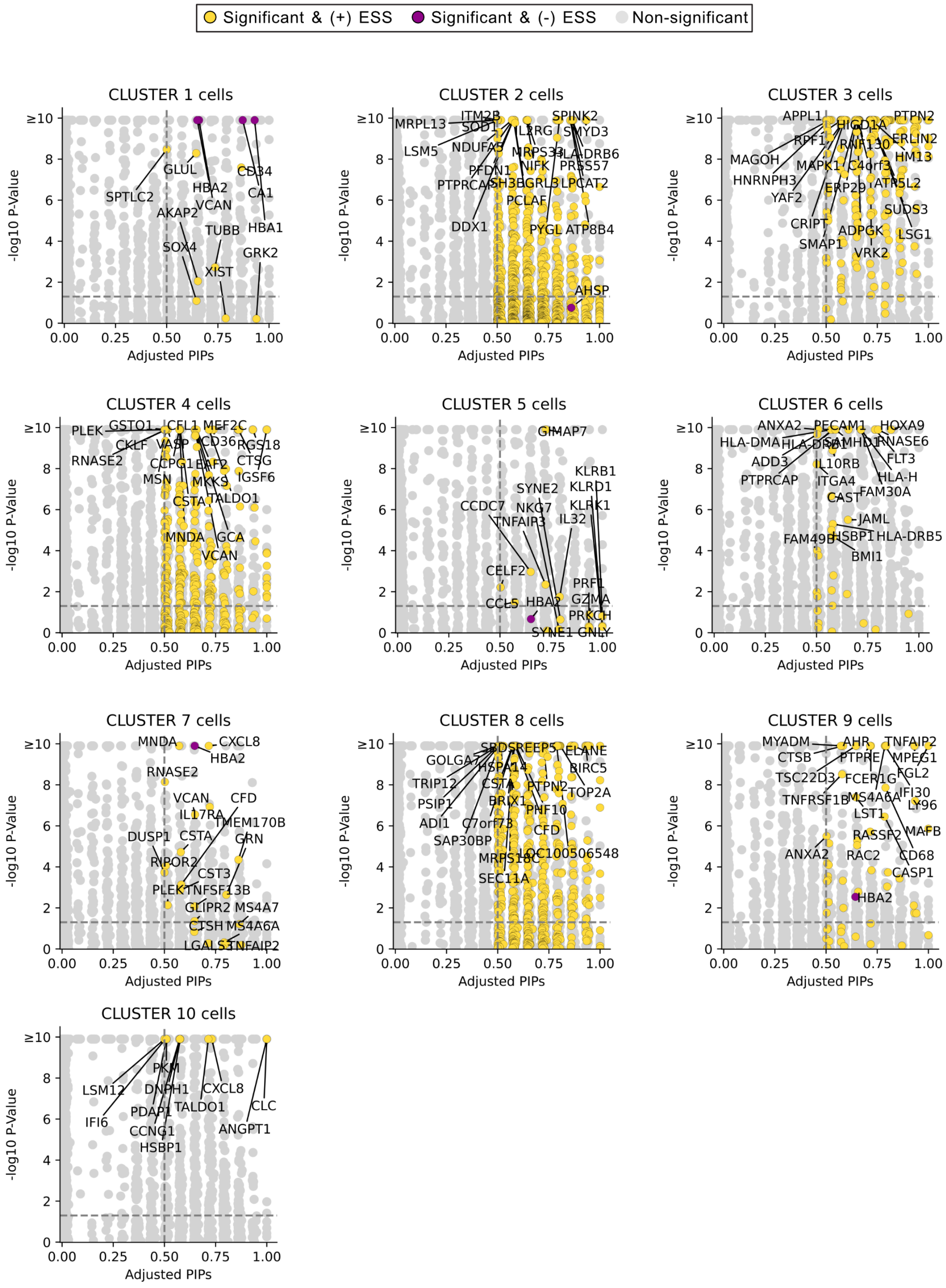

**Fig S23. Scatter plot comparing the marker genes identified using *post hoc* differential expression analysis with Seurat versus the variable selection approach with NCLUSION in the IMMUNE dataset ($N$ = 88,057 cells).** Here, our evaluation is based on NCLUSION's inferred cluster labels (depicted in each panel). The y-axis shows log-transformed $q$-values (i.e., adjusted $P$) for differentially expressed genes from Seurat found via a Wilcoxon rank sum test which is performed by doing a *post hoc* one-versus-all comparison for each cluster. The x-axis shows the adjusted posterior inclusion probabilities (PIPs) for each gene as computed by NCLUSION. All points in color are genes with PIP $\geq 0.5$. The specific color corresponds to the effect size sign (ESS) for each gene. Yellow points are genes with ESS = + (up-regulated), while purple points are genes with ESS = - (down-regulated). The vertical dashed line marks the median probability criterion[12]. The horizontal dashed line marks the Bonferroni-corrected threshold corresponding to significant $q$-values $\leq 0.05$. Genes found above the horizontal line and to the right of the vertical line are selected by both approaches; while, elements in the bottom right and top left quadrants are uniquely identified by NCLUSION and Seurat, respectively. Results in this figure continued in Fig S24.

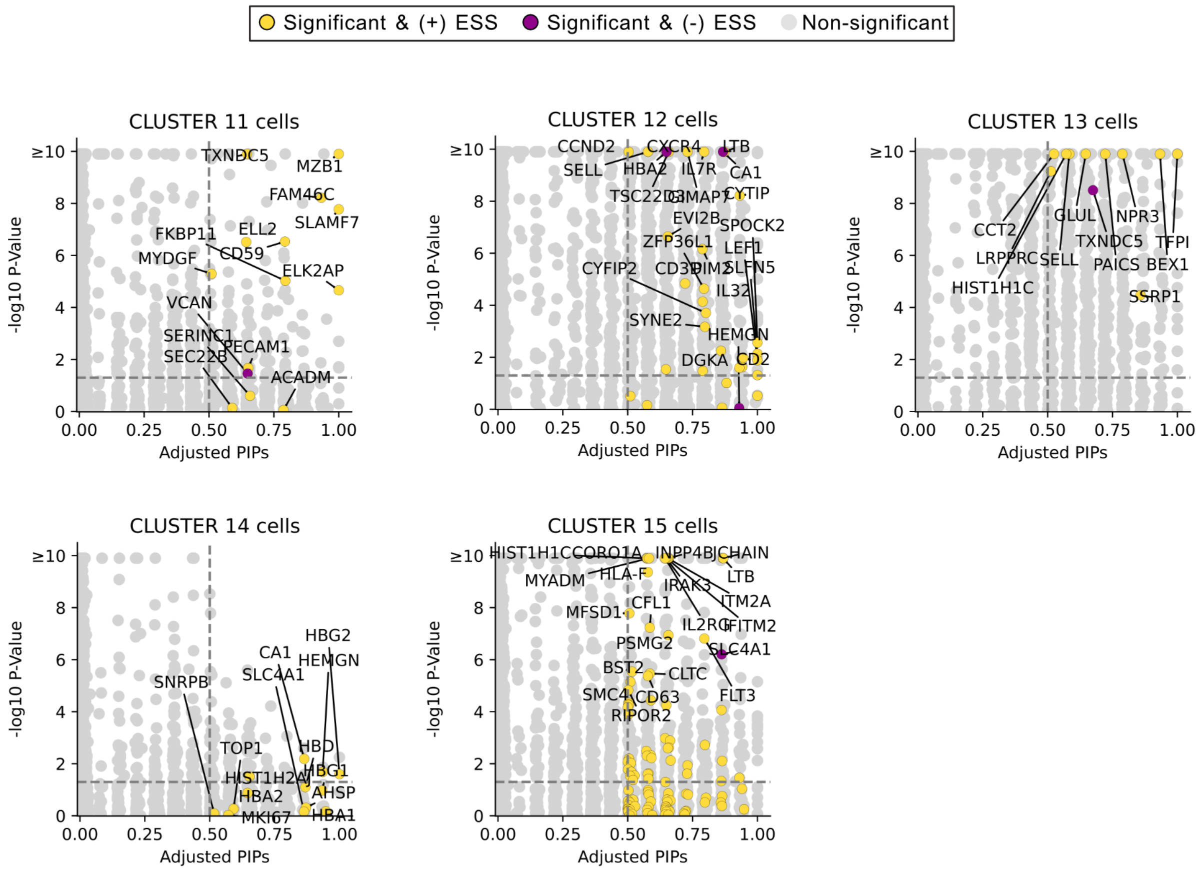

**Fig S24. Scatter plot comparing the marker genes identified using *post hoc* differential expression analysis with Seurat versus the variable selection approach with NCLUSION in the IMMUNE dataset ($N$ = 88,057 cells).** Here, our evaluation is based on NCLUSION's inferred cluster labels (depicted in each panel). The y-axis shows log-transformed $q$-values (i.e., adjusted $P$) for differentially expressed genes from Seurat found via a Wilcoxon rank sum test which is performed by doing a *post hoc* one-versus-all comparison for each cluster. The x-axis shows the adjusted posterior inclusion probabilities (PIPs) for each gene as computed by NCLUSION. All points in color are genes with PIP $\geq$ 0.5. The specific color corresponds to the effect size sign (ESS) for each gene. Yellow points are genes with ESS = + (up-regulated), while purple points are genes with ESS = - (down-regulated). The vertical dashed line marks the median probability criterion[12]. The horizontal dashed line marks the Bonferroni-corrected threshold corresponding to significant $q$-values $\leq$ 0.05. Genes found above the horizontal line and to the right of the vertical line are selected by both approaches; while, elements in the bottom right and top left quadrants are uniquely identified by NCLUSION and Seurat, respectively. Results in this figure are an extension of the scatter plots shown in Fig S23.

### 8 Supplementary Tables

**Table S1. Simulation clustering results.** This file gives the clustering evaluation metric values. We analyzed 20 independent simulated datasets based on the B cell, NK cell, Monocyte, and T Cell annotations provided by Zheng et al.[48]. We evaluated NCLUSION against the baseline methods: Seurat, K-nearest neighbors followed by the Leiden clustering algorithm (KNN+Leiden), scLCA, scCCESS-SIMLR, SOUP, SC3, scDeepCluster, and CIDR. For each method, we evaluated clustering performance according to the clustering metrics: normalized mutual information (NMI) and adjusted Rand index (ARI). We simulated 4 scenarios using `scDesign3`[47]. In Scenario I, we evenly distributed all synthetically generated cells across five clusters and each cluster had a unique set of 50 marker genes. In Scenario II, we implemented an imbalanced cluster design where one small cluster had 200 cells and the other four larger clusters each had 2450 cells. In Scenario III, there was also an imbalanced cluster design but a situation where one cluster had 20 (rare) cells and the other four larger clusters each had 2495 cells each. All clusters in Scenarios II and III had 50 unique marker genes each. Lastly, in Scenario IV, we generated balanced clusters of 2000 cells per cell type, but one cluster had only 20 marker genes while the other four clusters had 50 marker genes. The results in this table are related to Fig 2 and Fig S2. This supplementary table can also be accessed on Zenodo (`https://doi.org/10.5281/zenodo.11099807`). (CSV)

**Table S2. Simulated Marker genes recovery results.** This file gives the marker gene recovery evaluation metric values. We analyzed 20 independent simulated datasets based on the B cell, NK cell, Monocyte, and T Cell annotations provided by Zheng et al.[48]. We evaluated NCLUSION against the baseline methods: DUBStepR, FESTEM, and singleCellHaystack. For each method, we evaluated marker gene recovery performance according to the metrics: true positive rate (TPR; or power), false discovery rate (FDR), and false positive rate (FPR; computed as 1-Specificity). We simulated 4 scenarios using `scDesign3`[47]. In Scenario I, we evenly distributed all synthetically generated cells across five clusters and each cluster had a unique set of 50 marker genes. In Scenario II, we implemented an imbalanced cluster design where one small cluster had 200 cells and the other four larger clusters each had 2450 cells. In Scenario III, there was also an imbalanced cluster design but a situation where one cluster had 20 (rare) cells and the other four larger clusters each had 2495 cells each. All clusters in Scenarios II and III had 50 unique marker genes each. Lastly, in Scenario IV, we generated balanced clusters of 2000 cells per cell type, but one cluster had only 20 marker genes while the other four clusters had 50 marker genes. The results in this table are related to Fig 2 and Fig S3. This supplementary table can also be accessed on Zenodo (`https://doi.org/10.5281/zenodo.11099807`). (CSV)

**Table S3. Contingency tables showing the prevalence of each cell type within each inferred cluster from NCLUSION and other baseline methods in the PBMC dataset ($N$ = 94,615 cells).** This file gives the clustering contingency table for NCLUSION (page 1) and baseline methods including: SOUP (page 2), K-nearest neighbors followed by the Leiden clustering algorithm (KNN+Leiden) (page 3), scCCESS-SIMLR (page 4), scLCA (page 5), and Seurat (page 6). The columns of each table are the clusters inferred by each algorithm. The rows are the cellular annotations found in the original study by Zheng et al.[48]. The values in the table are calculated by taking the number of cells in each inferred cluster and dividing it by the total number of cells with a given experimental annotation label. Therefore, each row of the table sum to 1. The results in this table are related to Fig 3. This supplementary table can also be accessed on Zenodo (`https://doi.org/10.5281/zenodo.11099807`). (XLSX)

**Table S4. Evaluation metrics for each inferred cluster from NCLUSION and other baseline methods in all real datasets (PBMC, PDAC, IMMUNE, and AML).** This file gives the clustering evaluation metric values. On page 1, we analyzed five independent subsamples of each datasets where we subsampled 80% of the total number of cells to analyze. For each method, we evaluated clustering performance according to the clustering metrics: normalized mutual information (NMI) and adjusted Rand index (ARI). On page 2, we list the mean, standard deviation, and standard error, and two-sided t-test $P$ compared to NCLUSION's mean for each method according to each clustering metric. Lastly, on page 2, we give a table of the $P$-values from an ANOVA and Kruskal–Wallis $H$-test assessing the statistical difference between the median performance of NCLUSION compared to that of the competing methods in all datasets. The results in this table are related to Fig 3 and Fig 5. This supplementary table can also be accessed on Zenodo (`https://doi.org/10.5281/zenodo.11099807`). (XLSX)

**Table S5. List of adjusted posterior inclusion probabilities (PIPs), effect size sign (ESS), and the strictly standardized mean difference (SSMD) for each gene in each inferred cluster by NCLUSION in the PBMC dataset ($N$ = 94,615 cells).** On page 1, the first column represents the original NCLUSION inferred cluster label, and the second column represents the relabeled cluster label used in downstream analysis. On pages 2-4, the columns on each page correspond to the clusters inferred by NCLUSION and the rows correspond to the genes whose adjusted PIPs were significant in at least one cluster according to the "median probability model" threshold[12] (i.e., PIP $\geq$ 0.5). Page 2 provides the adjusted PIPs for each gene across the inferred clusters, where the adjustment weight is calculated to penalize genes according to the number of clusters in which they appear to be important (Methods). Page 3 provides the ESS for significant genes which is computed by taking the sign of Cohen's $d^{13}$ between the expression of the $j$-th gene for cells in the $k$-th cluster and cells not in the $k$-th cluster. Page 4 then provides the SSMD for the same set of significant genes. Pages 5-14 list the genes with significant adjusted PIPs, positive ESS, and significant SSMD that make up the cluster's gene module. Pages 15-24 display results from a *post hoc* differential expression analysis using Seurat with the cell annotations from NCLUSION's cluster label based on a Wilcoxon rank sum test. The results in this table are related to Figs 4, S5, and S6. This supplementary table can also be accessed on Zenodo (`https://doi.org/10.5281/zenodo.11099807`). (XLSX)

**Table S6. List of NCLUSION-derived gene module expression in each cell in the PBMC dataset ($N$ = 94,615 cells) and the overlap with Seurat-based module generation.** Page 1 lists the expression of each NCLUSION-derived gene module expression in each cell. Page 2 lists the percent overlap of genes between the NCLUSION-derived gene module and the Seurat-derived differentially expressed gene list for each cluster. The results in this table are related to Fig S4. This supplementary table can also be accessed on Zenodo (`https://doi.org/10.5281/zenodo.11099807`). (XLSX)

**Table S7. List of Seurat-based gene module generation and GO results for each inferred cluster by NCLUSION and Seurat in the PBMC dataset ($N$ = 94,615 cells).** Pages 1-10 display results from a *post hoc* differential expression analysis using Seurat with the cell annotations from the original study based on a Wilcoxon rank sum test. For marker gene selection approaches based on a Wilcoxon rank sum test, we performed a one-versus-all comparison with the clusters. We also included are the adjusted PIPs generated by NCLUSION for these genes in the same corresponding cluster for comparison. Pages 11-20 list the results from the Gene Ontology (GO) analysis using the cluster-specific marker genes identified by NCLUSION. Pages 21-30 list the results from the Gene Ontology (GO) analysis using the Seuarat's *post hoc* differential expression analysis. The results in this table are related to Fig 4. This supplementary table can also be accessed on Zenodo (`https://doi.org/10.5281/zenodo.11099807`). (XLSX)

**Table S8. Contingency tables showing the prevalence of each cell type within each inferred cluster from NCLUSION and other baseline methods in the PDAC dataset ($N$ = 23, 042 cells).** This file gives the clustering contingency table for NCLUSION (page 1) and baseline methods including: SOUP (page 2), K-nearest neighbors followed by the Leiden clustering algorithm (KNN+Leiden) (page 3), scCCESS-SIMLR (page 4), scLCA (page 5), and Seurat (page 6). The columns of each table are the clusters inferred by each algorithm. The rows are the cellular annotations found in the original study by Raghavan et al.[49]. The values in the table are calculated by taking the number of cells in each inferred cluster and dividing it by the total number of cells with a given experimental annotation label. Therefore, each row of the table sum to 1. The results in this table are related to Fig 5. This supplementary table can also be accessed on Zenodo (`https://doi.org/10.5281/zenodo.11099807`). (XLSX)

**Table S9. List of adjusted posterior inclusion probabilities (PIPs), effect size sign (ESS), and the strictly standardized mean difference (SSMD) for each gene in each inferred cluster by NCLUSION in the PDAC dataset ($N$ = 23,042 cells).** On page 1, the first column represents the original NCLUSION inferred cluster label, and the second column represents the relabeled cluster label used in downstream analysis. On pages 2-4, the columns on each page correspond to the clusters inferred by NCLUSION and the rows correspond to the genes whose adjusted PIPs were significant in at least one cluster according to the "median probability model" threshold[12] (i.e., PIP $\geq 0.5$). Page 2 provides the adjusted PIPs for each gene across the inferred clusters, where the adjustment weight is calculated to penalize genes according to the number of clusters in which they appear to be important (Methods). Page 3 provides the ESS for significant genes which is computed by taking the sign of Cohen's $d$[13] between the expression of the $j$-th gene for cells in the $k$-th cluster and cells not in the $k$-th cluster. Page 4 then provides the SSMD for the same set of significant genes. Pages 5-18 list the genes with significant adjusted PIPs, positive ESS, and significant SSMD that make up the cluster's gene module. Pages 19-32 display results from a *post hoc* differential expression analysis using Seurat with the cell annotations from NCLUSION's cluster label based on a Wilcoxon rank sum test. The results in this table are related to Figs 5, S10 and S11. This supplementary table can also be accessed on Zenodo (`https://doi.org/10.5281/zenodo.11099807`). (XLSX)

**Table S10. List of NCLUSION-derived gene module expression in each cell in the PDAC dataset ($N$ = 23,042 cells) and the overlap with Seurat-based module generation.** Page 1 lists the expression of each NCLUSION-derived gene module expression in each cell. Page 2 lists the percent overlap of genes between the NCLUSION-derived gene module and the Seurat-derived differentially expressed gene list for each cluster. The results in this table are related to Figs S8 and S9. This supplementary table can also be accessed on Zenodo (`https://doi.org/10.5281/zenodo.11099807`). (XLSX)

**Table S11. List of Seurat-based gene module generation and GO results for each inferred cluster by NCLUSION and Seurat in the PDAC dataset ($N$ = 23,042 cells).** Pages 1-31 display results from a *post hoc* differential expression analysis using Seurat with the cell annotations from the original study based on a Wilcoxon rank sum test. For marker gene selection approaches based on a Wilcoxon rank sum test, we performed a one-versus-all comparison with the clusters. We also included are the adjusted PIPs generated by NCLUSION for these genes in the same corresponding cluster for comparison. Pages 32-44 list the results from the Gene Ontology (GO) analysis using the cluster-specific marker genes identified by NCLUSION. Pages 45-57 list the results from the Gene Ontology (GO) analysis using the Seuarat's *post hoc* differential expression analysis. This supplementary table can also be accessed on Zenodo (`https://doi.org/10.5281/zenodo.11099807`). (XLSX)

**Table S12. Contingency tables showing the prevalence of each cell type within each inferred cluster from NCLUSION and other baseline methods in the IMMUNE dataset ($N$ = 88,057 cells).** This file gives the clustering contingency table for NCLUSION (page 1) and baseline methods including: SOUP (page 2), K-nearest neighbors followed by the Leiden clustering algorithm (KNN+Leiden) (page 3), scCCESS-SIMLR (page 4), scLCA (page 5), and Seurat (page 6). The columns of each table are the clusters inferred by each algorithm. The rows are the cellular annotations found in the original study by Domínguez Conde et al.[51]. The values in the table are calculated by taking the number of cells in each inferred cluster and dividing it by the total number of cells with a given experimental annotation label. Therefore, each row of the table sum to 1. On page 7, we list the mean, standard deviation, and standard error for each method according to four clustering metrics: Jaccard similarity index, Gini-Simpson index, McIntosh evenness measure, and adjusted Rand index. Here, we analyzed five independent datasets where we subsampled 80% of the total number of cells to analyze. Lastly, on page 8, we give a table of the $P$-values from a Kruskal–Wallis $H$-test assessing the statistical difference between the median performance of NCLUSION compared to that of the competing methods. The results in this table are related to Figs S18 and S19. This supplementary table can also be accessed on Zenodo (`https://doi.org/10.5281/zenodo.11099807`). (XLSX)

**Table S13. List of adjusted posterior inclusion probabilities (PIPs), effect size sign (ESS), and the strictly standardized mean difference (SSMD) for each gene in each inferred cluster by NCLUSION in the IMMUNE dataset ($N$ = 88,057 cells).** On page 1, the first column represents the original NCLUSION inferred cluster label, and the second column represents the relabeled cluster label used in downstream analysis. On pages 2-4, the columns on each page correspond to the clusters inferred by NCLUSION and the rows correspond to the genes whose adjusted PIPs were significant in at least one cluster according to the "median probability model" threshold[12] (i.e., PIP $\geq$ 0.5). Page 2 provides the adjusted PIPs for each gene across the inferred clusters, where the adjustment weight is calculated to penalize genes according to the number of clusters in which they appear to be important (Methods). Page 3 provides the ESS for significant genes which is computed by taking the sign of Cohen's $d$[13] between the expression of the $j$-th gene for cells in the $k$-th cluster and cells not in the $k$-th cluster. Page 4 then provides the SSMD for the same set of significant genes. Pages 5-24 list the genes with significant adjusted PIPs, positive ESS, and significant SSMD that make up the cluster's gene module. Pages 25-44 display results from a *post hoc* differential expression analysis using Seurat with the cell annotations from NCLUSION's cluster label based on a Wilcoxon rank sum test. The results in this table are related to Figs S20, S23 and S24. This supplementary table can also be accessed on Zenodo (`https://doi.org/10.5281/zenodo.11099807`). (XLSX)

**Table S14. List of NCLUSION-derived gene module expression in each cell in the IMMUNE dataset ($N$ = 88,057 cells) and the overlap with Seurat-based module generation.** Page 1 lists the expression of each NCLUSION-derived gene module expression in each cell. Page 2 lists the percent overlap of genes between the NCLUSION-derived gene module and the Seurat-derived differentially expressed gene list for each cluster. The results in this table are related to Figs S21 and S22. This supplementary table can also be accessed on Zenodo (`https://doi.org/10.5281/zenodo.11099807`). (XLSX)

**Table S15. List of Seurat-based gene module generation and GO results for each inferred cluster by NCLUSION and Seurat in the IMMUNE dataset ($N$ = 88,057 cells).** Pages 1-39 display results from a *post hoc* differential expression analysis using Seurat with the cell annotations from the original study based on a Wilcoxon rank sum test. For marker gene selection approaches based on a Wilcoxon rank sum test, we performed a one-versus-all comparison with the clusters. We also included are the adjusted PIPs generated by NCLUSION for these genes in the same corresponding cluster for comparison. Pages 40-58 list the results from the Gene Ontology (GO) analysis using the cluster-specific marker genes identified by NCLUSION. Pages 59-77 list the results from the Gene Ontology (GO) analysis using the Seuarat's *post hoc* differential expression analysis. This supplementary table can also be accessed on Zenodo (`https://doi.org/10.5281/zenodo.11099807`). (XLSX)

**Table S16. Contingency tables showing the prevalence of each cell type within each inferred cluster from NCLUSION and other baseline methods in the AML dataset ($N$ = 43,690 cells).** This file gives the clustering contingency table for NCLUSION (page 1) and baseline methods including: SOUP (page 2), K-nearest neighbors followed by the Leiden clustering algorithm (KNN+Leiden) (page 3), scCCESS-SIMLR (page 4), scLCA (page 5), and Seurat (page 6). The columns of each table are the clusters inferred by each algorithm. The rows are the cellular annotations found in the original study by Domínguez Conde et al. [51]. The values in the table are calculated by taking the number of cells in each inferred cluster and dividing it by the total number of cells with a given experimental annotation label. Therefore, each row of the table sum to 1. On page 7, we list the mean, standard deviation, and standard error for each method according to four clustering metrics: Jaccard similarity index, Gini-Simpson index, McIntosh evenness measure, and adjusted Rand index. Here, we analyzed five independent datasets where we subsampled 80% of the total number of cells to analyze. Lastly, on page 8, we give a table of the $P$-values from a Kruskal–Wallis $H$-test assessing the statistical difference between the median performance of NCLUSION compared to that of the competing methods. The results in this table are related to Fig S12. This supplementary table can also be accessed on Zenodo (`https://doi.org/10.5281/zenodo.11099807`). (XLSX)

**Table S17. List of adjusted posterior inclusion probabilities (PIPs), effect size sign (ESS), and the strictly standardized mean difference (SSMD) for each gene in each inferred cluster by NCLUSION in the AML dataset ($N$ = 43,690 cells).** On page 1, the first column represents the original NCLUSION inferred cluster label, and the second column represents the relabeled cluster label used in downstream analysis. On pages 2-4, the columns on each page correspond to the clusters inferred by NCLUSION and the rows correspond to the genes whose adjusted PIPs were significant in at least one cluster according to the "median probability model" threshold [12] (i.e., PIP $\geq$ 0.5). Page 2 provides the adjusted PIPs for each gene across the inferred clusters, where the adjustment weight is calculated to penalize genes according to the number of clusters in which they appear to be important (Methods). Page 3 provides the ESS for significant genes which is computed by taking the sign of Cohen's $d$[13] between the expression of the $j$-th gene for cells in the $k$-th cluster and cells not in the $k$-th cluster. Page 4 then provides the SSMD for the same set of significant genes. Pages 5-19 list the genes with significant adjusted PIPs, positive ESS, and significant SSMD that make up the cluster's gene module. Pages 20-34 display results from a *post hoc* differential expression analysis using Seurat with the cell annotations from NCLUSION's cluster label based on a Wilcoxon rank sum test. The results in this table are related to Figs S13, S16, and S17. This supplementary table can also be accessed on Zenodo (`https://doi.org/10.5281/zenodo.11099807`). (XLSX)

**Table S18. List of NCLUSION-derived gene module expression in each cell in the AML dataset ($N$ = 88,057 cells) and the overlap with Seurat-based module generation.** Page 1 lists the expression of each NCLUSION-derived gene module expression in each cell. Page 2 lists the percent overlap of genes between the NCLUSION-derived gene module and the Seurat-derived differentially expressed gene list for each cluster. The results in this table are related to Figs S14 and S15. This supplementary table can also be accessed on Zenodo (`https://doi.org/10.5281/zenodo.11099807`). (XLSX)

**Table S19. List of Seurat-based gene module generation and GO results for each inferred cluster by NCLUSION and Seurat in the AML dataset ($N$ = 43,690 cells).** Pages 1-31 display results from a *post hoc* differential expression analysis using Seurat with the cell annotations from the original study based on a Wilcoxon rank sum test. For marker gene selection approaches based on a Wilcoxon rank sum test, we performed a one-versus-all comparison with the clusters. We also included are the adjusted PIPs generated by NCLUSION for these genes in the same corresponding cluster for comparison. Pages 32-45 list the results from the Gene Ontology (GO) analysis using the cluster-specific marker genes identified by NCLUSION. Pages 46-59 list the results from the Gene Ontology (GO) analysis using the Seuarat's *post hoc* differential expression analysis. This supplementary table can also be accessed on Zenodo (`https://doi.org/10.5281/zenodo.11099807`). (XLSX)

13. Jacob Cohen. Statistical Power Analysis for the Behavioral Sciences. Academic press, 2013.

19. Xiaohua Douglas Zhang, Raul Lacson, Ruojing Yang, Shane D. Marine, Alex McCampbell, Dawn M. Toolan, Tim R. Hare, Joleen Kajdas, Joel P. Berger, Daniel J. Holder, Joseph F. Heyse, and Marc Ferrer. The use of ssmd-based false discovery and false nondiscovery rates in

genome-scale rnai screens. SLAS Discovery, 15(9):1123–1131, October 2010. ISSN 2472-5552. doi: 10.1177/1087057110381919.

20. Michael I Jordan, Zoubin Ghahramani, Tommi S Jaakkola, and Lawrence K Saul. An introduction to variational methods for graphical models. Machine Learning, 37(2):183–233, 1999.

21. John T Ormerod and Matt P Wand. Explaining variational approximations. The American Statistician, 64(2):140–153, 2010.

22. Christopher M Bishop. Pattern Recognition and Machine Learning. Springer, 2006. ISBN 1493938436.

28. Percy Liang, Slav Petrov, Michael Jordan, and Dan Klein. The infinite pcfg using hierarchical dirichlet processes. In Jason Eisner, editor, Proceedings of the 2007 Joint Conference on Empirical Methods in Natural Language Processing and Computational Natural Language

Learning (EMNLP-CoNLL), page 688–697, Prague, Czech Republic, June 2007. Association for Computational Linguistics. URL `https://aclanthology.org/D07-1072/`.

36. F. Pedregosa, G. Varoquaux, A. Gramfort, V. Michel, B. Thirion, O. Grisel, M. Blondel, P. Prettenhofer, R. Weiss, V. Dubourg, J. Vanderplas, A. Passos, D. Cournapeau, M. Brucher, M. Perrot, and

E. Duchesnay. Scikit-learn: Machine learning in Python. Journal of Machine Learning Research, 12:2825–2830, 2011.

49. Srivatsan Raghavan, Peter S. Winter, Andrew W. Navia, Hannah L. Williams, Alan DenAdel, Kristen E. Lowder, Jennyfer Galvez-Reyes, Radha L. Kalekar, Nolawit Mulugeta, Kevin S. Kapner, Manisha S. Raghavan, Ashir A. Borah, Nuo Liu, Sara A. Väyrynen, Andressa Dias Costa, Raymond W. S. Ng, Junning Wang, Emma K. Hill, Dorisanne Y. Ragon, Lauren K. Brais, Alex M. Jaeger,

Liam F. Spurr, Yvonne Y. Li, Andrew D. Cherniack, Matthew A. Booker, Elizabeth F. Cohen, Michael Y. Tolstorukov, Isaac Wakiro, Asaf Rotem, Bruce E. Johnson, James M. McFarland, Ewa T. Sicinska, Tyler E. Jacks, Ryan J. Sullivan, Geoffrey I. Shapiro, Thomas E. Clancy, Kimberly Perez, Douglas A. Rubinson, Kimmie Ng, James M. Cleary, Lorin Crawford, Scott R. Manalis, Jonathan A. Nowak, Brian M. Wolpin, William C. Hahn, Andrew J. Aguirre, and Alex K. Shalek. Microenvironment drives cell state, plasticity, and drug response in pancreatic cancer. Cell, 184 (25):6119–6137.e26, Dec 2021. ISSN 0092-8674. doi: 10.1016/j.cell.2021.11.017.

50. Peter van Galen, Volker Hovestadt, Marc H. Wadsworth II, Travis K. Hughes, Gabriel K. Griffin, Sofia Battaglia, Julia A. Verga, Jason Stephansky, Timothy J. Pastika, Jennifer Lombardi Story, Geraldine S. Pinkus, Olga Pozdnyakova, Ilene Galinsky, Richard M. Stone, Timothy A. Graubert, Alex K. Shalek, Jon C. Aster, Andrew A. Lane, and Bradley E. Bernstein. Single-cell rna-seq reveals aml hierarchies relevant to disease progression and immunity. Cell, 176(6):1265–1281.e24, Mar 2019. ISSN 00928674. doi: 10.1016/j.cell.2019.01.031. tex.ids: vangalenSingleCellRNASeqReveals2019a.

51. C. Domínguez Conde, C. Xu, L. B. Jarvis, D. B. Rainbow, S. B. Wells, T. Gomes, S. K. Howlett, O. Suchanek, K. Polanski, H. W. King, L. Mamanova, N. Huang, P. A. Szabo, L. Richardson, L. Bolt, E. S. Fasouli, K. T. Mahbubani, M. Prete, L. Tuck, N. Richoz, Z. K. Tuong, L. Campos, H. S. Mousa, E. J. Needham, S. Pritchard, T. Li, R. Elmentaite, J. Park, E. Rahmani, D. Chen, D. K. Menon, O. A. Bayraktar, L. K. James, K. B. Meyer, N. Yosef, M. R. Clatworthy, P. A. Sims, D. L. Farber, K. Saeb-Parsy, J. L. Jones, and S. A. Teichmann. Cross-tissue immune cell analysis reveals tissue-specific features in humans. Science, 376(6594):eabl5197, May 2022. doi: 10.1126/science.abl5197.
